# Missense variants in health and disease affect distinct functional pathways and proteomics features

**DOI:** 10.1101/512764

**Authors:** Anna Laddach, Joseph Chi-Fung Ng, Franca Fraternali

## Abstract

1

Missense variants are present amongst the healthy population, but some of them are causative of human diseases. Therefore, a classification of variants associated with “healthy” or “diseased” states is not always straightforward. A deeper understanding of the nature of missense variants in health and disease, the cellular processes they may affect, and the general molecular principles which underlie these differences, is essential to better distinguish pathogenic from population variants. Here we quantify variant enrichment across full-length proteins, their domains and 3D-structure defined regions. We integrate this with available transcriptomic and proteomic (protein half-life, thermal stability, abundance) data. Using this approach we have mined a rich set of molecular features which enable us to understand the differences underlying pathogenic and population variants: pathogenic variants mainly affect proteins involved in cell proliferation and nucleotide processing, localise to protein cores and interaction interfaces, and are enriched in more abundant proteins. In terms of their molecular properties, we find that common population variants and pathogenic variants show the greatest contrast. Additionally, in contrary to other studies, we find that rare population variants display features closer to common than pathogenic variants. This study provides molecular details into how different proteins exhibit resilience and/or sensitivity towards missense variants. Such details could be harnessed to predict variant deleteriousness, and prioritise variant-enriched proteins and protein domains for therapeutic targeting and development. The ZoomVar database, which we created for this study, is available at http://fraternalilab.kcl.ac.uk/ZoomVar. It allows users to programmatically annotate a large number of missense variants with protein structural information, and to calculate variant enrichment in different protein structural regions.

**Significance Statement:** One of the greatest challenges in understanding the genetic basis of diseases is to discriminate between likely harmless and potentially disease-causing sequence variants. To better evaluate the pathogenic potential of missense variants, we developed a strategy to quantitatively measure the enrichment of both disease and non disease-related variants within a protein based on its structural and domain organisation. By integrating available transcriptomics and proteomics data, our approach distinguishes pathogenic from population variants far more clearly than previously possible, and reveals hitherto unknown details of how different proteins exhibit resilience and/or sensitivity towards genetic variants. Our results will help to prioritise variant-enriched proteins for therapeutic targeting; we have created the ZoomVar database, accessible at http://fraternalilab.kcl.ac.uk/ZoomVar, for programmatic mapping of user-defined variants to protein structural and domain information.

## 2 Introduction

The genomic revolution has brought about large advances in the identification of disease-associated variants. However, despite the recent explosion of genetic data, the problem of “missing heritability” still persists [1], where the genetic component of a phenotype remains poorly identified. It is often the case that for some variants, a causal link to the disease in question is difficult to establish. Prime examples include variants with low penetrance, and/or those with higher penetrance which are unique to single/few individuals, such as *de novo* variants implicated in developmental disorders [2]. Compensated pathogenic mutations represent another such case, where mutations are present as the wild-type in other species, but their pathogenic effects are negated by another variant [3]. Difficult cases also arise in the analysis of somatic cancer variants, where driver mutations can be challenging to segregate from passenger mutations; moreover this classification may vary from case to case [4]. These variants pose challenges to the detection of disease association using existing statistical methods. In head-to-head comparison against large-scale saturation mutagenesis screens, where mutational impact could be measured *in vitro*, current predictive methods were shown to be limited in accuracy [5, 6]. Moreover, variant impact prediction has focused primarily on detecting differences between disease-associated and common variants, neglecting the distinction between disease-associated and rare variants; thus it has been suggested that these do not perform so well when distinguishing rare neutral variants from those which are pathogenic [7]. The boundary separating disease-causing from neutral variants can be fluid: for example, a number of missense variants thought to lead to severe Mendelian childhood disease were identified in nominally healthy individuals in the ExAC database [8]. Moreover, whether common population variants have more functional impact than rare variants is hotly debated [9, 10]. A better understanding of the molecular principles which underlie differences between disease-associated and population variants is necessary to improve variant classification, and define more clearly the boundaries which distinguish variants in health and disease.

Here we present a detailed analysis of different classes of missense variants, including germline disease variants, somatic cancer variants (both “driver” and “passenger” variants with varying effects on tumour progression), as well as population variants of different frequencies, in the quest to extract the governing principles of variant pathogenicity. We rely on the synergy between utilising two types of data: first, we place emphasis on mapping the localisation of variants on protein structures, taking into account their positions in the protein fold, as well as their proximity to functional sites (e.g. post-translational modifications, or PTMs) [11, 12, 13, 14, 15, 16]. Such protein structural information has been shown to be effective in uncovering the impact of variants at the molecular level [17]. In the field of cancer research, protein structure-based methods have been used to successfully predict cancer driver genes [18, 19], as validated by a recent large-scale study by Bailey and colleagues [20]. Despite such success, 3D structure-based evaluation does not appear to have been applied systematically to other classes of variants (i.e. population and Mendelian disease-associated variants). Second, we also make use of recently available large-scale proteomic measurements, including protein half-life [21], abundance [22], thermal stability [23] and transcriptomics data [24], to uncover biophysical and biochemical principles governing the impact of variants. Our analyses highlight a striking difference in the enrichment of pathogenic and population variants, which depends upon their localisation to protein domain and structural features. Using these features, we demonstrate that rare population variants display characteristics which are more similar to common population variants than to disease-associated variants, reinforcing the boundary between variants in health and disease. This integrative analysis provides molecular details into how resiliance and sensitivity to missense variants are manifested in different proteins and functional pathways. We have created the ZoomVar database (http://fraternalilab.kcl.ac.uk/ZoomVar), which holds the data generated in this analysis. ZoomVar is designed for large-scale programmatic structural annotation of missense variants, and calculation of the enrichment of missense variants in different protein structural regions. Comprehensive mapping of structural localisation of variants could inform the development of therapeutic interventions, e.g. structure-based drug design and/or drug repurposing [25]. More generally, the wealth of features that separates missense variants in health and disease could contribute to building and training next-generation predictors, which hold the promise of improving the accuracy of variant impact prediction.

## 3 Materials and Methods

See Supplementary Methods in SI Appendix for a more detailed account.

### 3.1 Data sources

In this study, we have used variant data from ClinVar (dbSNP BUILD ID 149) [26] (for germline disease variants), COSMIC coding mutations (v80) [27] (for somatic cancer variants) and gnomAD exome data [28] (for population variants). These were mapped against protein sequence data from UniProt [29] and Ensembl [30], protein structural data from the Protein Data Bank (biounit database, downloaded 28/04/2017), and protein interaction data from a large non-redundant protein-protein interaction network (UniPPIN) [31], which incorporates various interaction databases [32, 33, 34, 35, 36] and recent large-scale experimental studies [37, 38, 39]. Protein thermal stability and half-life data were obtained from separate large-scale studies [21, 23]. Transcriptomic data were taken from GTEx [24], while protein abundance data (protein per million [ppm]) for each tissue/sample type were obtained from PaxDb [22].

### 3.2 ZoomVar Database

ZoomVar was constructed by mapping human protein sequences to resolved structures/homologues from the PDB using BLAST [40]. Protein domains were defined by scanning UniProt sequences against the PFAM seed library [41] using HMMER [42]. Per-residue mappings were performed by the alignment softwares T-COFFEE [43] or Stretcher [44] (which was used to map UniProt and Ensembl sequences which were not of the same length, and were too long to align using T-COFFEE). These generated correspondences between PDB structures and those proteins/domains with structural coverage. Interaction complexes were inferred from homologues (defined using BLAST). As an example, if protein *A* and *B* are annotated as interacting in UniPPIN, and their structure homologues *A′* and *B′* are located in a resolved structural complex (and at least one residue from each protein is involved in a shared interface), residues from *A* and *B* are mapped onto *A′* and *B′* to infer their interaction interface. The partner-specific regression formula from HomPPI [45] was used to assign a score and zone to each interaction interface inferred in this way.

### 3.3 Definition of regions

#### Structural regions

We partitioned protein/domain into surface, core and interface regions. Interface regions were considered to be composed of residues which bind to at least one protein interaction partner. The interfaces were assigned using POPSCOMP [46]. Residues with a change in solvent accessible surface area [SASA] *>* 15 Å^2^ were annotated as interface residues [47]. For surface and core regions, these were classified by considering the quotient SASA [Q(SASA)] per residue, which was computed using POPS [48]. Core residues were defined as those with a Q(SASA) *<* 0.15 [47]. Surface residues were defined as those with a Q(SASA) ≥ 0.15 which do not take part in protein-protein interaction interfaces.

#### Order and disorder

Disordered protein regions were predicted using DISOPRED3 [49]. We overlaid these definitions of ordered and disordered regions with Pfam domain boundaries, and partitioned protein sequences into intra-domain ordered, intra-domain disordered and inter-domain disordered regions.

#### Functional sites

Post-translational modification (PTM) sites, specifically ubiquitination and phosphorylation sites, were obtained from PhosphoSitePlus [50]. Regions close to phosphorylation and ubiquitination sites were defined as those within 8Å in Euclidean distance.

#### 3.3.1 Mapping of variant data

Variants in each dataset were annotated according to protein region localisation using the ZoomVar database. Table 1 listed the total number of missense variants in each dataset which have been mapped to each region considered in this study.

**Table 1:** Numbers of missense variants which localise to different levels and regions of protein anatomy. Data are listed for each of gnomAD, COSMIC and ClinVar datasets. Here “common” and “rare” variants are subset of gnomAD defined using the Minor Allele Frequency (MAF) cut-off of 0.01, below which variants are classified as “rare”. This definition is used throughout this work except for the analysis on varying the “rarity” of variants (see Figure 8). Note that the full length domain-type statistics are omitted here, as by definition they will be identical to the “full-length domain” row. Figure 2B illustrates the definition of regions listed here in the first column.

|  | Common | Rare | COSMIC | ClinVar |
| --- | --- | --- | --- | --- |
| full-length protein | 54,571 | 3,806,698 | 1,731,030 | 21,272 |
| full-length domain | 23,634 | 1,772,768 | 852,597 | 15,831 |
| surf | 12,151 | 966,409 | 491,179 | 8,558 |
| interact | 403 | 38,108 | 22,205 | 768 |
| core | 2,789 | 296,291 | 152,356 | 5,194 |
| intra-ord | 20,650 | 1,575,286 | 755,683 | 14,620 |
| intra-dis | 2,984 | 197,482 | 96,914 | 1,211 |
| inter-dis | 17,352 | 1,045,997 | 439,437 | 1,128 |
| phos | 1,661 | 158,192 | 82,364 | 2,362 |
| ubiq | 440 | 52,250 | 25,778 | 607 |

In the exploration of variant enrichment in different structural regions, the COSMIC data was divided into “driver” and “non-driver” subsets, taking drivers as variants which map to all proteins from both tier 1 and tier 2 of the Cancer Gene Census (CGC) (COSMIC v84). The non-driver subset contains all other variants.

### 3.4 Missense variant enrichment across levels of protein anatomy

#### 3.4.1 The protein anatomy

In this study missense variant enrichment was quantified across the “protein anatomy”, in which we partition the human proteome in different ways (see Figure 2A-B). We first define a list of *levels* of the anatomy, namely: (i) individual *proteins*; (ii) specific *domains* of proteins, and; (iii) instances of a *domain-type* across the human proteome. See Results for detailed examples. Here, missense variant enrichment quantification was considered in both of the following scenarios: (i) for a given full-length instance of a *level*, relative to all other instances at the given *level* (e.g. for epidermal growth factor receptor [EGFR] relative to all other proteins in the human proteome), or (ii) for a given *region*, relative to all regions defined under a given criteria at a relevant level (e.g. for the protein core relative to all protein structural regions). For the sake of clarity, in this Methods section the instance of interest is hereafter referred to as the *entity* of interest; its use is clarified in the next paragraph, and illustrated in Figure 1.

**Figure 1:**
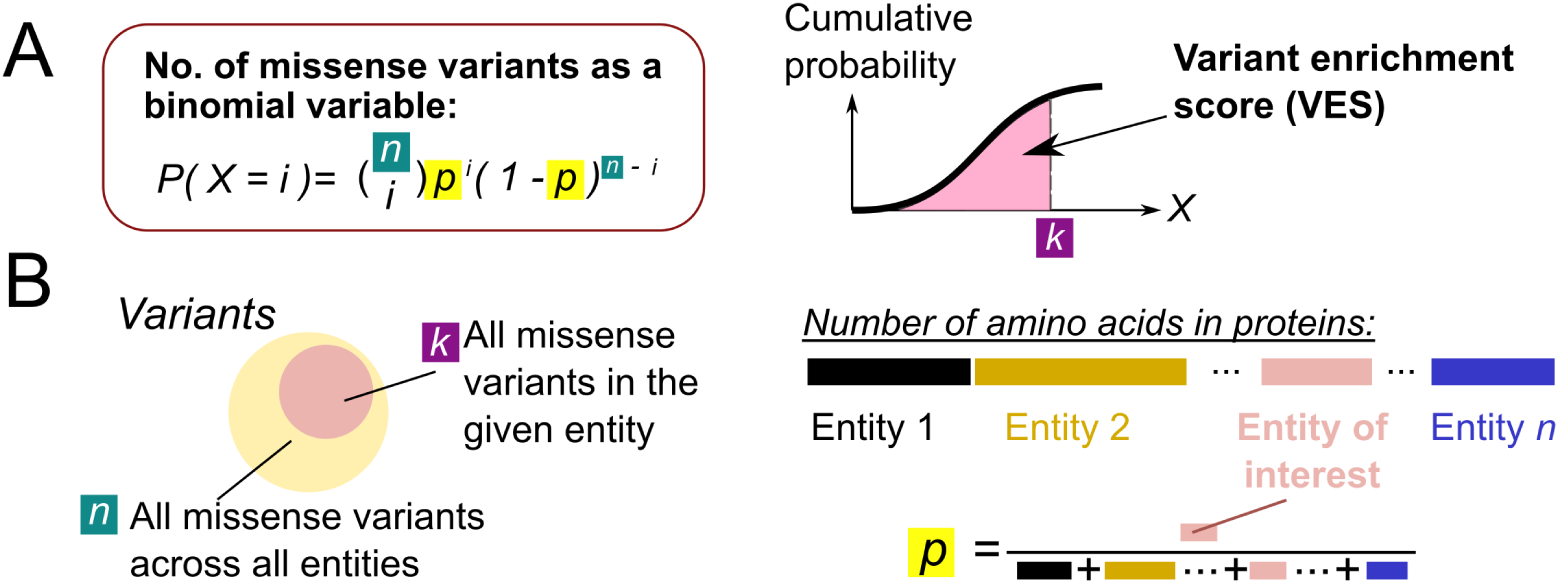
Illustration of missense variant enrichment calculation. (A) The number of missense variants is modelled as a binomial variable. The cumulative distributive function of this binomial variable is taken as a Variant Enrichment Score (VES) for the level examined. (B) Illustration of the choice of parameter in defining the binomial variable used in calculating the VES.

**Figure 2:**
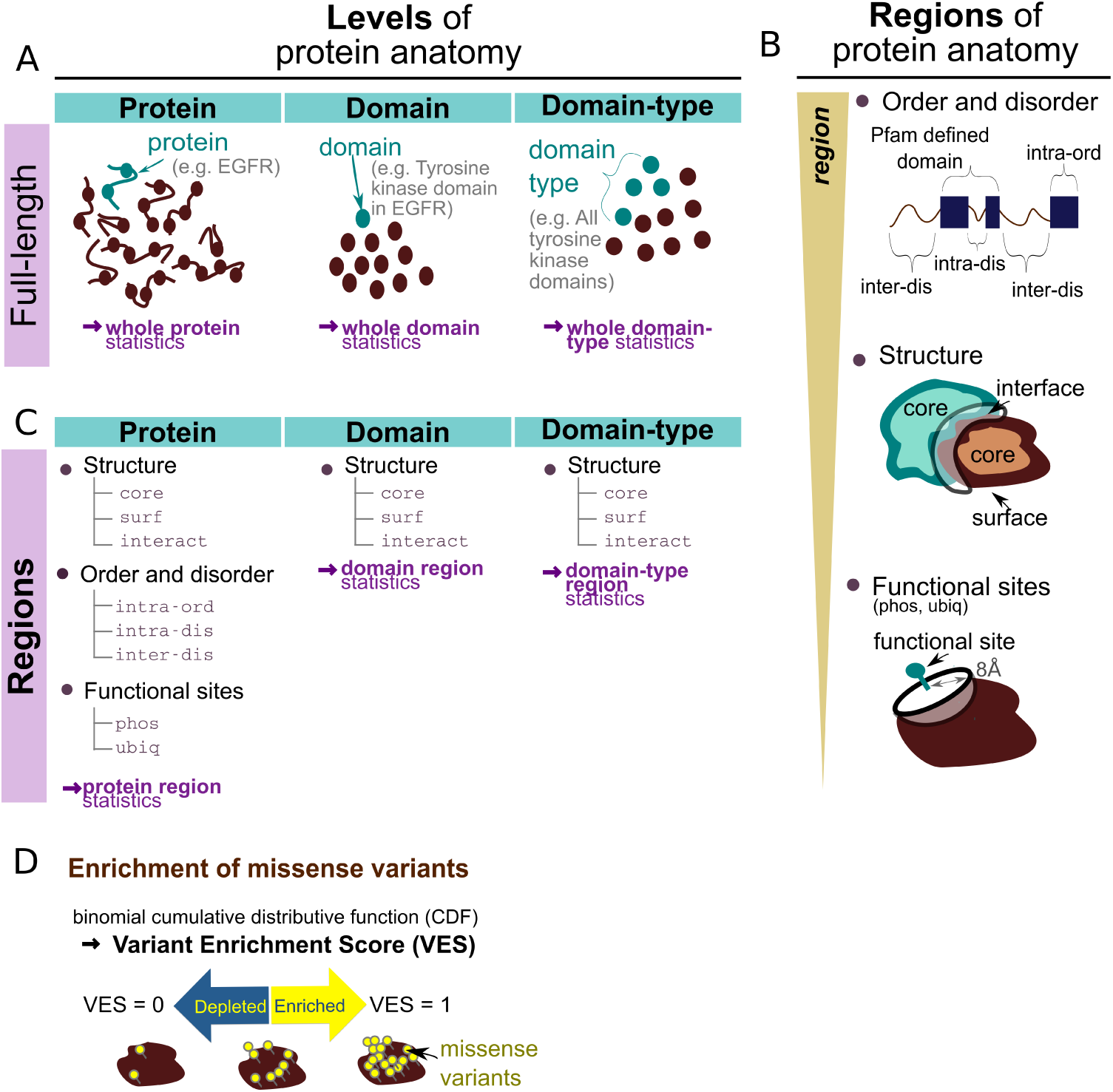
Variant enrichment crossing scales of protein anatomy. (A) Levels of the protein anatomy. At the protein/domain level the number of missense variants in a protein or domain is compared to the number of missense variants in the whole dataset which localise to defined proteins/domains. At the domain-type level the number of missense variants in a particular Pfam defined domain-type, are compared to the total number of missense variants which localise to any Pfam domain-type. These calculations are referred to as the “full-length” protein/domain/domain-type variant enrichment in this manuscript, in contrast with the calculations at regions of protein anatomy defined next. (B) Regions of the protein anatomy. We considered different levels of definition of protein regions, including i) regions close to functional (phosphorylation/ubiquitination) sites; ii) structural regions (core, surface [`surf`] and interface [`interact`]) of a protein, and; iii) regions predicted to be ordered or disordered which lie either within or outside of Pfam-defined domains. (C) Lists of regions considered at each level of the protein anatomy in this study. (D) The calculation of enrichment at the different levels is statistically assessed using the binomial distribution. The binomial cumulative distributive function constitutes a Variant Enrichment Score (VES) with value range 0 to 1, which quantifies enrichment.

#### 3.4.2 Calculation of missense variant enrichment

The binomial cumulative distributive function (Equation 1, also illustrated in Figure 1A) was used to assess the missense variant enrichment of a given *entity*, and the two-tailed binomial test was used to assess the significance of enrichment/depletion. Briefly, the number of missense variants in the given entity is modelled by a Binomial variable *X*, parameterised by: (i) *k*, the number of observed missense variants which localise to the given entity, (ii) *n*, the total number of missense variants which localise to all relevant entities, and (iii) *p*, the ratio of the size of the entity of interest (in terms of number of residues) to the total size of all relevant entities:

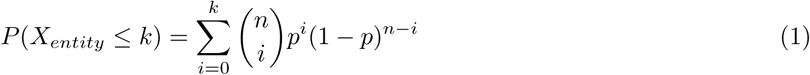

Here, *P* (*X*_*entity*_) ≤ *k*) is the cumulative probability of observing *k* missense variants in the chosen entity. Figure 1B illustrates the definition of parameters in Equation 1, in the calculation of missense variant enrichment. Under this scheme, for example if the missense variant enrichment of a particular protein *W* in dataset *D* is to be calculated, we took

the total number of missense variants in dataset *D* as *n*,

the number of missense variants which localise to *W* as *k*, and

the ratio between the size of protein *W* to the sum of sizes of all proteins in the proteome as *p*.

Hereafter, we refer to the binomial cumulative distributive function as the Variant Enrichment Score (VES). For each analysis at the protein or domain level, the background proteome is defined as all UniProt proteins/domains containing missense variants in any of the datasets analysed. Proteins belonging to immunoglobulin and T cell receptor gene family products were filtered from all analyses (HGNC definition [51]), to avoid the inclusion of variants which could have arisen from the process of affinity maturation. For all calculations of enrichment and simulations involving protein or domain *regions* (e.g. core, surface and interface), cases where the region is of size 0, or where the protein/domain contains no missense variants, were omitted in this analysis. Note that this framework of variant enrichment quantification, in contrary to others [52, 53, 54], is not designed to detect mutational “hotspots” clustered in sequence or structure space. Instead, it quantifies the extent to which missense variants are found in the entity concerned, evaluating whether the number of such variants are more or less than expected.

The overall missense variant enrichment for each dataset was also calculated using a density-based metric *ω* (see Equation 2).

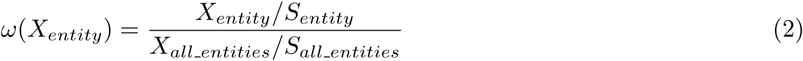

where *S* refers to the size in terms of the number of amino acids.

Here 95 % confidence intervals were estimated via bootstrapping (10,000 iterations). The 2-tailed significance of enrichment/depletion was estimated by simulation of the null background. 10,000 simulations were carried out for each dataset, in which the number of variants which localise to a given entity was kept constant, but their location within the entity randomised. The density of variants was calculated for each simulation and compared to the actual value in order to derive a *p*-value. Simulations were performed in this way, keeping the observed number of missense variants fixed, in order to overcome bias which stems from the assumption that variants are uniformly distributed throughout the proteome.

### 3.5 Enrichment analysis of gene sets

#### 3.5.1 Gene set enrichment

Gene enrichment analyses were performed using Gene Set Enrichment Analysis (GSEA), using the implementation provided by the R fgsea package [55]. Given an enrichment statistic for each query gene, the GSEA algorithm outputs a score per gene set, which quantifies the enrichment of query genes in the sets examined. This is then normalised by the size of the gene set, to give a normalised enrichment score (NES) [56]. We utilise the centred VES enrichment statistic, i.e. substrating 0.5 from the VES, as input into the GSEA algorithm. Thus, proteins with the number of missense variants observed fewer than expected would have a negative score.

#### 3.5.2 Definition of pathway clusters

The pathway normalised enrichment scores (NESs), calculated at the whole protein level for each dataset, were used to perform K-means clustering of KEGG pathways [57]. The R package NbClust [58] was used to determine the optimum number of clusters.

### 3.6 Analysis of expression, abundance, and stability data

Spearman correlations of protein-wise and region missense variant enrichments with expression levels (RPKM), abundance (ppm), half-life (hours), thermal stability (Tm, in °C), and density (mean contacts of core C*α*s) were calculated. Additionally, gene set enrichment analysis was performed as detailed above, except that the mean value for each quantity of interest was subtracted to obtain values centred around 0, allowing both pathway enrichment and depletion to be assessed (see SI Appendix).

### 3.7 Statistics and data visualisation

The majority of data analyses were performed in the R statistical programming environment. All corrections for multiple testing have been done using the Benjamini-Hochberg method in R (p.adjust function). Bootstrapping was performed using the boot package [59]. Spearman correlations were performed using the SpearmanRho function of the DescTools package [60]. Heatmaps were produced with either the heatmap.2 function in the gplots package [61] or the ComplexHeatmap package [62], in which clustering, wherever shown, was performed with hierarchical clustering (hclust function) using default parameters unless otherwise stated. Circos plots were generated with the Circos package [63]. Additionally, binomial cumulative distributive functions were calculated and two-tailed binomial tests performed using the NumPy package in Python [64].

## 4 Results

### 4.1 A detailed protein-centric anatomy of variant enrichment across scales

We present a multifactorial analysis of missense variants observed in the general population (gnomAD database) [28], in comparison to somatic cancer-associated missense variants from the COSMIC database [27] and disease-associated missense variants from the ClinVar database [26]. Throughout this analysis we further divide the gnomAD data by their minor allele frequencies (MAF) into common and rare variants, to investigate whether there are differences between these two subsets. A summary of the numbers of missense variants investigated in each dataset is given in Table 1, and a more detailed breakdown is given in the SI Appendix S4.1.

We compare pathogenic and population variants in terms of their associations with specific features across the molecular scale, in a framework we call “protein anatomy”, in which we partition the human proteome in different ways. This includes the consideration of individual *proteins* (for example, investigating the enrichment of missense variants in the epidermal growth factor receptor [EGFR] protein), specific constituent *domains* of proteins (e.g. the EGFR tyrosine kinase domain), or generally for all instances of a *domain-type* (e.g. all tyrosine kinase domains) found in the human proteome. These are referred to as the *levels* of protein anatomy (Figure 2A). For each level, we analyse variant enrichment in full-length entities (i.e. protein/domain/domain-type, dependent on the level of interest), and also various constituent *regions* defined using different criteria (Figure 2B-C). These include protein structural information (partitioning into surface, core and interacting interfaces), protein disorder predictions (segregating into ordered and disordered regions, within or outside of Pfam domains) and vicinity to functional sites such as phosphorylation and ubiquitination sites. To quantify missense variant enrichment, we employ a similar approach to that used in the prediction of cancer driver genes [65]: variant enrichment has been modelled using a binomial distribution (Methods Equation 1), which allowed for quantification of a Variant Enrichment Score (VES), ranging from 0 to 1, and normalised by the size of the region/protein/domain in question (Figure 2D, also see Methods). We also quantified the robustness of such quantification of variant enrichment: the significance of the enrichment/depletion of missense variants, in terms of their density, is assessed by comparison to simulated null distributions, in which the number of missense variants is kept identical to that observed in the data, but their positions within the protein are randomised. This goes beyond similar studies (e.g. [14, 15, 16]) and addresses biases which could result from the selective focus in molecular studies of disease-related proteins. We use this method to quantify and compare pathogenic and population variants in terms of “microscopic”, atomistic protein structural features, and examined the association between these enrichment statistics and large-scale, “macroscopic” features like functional pathways, as well as the various proteomics features collated (see Methods and below).

### 4.2 Disease-associated and population variants affect different functional pathways

We first investigated whether variants from each dataset localise to proteins which are involved in distinct functional pathways. Gene Set Enrichment Analysis (GSEA) was performed on a pre-ranked list of proteins using their whole-protein VESs computed for each dataset. The pathway enrichment scores were then subjected to clustering and Principal Component Analysis (PCA) (see Materials and Methods). As shown in Figure 3A, variant enrichment segregates pathways into three clusters. Strikingly, each pathway cluster appears to have distinct characteristics (see Figure 3B-D for the pathway terms belonging to each cluster). The cluster visualised in orange is primarily composed of terms associated with cancer, growth and proliferation, whereas that coloured in pink contains pathways associated with splicing, transcription, translation and metabolic terms. Pathways associated with sensory perception and the immune response are found in the “green” cluster. A handful of metabolic pathways also localise to this cluster, however, these appear to be more associated with environmental response and adaptation than those pathways found in the “pink” cluster; for example, pathways associated with the metabolism of drugs and xenobiotics are found here. For brevity, the “orange”, “pink” and “green” clusters will be termed the “proliferation”, “nucleotide processing” and “response” clusters respectively, for the remainder of this text. A list of pathways assigned to each cluster is given in the SI Appendix Section S4.2.

**Figure 3:**
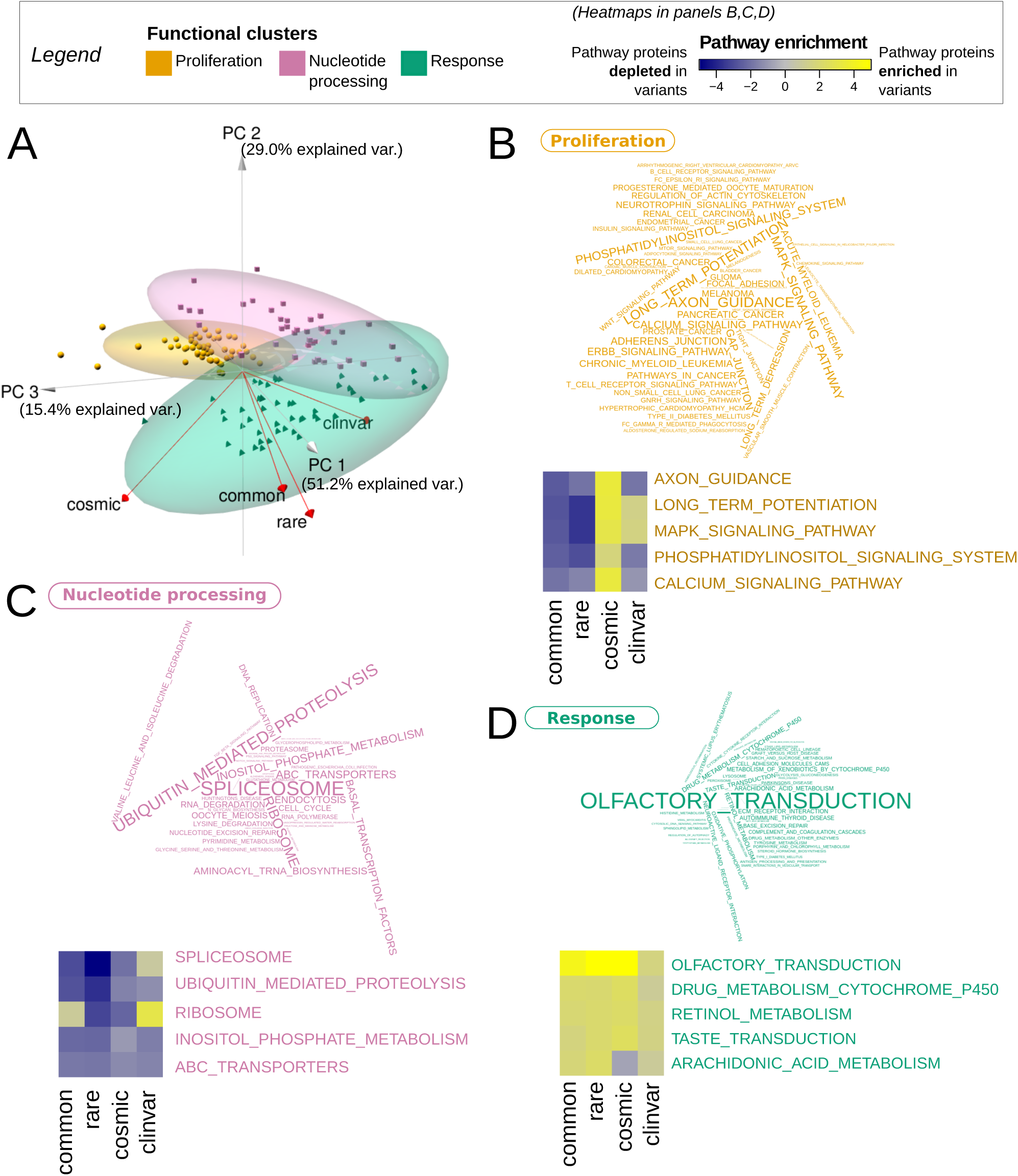
Pathway clusters defined according to protein-wise variant enrichment. (A) At the whole protein level, KEGG pathways form 3 clusters (*k*-means), here visualised as projected onto the first three principal components of the Principal Component Analysis. Pathway enrichment patterns are clearly distinct between COSMIC, ClinVar and gnomAD (rare/common) data, as evidenced by the visualisation of factor loadings (red arrows). (B-D) Pathway terms visualised for the “proliferation” (B), “nucleotide processing” (C) and “response” (D) clusters, and sized by their cluster uniqueness score. The latter is defined as the average of the Euclidian distances to the two other cluster centres. For the top 5 unique pathway terms for each cluster, their pathway enrichment scores calculated with the four variant sets are also visualised in a heatmap.

Strikingly, this visualisation reveals that population variant datasets (gnomAD rare and common) are clearly separated from the disease-associated variant datasets by the first principal component (PC1), whereas COSMIC variants are separated from ClinVar variants along the third principal component (PC3) (Figure 3A). Closer inspection of the pathway enrichment data for the top (most unique) pathways in each cluster reveals a distinction in terms of the functions that different variant datasets implicate (Figure 3B-D). “Response” pathways appear to be enriched in variants in all four datasets, while “proliferation” pathways are consistently enriched in COSMIC variants alongside enrichment in ClinVar variants in a subset of these pathways. Enrichment in ClinVar variants is visible for specific “nucleotide processing” functions, e.g. ribosomes. Across all pathways, the trends of functional distinction can be further visualised in the Circos [63] plot (Figure 4A), whereby the extent of shared enrichment of pathways is visualised by arcs across the different coloured segments. Identical to Figure 3D, pathways in the “response” cluster are enriched in missense variants across all four datasets; enrichment in the “proliferation” cluster is shown only in the disease-associated variants and the “nucleotide processing” cluster is unique to ClinVar variants. This analysis clearly distinguishes missense variants in health from those which are found in diseased individuals; moreover their different tendencies to perturb specific functional pathways are also highlighted.

**Figure 4:**
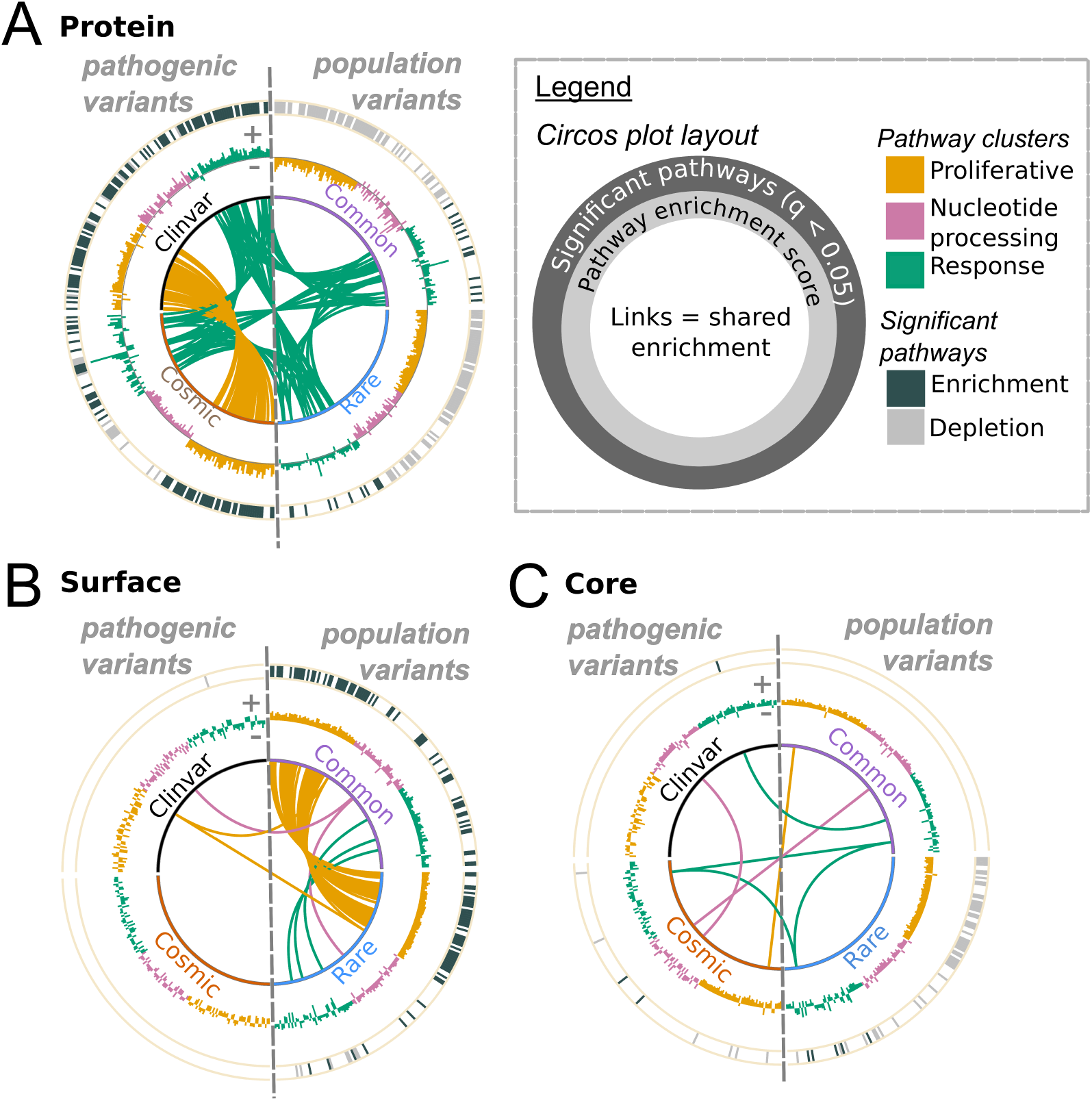
Functional differences of proteins enriched in pathogenic and population missense variants. (A) Enrichment at the whole-protein level for each dataset visualised on a Circos plot (see legend). Pathogenic variants (here referring to ClinVar and COSMIC variants) are depicted on the left, and population variants (gnomAD rare and common) on the right, as indicated on the Circos plot. From the outer to inner layer of the plots, the following are depicted: (i) in the outermost layer of the plot, significant enrichment (dark grey) or depletion (light grey) of a pathway (*q*-value *<* 0.05) is depicted; (ii) in the middle layer of the plot, the normalised pathway enrichment score for each pathway is plotted as a bar graph (the further from the centre, the more positive; see ‘+’ and ‘-’ symbols on the plot); (iii); (iii) in the centre of the plot, links indicate enrichment (*p*-value *<* 0.05) shared between datasets. (B-C) Functional analysis of proteins according to variant enrichment at the protein surface and core, visualised on a Circos plot. Data are visualised in the same format as in panel (A).

We went on to extend this analysis to across protein structural regions (Figures 4 and SI Appendix Figure S4). Here we find that proteins enriched in gnomAD variants at the surface (Figure 4B) are significantly enriched in pathways belonging to the “proliferation” cluster. Moreover, this enrichment is shared between common and rare variants (albeit not significant for common variants in individual pathways after false discovery rate [FDR] correction). Proteins with surfaces enriched in disease-associated variants (from COSMIC and ClinVar) are, contrastingly, not enriched in “proliferation” cluster pathways. Pathways in the “proliferation” cluster show either depletion (for rare variants) or no patterns at all (for common variants) for population variants, when the protein core (Figure 4C) and interface (SI Appendix Figure S4B) are concerned. This could indicate that population variants avoid the core of proliferation-related proteins. Interestingly, the “nucleotide processing” cluster does not show such a marked enrichment of variants which localise to the surface in the gnomAD dataset, a possible indication that proteins in these pathways are more robust to disruption of their structural fold, compared to those in the “proliferation” cluster. These data show that there is clearly an interplay between variant localisation at macroscopic (functional pathways) and microscopic (structural regions) protein features.

### 4.3 Population and disease-associated variants localise to different protein regions

We then zoomed in to view trends in the enrichment of variants in different regions of the protein anatomy, defined using structural information, order and disorder, and the vicinity (distance ≤ 8 Å) of the variant positions to post-translational modifications (see above). The following findings are highlighted:

#### Different structural localisation of pathogenic *vs* population missense variants

In agreement with previous research [13, 14, 15, 16], we find ClinVar variants to be enriched in both protein cores and interfaces, but depleted on protein surfaces (Figure 5A-C and SI Appendix Figure S4). This reflects the potential disruption, caused by such mutations, of structurally and functionally important sites. The enrichment of ClinVar variants in structurally important sites is further demonstrated by their tendency to affect residues which are highly connected when considering network representations of protein structures (SI Appendix Section S2.1). GnomAD variants (both common and rare) and somatic non-driver variants display the opposite trend, as variants tend to localise preferentially to protein surfaces, and are therefore less likely to impact on protein structure and function than either core or interface mutations. Somatic driver variants follow trends closer to ClinVar variants, with slight, but significant, depletion on the surface, but enrichment in the core. Protein interfaces are enriched in disease-associated variants but depleted of gnomAD rare variants. GnomAD common variants appear neither significantly enriched nor depleted, however this may result from the relative sparsity and high dispersion of the data (fewer variants are shared between many individuals; see Table 1). Interestingly, COSMIC non-driver variants appear depleted in interacting interfaces. However, it becomes clear that they are actually significantly enriched when compared to simulated null distributions (see SI Appendix Figure S5), and that this enrichment is due to a small subset of proteins which harbour a large number of variants at interface regions. Genes in which these variants reside may be putative driver genes (see SI Appendix Section S4.3), as a number of known driver genes are enriched in variants in protein interface regions [14, 20, 65], and this phenomenon has been exploited by by Porta-Pardo and colleagues [65] to identify cancer driver genes.

**Figure 5:**
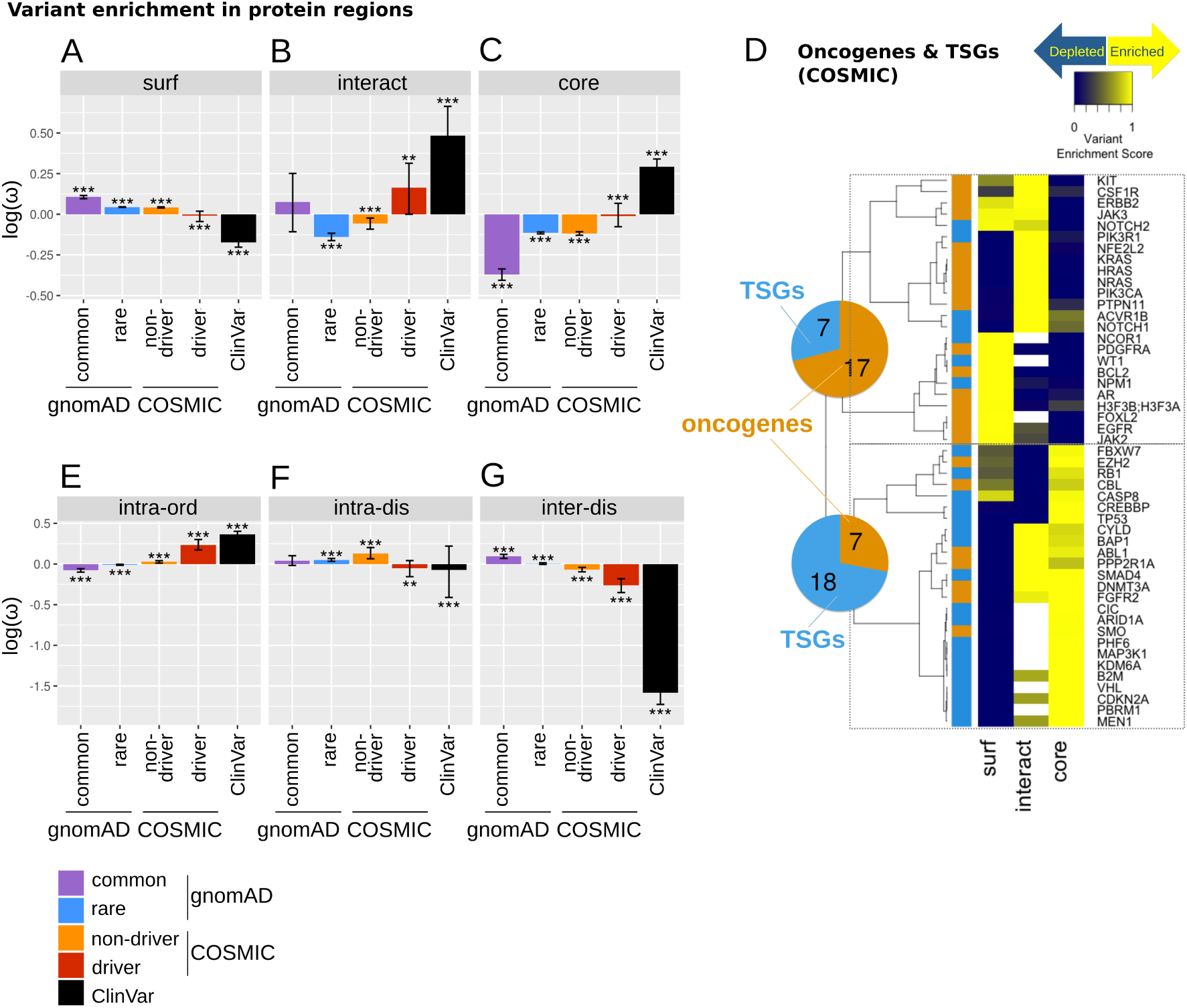
The localisation of variants to protein regions. (A-C) The density of mutations in different protein regions, calculated using Equation 1. Density values (*ω*) were log-transformed such that negative values indicate a depletion of missense variants, while positive values indicate enrichment. Error bars depict 95% confidence intervals obtained from bootstrapping. Significance was calculated by comparison to simulated missense variant distributions (significance level indicated by: * *q*-value *<* 0.05, ** *q*-value *<* 0.001, *** *q*-value *<* 0.0001). Note here the COSMIC set is split into driver (i.e. mutations mapping to proteins found in the COSMIC Cancer Gene Census [CGC]) and non-driver subsets. Data are shown for protein surface (`surf`, panel A), interacting interface (`interact`, B) and core (C). (D) Enrichment of COSMIC missense variants in protein core, surface and interface regions, across a list of annotated oncogene (orange annotations next to the dendrogram) and tumour suppressor gene (TSG, blue) products. The genes were grouped into two clusters using hierarchical clustering (see dendrogram by rows), with the pie charts enumerating the number of oncogenes and TSGs in each cluster. (E-G) Density of mutations analogous to panels (A-C) but in regions defined by order and disorder. Data are shown here for intra-domain ordered (`intra-ord`, panel E), intra-domain disordered (`intra-dis`, F) and inter-domain disorered (`inter-dis`, G)

#### Protein structural information distinguishes oncogenes and TSGs

We analysed, in greater granularity, missense variant enrichment in the COSMIC dataset. By examining a curated list of oncogenes and tumour suppressor genes (TSGs) [66], we found, in agreement with others [13, 14, 67], that these proteins could be classified into two groups by considering their patterns of variant distribution, one comprising proteins enriched in mutations mainly at interaction interfaces and surfaces, and another group in the core (Figure 5D). Some proteins in the latter group also show enrichment in interacting interfaces, but a clear depletion of mutations at the surface is evident. The segregation of these two groups in terms of cancer driver status has strong statistical support (Fisher-exact test *p*-value = 0.0042): the first group of proteins is mainly (17 out of 24) composed of oncogenes, and the other mainly of TSGs (17 out of 25). These results are consistent with the hypotheses that activating mutations in oncogenes are likely to affect particular functions by perturbing specific interactions, whilst inactivating mutations in TSGs abrogate protein function [14, 67]. Taking mutations in oncogenes and TSGs as two separate groups, the GSEA results confirm a similar trend of structural localisation (SI Appendix Figure S6); moreover, it can also be seen that the disease-associated datasets (ClinVar and COSMIC) show opposite patterns of enrichment in comparison to the gnomAD data (SI Appendix Figure S6) [13, 14, 67]. These results go beyond similar studies, and show that by analysing the structural localisation of variants, cancer driver genes can be separated into the distinct groups of oncogenes and TSGs.

#### Pathogenic variants tend to localise to ordered regions within domains

For variant enrichment in ordered and disordered regions, we again observe clear segregation between disease and population variants (Figure 5E-G). ClinVar and COSMIC variants are depleted in inter-domain disordered regions and enriched in intra-domain ordered regions. In contrast, gnomAD variants (both rare and common) appear enriched in inter-domain disordered regions and depleted in intra-domain ordered regions. These results suggest, as one would intuitively expect, that variants are more likely to be pathogenic if they fall within ordered domain regions.

#### Pathogenic variants are close to phosphorylation sites

When proximity to PTMs is considered (SI Appendix Figure S5), ClinVar variants appear enriched in terms of the density of missense variants close to phosphorylation sites, but not significantly so in comparison to the simulated null background; this may be again due to data sparsity, as suggested by large bootstrapped confidence intervals (SI Appendix Figure S5). COSMIC driver variants are also close to phosphorylation sites; however, COSMIC non-driver variants, which appear depleted in terms of variant density, are also significantly enriched close to phosphorylation sites in comparison to simulated null distributions (SI Appendix Figure S5). This indicates that, in agreement with a number of other studies [68, 69], the disruption of phosphorylation sites may play a particularly important role in cancer. In contrast to phosphorylation sites, all datasets appear depleted of variants close to ubiquitination sites (SI Appendix Figure S5).

These analyses conclude that the enrichment of missense variants at various structural features consistently segregate population variants from disease-associated ones. For the majority of structural regions defined here, the greatest, most consistent distinction is always seen between common and ClinVar variants, provided that the data are not too sparse.

### 4.4 Towards a domain-centric landscape of variant enrichment

We now proceed from the protein level to examine variant enrichment across domain-types. Agglomerating missense variants at the domain-type level has the advantage of enhancing the statistical power to detect variant enrichment in terms of different protein structural features ([53] and references therein). Here, in contrast to previous studies which focus on variants clustered in sequence or structure space [52, 53, 54], we present an unbiased landscape of variant enrichment across domain-types, and compare the patterns of enrichment of variants from the different health and disease datasets which we have examined above. We focus our discussion on the most variant-enriched domain-types from each of the four variant sets. A comprehensive list comprising the union of the top 20 enriched domain-types for each dataset can be found in SI Appendix Figure S7. Figure 6 shows some selected examples of this list; the missense variant enrichment at the full-length domain-type level (Figure 6A) and in each structural regions (Figure 6B) are depicted. Here domain-types which are enriched in variants only in the COSMIC and ClinVar datasets can be seen, including known drug targets such as tyrosine kinase (Pkinase Tyr) and ion channel (Ion trans) domain-types. A handful of domain-types, which are only enriched in COSMIC variants, include Cadherin tail and Laminin G 2 (Figure 6A), both of which play an important role in cancer [70, 71]. Some domain-types (e.g. Serpin, UDPGT, Collagen and EGF CA) contain variants from all four datasets. In such domains, it is likely that the precise structural localisation of a variant determines whether it plays a pathogenic role. Intriguingly a few domain-types, such as NPIP (Nuclear pore complex interacting protein) and NUT (Nuclear Testis protein) appear only enriched in common variants (Figure 6A). This could suggest that these domains take part in functions for which it is desirable to maintain diversity within a population; however, little is known about either domain-type [72, 73]. Thereby this further highlights the bias in the number of studies targeting domains associated with disease, rather than those enriched in population variants.

**Figure 6:**
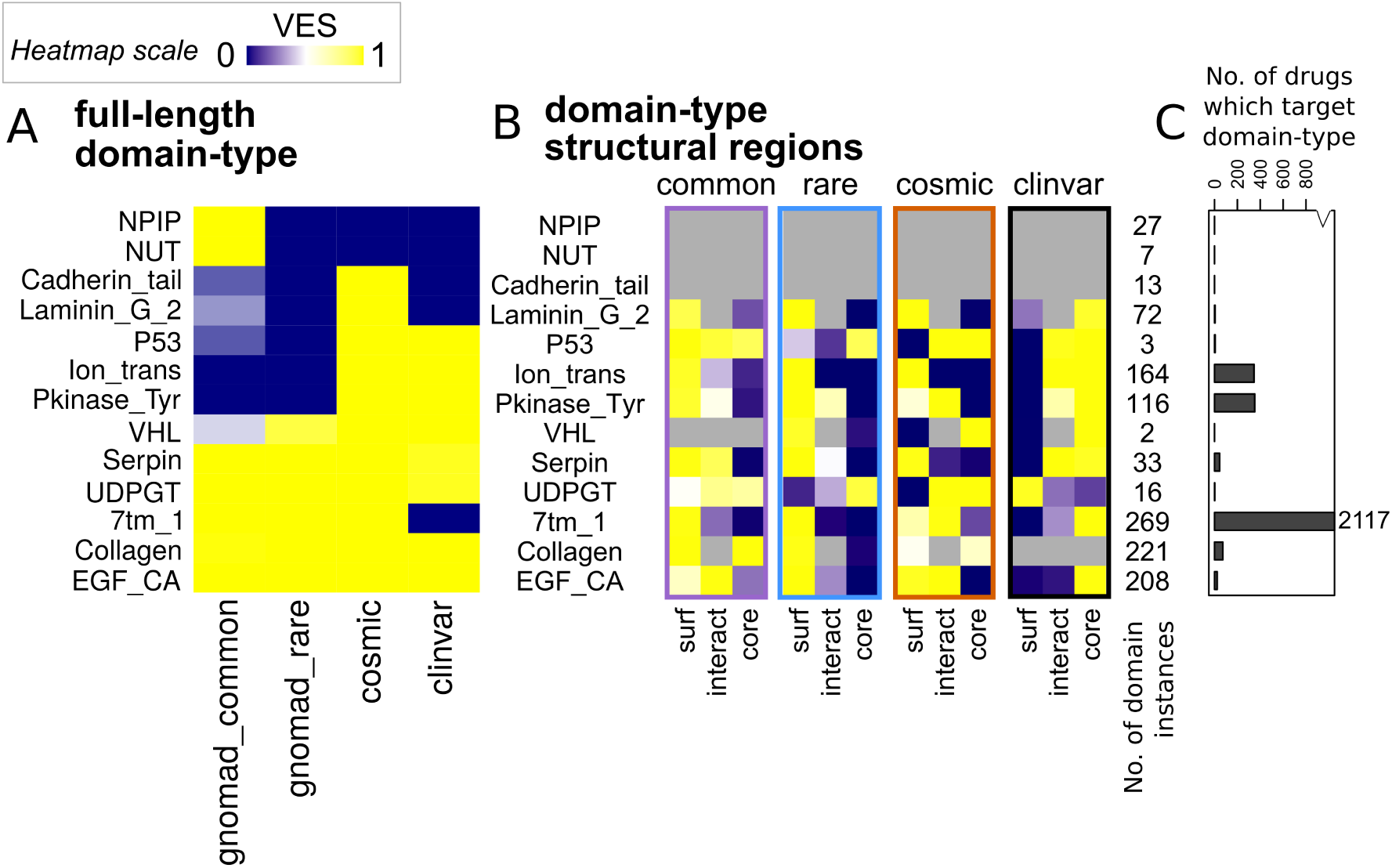
A domain-centric landscape of variant enrichment. Here selected domain-types discussed in the main text are depicted. See SI Appendix Figure S7 for the data on a more comprehensive list of domain-types. (A) VES at the full-length domain-type level. (B) VES calculated for each structural region (surface [`surf`], interacting interface [`interact`] and core) for the selected domain-types. (C) The number of drugs known to target proteins containing each domain-type is depicted as a bar graph. Note the cut numeric axis; the number of drugs which target the only outlier, the 7tm 1 (GPCR) domain, is noted on the plot.

It also becomes apparent that the global trends in variant localisation to the core, surface and interface regions observed above are recapitulated here (Figure 6B) for those domain-types with structural coverage. The majority of domains are enriched in gnomAD (rare and common) variants at the surface but ClinVar variants at the core. For COSMIC the patterns of localisation are more mixed, but it is clear that in comparison to the gnomAD sets, a larger proportion of domain-types are enriched in COSMIC variants at the core or interface. These include domain-types with known cancer driver associations, such as the P53 and VHL domains [74]. Case studies on CATH architectures [75] and DNA-binding domains further highlight our observed patterns of variant enrichment (see SI Appendix Sections S2.3 and S2.2).

We also explored how the targeting of domains by drugs and small molecules mapped to the observed landscape of variant enrichment. Using DrugBank [76] data we observe that the targeting of domain-types by existing drugs is highly biased towards a small number of domain-types (Figure 6C), such as GPCRs and tyrosine kinase, as already extensively pointed out [77]. Indeed, we observe a large number of drugs targeting proteins containing 7tm (GPCR) domains. These domains are enriched in variants from the gnomAD and COSMIC database, but are devoid of disease-associated ClinVar variants (Figure 6A). By analysing drug availability together with variant enrichment, this approach allows for more informed decisions in selecting new therapeutic targets. For example, there are domain-types which could be targeted by few or no drugs, but are enriched in COSMIC and/or ClinVar variants. This could offer a starting point to prioritise drug discovery efforts for these domain-types. For domain-types already targetable by drugs, our analysis highlights domains to which multiple disease-associated variants localise, which could give scope for drug repurposing or redesign (see Discussion).

### 4.5 Proteomics and transcriptomics features associate with variant localisation

Proteins, of course, do not function in isolation but in the crowded environment of the cell [78]. Therefore, the properties of proteins in cells, including their quantities, turnover rates and thermal stability, can crucially affect the fitness of a protein to perform its function. Here we ask if variant enrichments are associated with these proteomics features. We have made use of large-scale proteomics data, including protein abundance data for various organs from PaxDb [22], proteomics surveys of protein half-lives and thermal stability [23, 21], together with transcriptomics data (GTEx database [24]), to explore relationships between these features and variant localisation.

We first compared the thermal stability and abundance of proteins enriched in each class of variants. This comparison demonstrates that for proteins affected by ClinVar variants, their wild-types tend to be more stable and abundant in comparison to those proteins enriched with gnomAD variants (SI Appendix Figure S8). Extending to the entire proteome, the protein-wise Variant Enrichment Scores of disease-associated variants displays positive correlations with protein abundance, expression, half-life and thermal stability, whereas population variants exhibit the opposite trend (Figure 7 and SI Appendix Figures S10-S11). However, zooming into the enrichment of variants in the core of protein structures, we found that in comparison to all regions of proteins with resolved structure, proteins more enriched in ClinVar variants in the core tend to be less abundant and less stable, whereas the contrary is true for rare population variants (Figure 7). Thus our results indicate two competing trends for disease-associated variants: (i) disease-associated variants tend to localise to more abundant and stable proteins, which may suggest that these proteins are more sensitive to perturbation by variants; (ii) disease-associated variants in protein cores tend to localise to less stable proteins, which is consistent with the idea that such proteins might be more easily destabilised to a degree at which function is deleteriously impacted (see Discussion). gnomAD common data also show negative correlations with protein stability, for variants occurring at the core; this could potentially support the argument presented by Mahlich and colleagues [9] that common variants could affect molecular function more than rare variants. However, we believe this is more likely to be due to the fact that very few common variants localise to protein cores, as shown by Figure 5B, resulting in sparse statistics (i.e. the correlation is calculated over Variant Enrichment Scores which are already very low). Analogous correlations for variant enrichments at protein surfaces display opposite trends to those observed at the protein cores (see SI Appendix Figure S9). Due to the relative sparsity of variants which map to protein interfaces, we believe it is difficult to draw robust conclusions from any trends observed for correlations of proteomics data with variant enrichment at protein-protein interaction sites.

**Figure 7:**
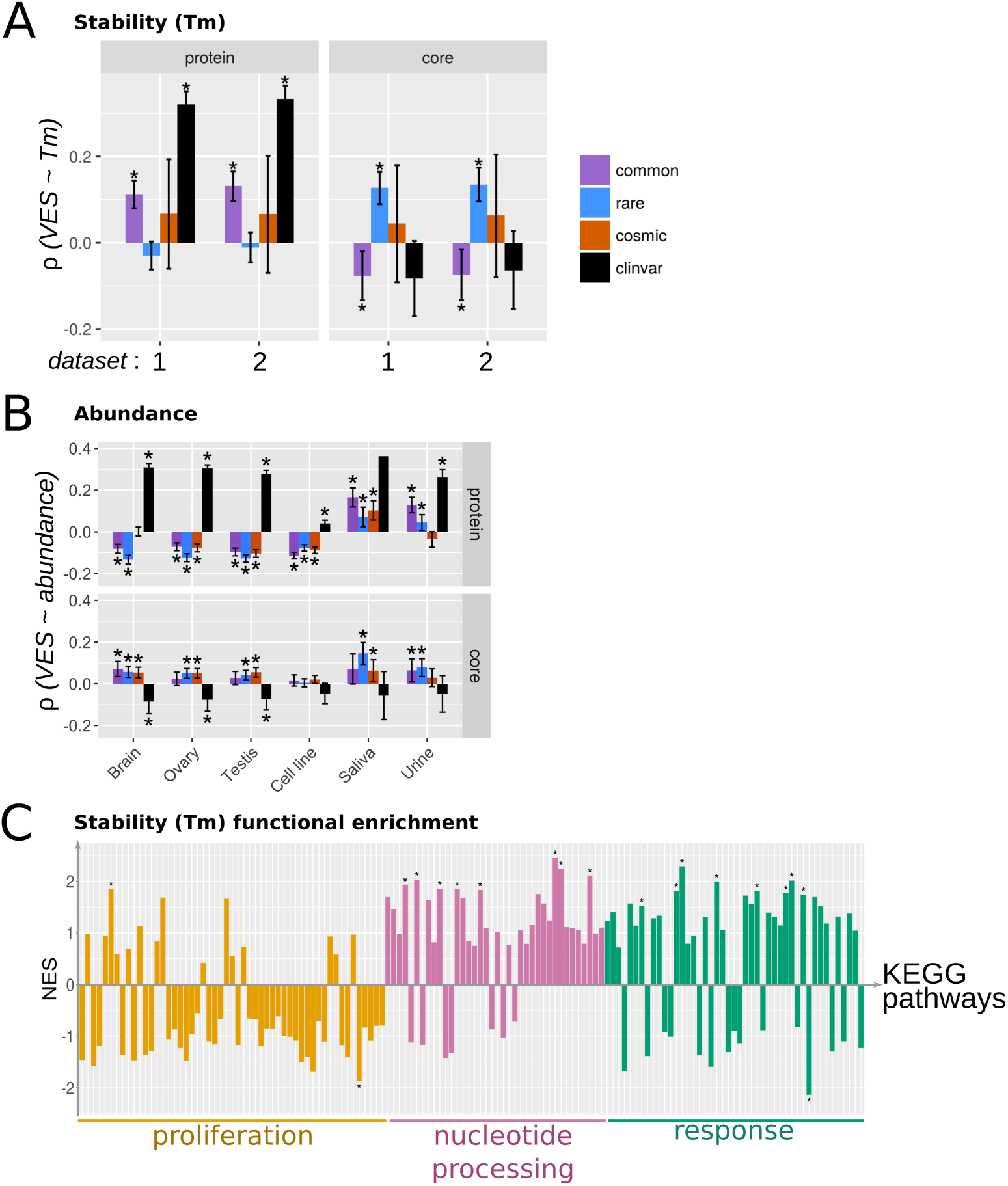
The protein-wise enrichment of missense variants in comparison to protein abundance, expression and stability. Spearman correlations for missense variant enrichment (quantified as VESs) at the full-length protein and the core with (A) protein stability in terms of melting temperature (Tm, °C) and (B) protein abundance (ppm) are depicted here. For (A), the Tm data was taken from [21], in which two measurements of Tm in the absence of any drug treatment were available; both measurements are considered, and are denoted datasets “1” and “2” in the plot. For (B), only data from selected tissue types are listed. See SI Appendix Figure S9 for the complete list. (C) Functional enrichment of proteins in KEGG pathways according to Tm. The Normalised Enrichment Score (NES) is shown on the vertical axis. KEGG Pathways are listed on the horizontal axis, and grouped to the 3 clusters as defined in Figure 3. See SI Appendix Figure S13 for a complete list of pathways depicted here.

One might expect that mutations would be less easily accommodated in cores of densely packed proteins, i.e. those likely to have higher thermal stability. To assess this we calculate the mean number of C*α* contacts within 8 Å of core residues, as a proxy for protein density. We observe weak but significant correlation between this metric and protein thermal stability (Tm measurements from two replicates reported by Franken and colleagues [21]: replicate 1 [*ρ* = 0.168, *q* = 1.464e-12] and replicate 2 [*ρ* = 0.185, *q* = 1.529e-13]). This metric of core density (see SI Appendix Section S1.4 for details) is negatively correlated with the core Variant Enrichment Score for the gnomAD common dataset. No other datasets show significant correlations with core density, however a clear trend emerges in which correlations with the core density become progressively more positive in the order of gnomAD common, gnomAD rare, COSMIC driver, COSMIC non-driver and ClinVar (see SI Appendix Figure S12). This suggests variants may be more disruptive if they localise to a densely packed protein core.

We highlight two more observations in terms of the interplay between proteomics features. First, the various proteomics features examined here are inter-dependent. Protein abundance and thermal stability are significantly correlated with one another (see SI Appendix Section S4.6), in agreement with the work of Leuenberger and colleagues [79]. Moreover, core packing and thermal stability are correlated, albeit with a low correlation coefficient. The correlation values displayed in Figure 7 are also typically of a weak effect. Therefore, the interplay between variant enrichment and proteomics features appear multifaceted and complex. Secondly, in the analysis of protein abundance, the trends observed with variant enrichment at both full-length proteins and specifically the protein core, are less pronounced for cell line data and break down for extracellular fluids (saliva and urine, Figure 7B). The correlation is most evident for tissues containing long-lived cell-types, such as the brain, ovary and testis. Transcriptomics data (SI Appendix Figure S10) again reinforces this picture, albeit with less contrast between datasets (particularly at the protein core). This brings finer granularity into assessing the impact of variants in different organs and contexts.

We finally ask whether correlations with these proteomic and transcriptomic features could be associated with the specific functional roles of the involved proteins. For the majority of proteomic and transcriptomic features, no clear associations with the functional clusters identified in Figure 3 can be detected (see SI Appendix Figures S11-S13). An exception to this is protein thermal stability: pathways which belong to the “proliferation” cluster are clearly enriched in proteins of lower stability than the other two clusters (Figure 7C). This suggests that proliferation-related proteins may be vulnerable to disruption by mutations which localise to their already unstable cores. Moreover, this agrees with the idea proposed above (Figure 4), that “proliferation” cluster proteins may be less robust to disruption. Taken together, these analyses provide fine molecular details into defining both the resilience towards variants, and the sensitivity towards variants, for a given protein (see Discussion). Moreover, the association of variant enrichment with features such as abundance and stability is indicative of the condition (disease/health) associated with the variants.

### 4.6 Rare variants are similar to common variants

Throughout the majority of analyses, the greatest segregation of data can be seen between common and disease-associated variants (Figures 3–5). Here we vary the criteria with which to define rarity of variants in the gnomAD set, to examine whether extremely rare variants would show characteristics akin to disease-associated variants. Figure 8 demonstrates that rare variants are more similar to common variants, both in terms of the functional pathways they affect, and in terms of the protein regions they localise to (core, surface and interface, order and disorder). If more stringent minor allele frequency (MAF) thresholds are used to define rare variants, their properties move towards those of disease-associated variants, but still remain closest to those of common variants (Figure 8 and SI Appendix Figure S14). A visible separation between common and rare variants, especially in the pathway analysis, can only be seen if an extreme MAF cutoff (*<*0.00001) is used. This reinforces the boundary between population and disease-associated variants, and supports the distinction in terms of molecular characteristics associated with rare population variants and disease-associated variants.

**Figure 8:**
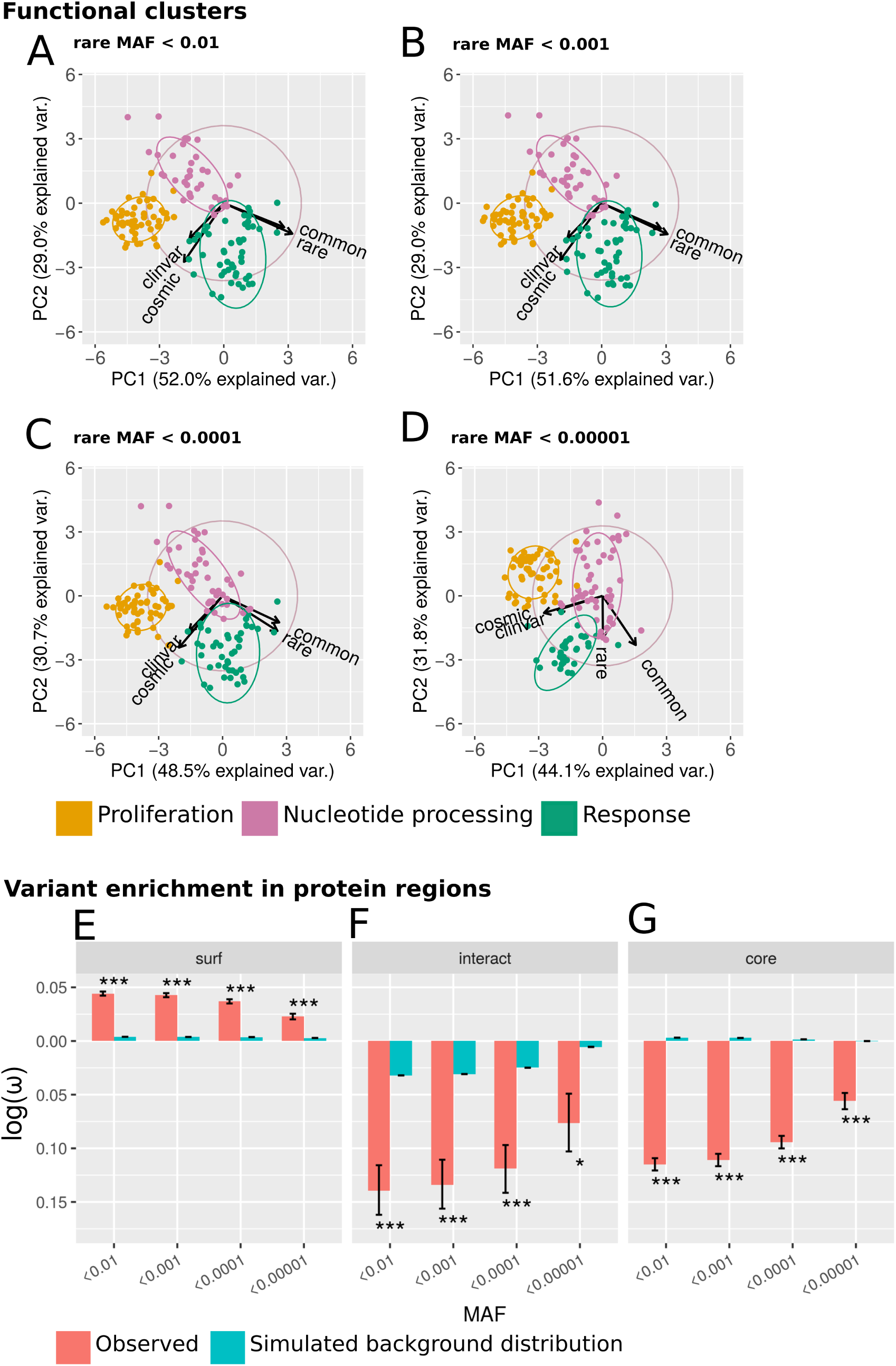
Rare variants are similar to common variants. (A-D) As for Figure 3A, but only the first 2 PCs are depicted, and, in separated panels, increasingly stringent minor allele frequencies (MAFs) used to define rare variants. MAF cutoffs of 0.01 (panel A, data identical to Figure 3A), 0.001 (B), 0.0001 (C) and 0.00001 (D) are considered here. (E-G) The localisation of rare variants to protein surface (E), interface (F) and core (G). Rare variants have been defined using different MAF cut-offs as shown on the x-axes. Here the density metrics (*ω*) were log-transformed such that negative values indicate a depletion of missense variants, while positive values indicate enrichment. Results for the observed variants (red bars), as well as the background levels based on simulated null distributions (cyan bars) are shown.

## 5 Discussion and conclusions

Variants found in diseased and healthy populations are distributed across the proteome, each exerting a varying impact on molecular function. A detailed analysis of the patterns of variant localisation could help in understanding the functional constraints that different parts of the genome experience, and improve the interpretation of variant impact. Throughout this work, we show that missense variants in the general population, considered nominally healthy, show properties distinct from those in disease cohorts, from both macroscopic (“omics” features and functional pathways) and microscopic (protein structural localisation) perspectives. Additionally, we find that the properties of rare variants remain close to those of common variants. These findings contrast with other observations [9], which suggest that common variants have more impact on molecular function than rare variants. In this study, only for a few proteomics properties, such as the thermal stability and abundance of the affected proteins, common variants appear closer in character to disease-associated variants than to rare variants. However, for these few properties, the results might not be robust due to the sparsity of the data. Rare genetic variations are abundant across individuals [80, 81], with some predicted to confer a regulatory impact [82] or loss of function [83]. Alhuzimi and colleagues [10] suggest that the properties of genes enriched in rare population variants are similar to those enriched in disease-associated variants, and are thus good candidates for harbouring unknown disease associations. Instead, we show that proteins enriched in rare variants are, based on the associated functional pathways, most similar to those enriched in common variants (Figure 8). Moreover, our results show that population variants implicate functions mainly associated with environmental response (Figure 3), in agreement with results from evolutionary studies reviewed in [84].

We have dissected the extent of variant enrichment in diverse datasets and across different protein regions (Figure 2). Whereas protein structural information has been utilised to annotate genetic variants and prioritise impactful variants for further investigations, many of the published methods focus on 3D-structural “hotspots”, prioritising variants which cluster together in three-dimensional space (e.g. in [52, 53, 54]). Here we have adopted an alternative approach, and quantified enrichment of missense variants without the pre-condition of spatial clustering. This provides an unbiased resource to map missense variants to protein structural data. The calculation of variant enrichment, as an additional layer of annotation, provides a unique link between cataloguing sequence variants and understanding both their mechanistic and functional effects. This supplies invaluable information to researchers studying specific proteins or domains, or focusing on proteins involved in a particular function (e.g. DNA binding; SI Appendix Figure S3). By analysing the enrichment of variants in protein regions (core, surface, interface, disorder and order, PTM vicinity), we recapitulate trends observed by previous studies (e.g. in the comparison of oncogenes and TSGs; Figure 5D) [14, 16, 15, 13, 67], but also shed light on the debate as to whether somatic cancer variants are enriched in interface regions, by simulating null distributions of variants. These simulations show that it is essential to consider that variants from different datasets are not uniformly and randomly distributed throughout the proteome. Through density-based metrics we find that somatic cancer variants appear at first sight not enriched in protein interfaces, however by comparison to a simulated null background we do find an enrichment (SI Appendix Figure S5). A similar simulation-based approach was taken by Gress and colleagues [13], but they found no significant enrichment for COSMIC variants in interface regions. Whilst they analysed a filtered set of mutations likely to play a driver role, we investigated all somatic variants and addressed separately mutations that localise to defined driver and non-driver genes. Throughout this analysis, we have of course been limited by the number of proteins with available structural data, despite enrichment with homologous structures. We are also still limited by the structural coverage of protein interactions; although enough data exists to uncover broad trends, our analyses of protein-protein interaction sites generally lacked statistical power. Moreover, it is likely that a more detailed picture will emerge if different classes of protein interactions (e.g. transient *vs* permanent interactions) could be probed systematically. We envisage that the recent advances in cryo-electron microscopy [85], and the integration of structural data derived by a variety of techniques [86], will further increase the structural coverage of the protein-protein interaction network, enabling such finer-grained analyses in the future.

Our analysis of probing the associations between missense variant enrichment and proteomic features, is, to the best of our knowledge, unprecedented, and has only been made possible due to the recent release of large-scale proteomics data [21, 22, 23, 79]. We observe correlations which suggest an interplay between variant enrichment, protein abundance and thermal stability (Figure 8). These associations are indicative of the relative resilience (tolerance) and sensitivity of proteins towards missense variants. Our results thus illustrate a set of features, based on which different parts of the proteome could be assessed for their tendency to be enriched in disease-associated or population variants (Figure 9). Population variants tend to be enriched on protein surfaces but depleted in core and interacting sites, and tend to be found in less abundant, less stable proteins. These features could potentially contribute to limit the functional impact of missense variants found in the general population. On the other hand, disease-associated variants localise preferentially to proteins which are highly expressed and abundant (Figure 9). However, when selectively looking at variants mapping to protein cores, which presumably could bring about the most dramatic impact on fitness, disease-associated variants are actually associated with cores of less stable, less abundant proteins (Figure 7). Such proteins are conceivably easier to inflict damage at the core (although it is also possible that some of these proteins are not globular, but are instead more extended in conformation, see below). The combination of these molecular features could also suggest the likely selection pressure a protein could experience under different contexts. For example, certain proteins, possibly further to the left of the spectrum presented in Figure 9, could show a more extreme combination of molecular features compared to those proteins discussed here to be enriched in disease-associated variants. These proteins are likely to be highly sensitive towards variants, such that any of such variants would be lethal (Figure 9) and be eliminated via selection; these lethal variants suffer from under-sampling in the data analysed here. On the other hand, variants, if localised to sensitive proteins, may bring benefits to cell viability; these variants could ascertain a role in driving cancers (Figure 5).

**Figure 9:**
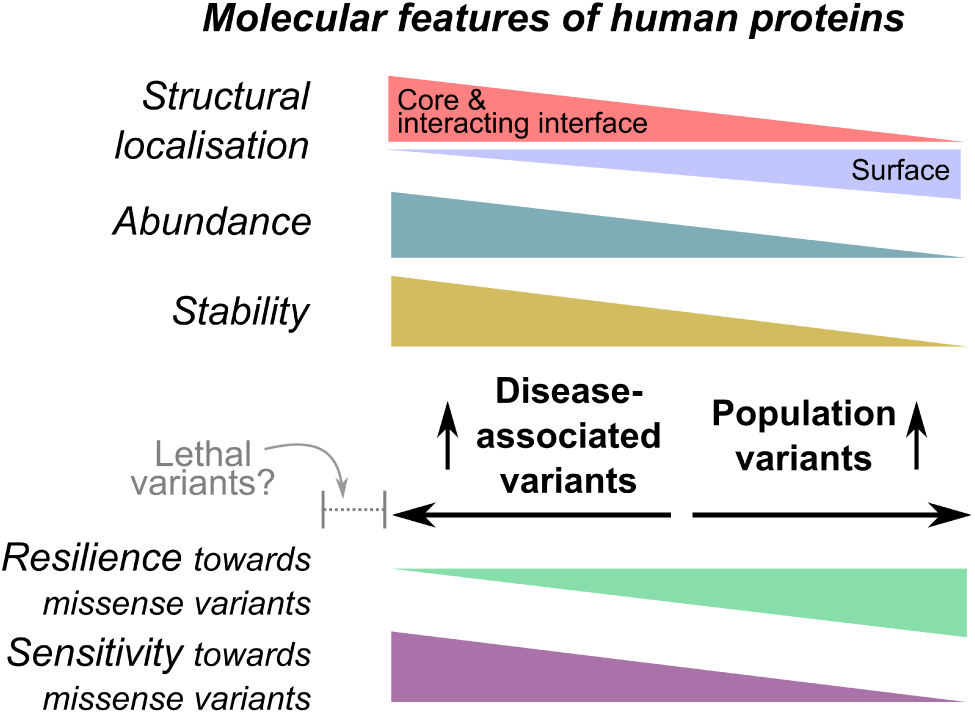
Summary of analyses. Here we have explored patterns of structural localisation, abundance and stability for proteins enriched in disease-associated and population variants respectively. These molecular attributes determine the resilience and sensitivity of the proteins towards missense variants (see Discussion).

Our work highlights a set of rules in predicting the impact of variants. For instance, one could be fairly confident that a variant can be disruptive if it localises to the core of an abundant, stable protein. This type of variant annotation could be valuable to clinicians in interpreting variants observable in any given patient. The detailed set of features we provide could also be harnessed for more systematic improvement of variant impact prediction and interpretation. Firstly, the analysis concerning protein stability suggests it is important to consider the base-line stability of the protein in question when assessing the impact a variant could bring. A number of algorithms have used the estimated change in protein stability upon mutation (∆∆*G*) as a proxy for variant impact. From their analysis of the ProTherm database, Serahijos and colleagues [87] found that mutations in more stable proteins generally led to greater destabilisation (∆∆*G* variation). They interpret this as suggesting that proteins which have evolved to become more stable are in a state closer to their peak stability, where any changes will result in strong destabilisation. Similarly, Pucci and Rooman [88] used temperature-dependent statistical potentials to investigate the thermal stability of the structurome (all proteins with resolved structure), and concluded that mutations in proteins which are highly thermally stable lead to a larger decrease in thermal stability, compared with those in less thermally stable proteins. We believe that our results point to the fact that, even under a scenario in which mutations in proteins with higher stability result in a greater change in stability, a mutation in an already unstable protein is more likely to result in complete/partial unfolding under physiological conditions. This is likely to be relevant to globular proteins, whereas for other types of proteins, e.g. intrinsically disordered proteins, function will be related less to the fold and the density of the protein core. These factors should be brought into consideration when the impact of missense variants.

Secondly, we show that greater insight into the properties of variants in health and disease can be obtained by combining protein structural and functional pathway information. For example, it can be clearly seen that population variants are most enriched on the surface of proteins which take part in pathways we have defined as belonging to the “proliferation” cluster (Figure 4B). Moreover, pathways belonging to this cluster also appear to be enriched in proteins with less thermal stability (Figure 7C), suggesting a possible mechanistic basis underlying the localisation of variants (variants tend to localise to the surface and avoid disrupting the core of these already unstable proteins). This indicates that the combined use of such features may aid in both improving the prediction of variant impact, and in assessing the underlying molecular mechanisms.

Thirdly, our analysis highlights the tissue specificity of variant impact, in terms of the stability and abundance of the altered protein. Our association analysis (Figure 7) of variant enrichment with proteomics features complements a body of research which concludes that the rate of protein evolution correlates negatively with protein expression and abundance [89], the extent of which has been found to be tissue-specific; those tissues with a high neuron density demonstrating the highest anti-correlation [90]. Consistent with this, we found the largest negative correlation for the protein-wise enrichment of rare variants, from the gnomAD dataset, with protein abundance in the brain, and, interestingly also in the ovary and testis, which both harbour long-lived germline progenitor cells (Figure 7B; SI Appendix Figure S10). Purportedly the lifespan of long-lived cells renders them more sensitive to (and therefore necessitate protective strategies [91] against) the toxicity of misfolded proteins. Our analysis highlights the importance of considering the underlying context, specific to the affected organ alongside with the abundance and stability levels of the affected proteins, in assessing the potential impact a missense variant could pose.

Fourthly, the wealth of data presented here could have implications in the development of therapeutic strategies. Rare population variants are known to be abundant in known drug targets, potentially modulating disease risk and drug response [92]. Here we envisage that our domain-centric landscape of variant enrichment (Figure 6), which includes the mapping of targeted drugs, besides providing another feature for the characterisation of variants, will allow for more informed decisions in optimising therapeutic strategies. For example, targets with few population variants could be selected, to minimise differential drug response due to genetic differences between individuals. Interestingly it has recently been shown that genetic variants in such domains (GPCRs), identified in the general population, may be associated with differential drug response between individuals [93]. By viewing variant enrichment and drug availability together, such a domain-centric landscape of variant localisation has implications useful for both understanding variant impact and motivating therapeutic design.

In conclusion, our work highlights the complex interplay between different factors which may determine variant pathogenicity, from atomistic protein structural features (“microscopic”) to large-scale (“macroscopic”) functional pathways and proteomics features. We believe that these insights will prove important in the prioritisation of likely disease-associated variants, and the prediction of variants which drive disease phenotypes. Moreover, the ZoomVar database, which we have made available at http://fraternalilab.kcl.ac.uk/ZoomVar, will facilitate users in the structural analysis of variants. A script is downloadable from the site to allow large-scale programmatic access to the webpage for the structural annotation of user-input variant data; we also provide in the webpage precomputed data underlying all analyses presented here. Further advancement in the structural coverage of the proteome, and the exploitation of high throughput proteomics technologies, such as those analysed here [23, 79], will ultimately offer a finer-grained picture of features which segregate variants in “health” and “disease”.

## 6 Acknowledgement

We thank Dr. Jens Kleinjung for his critical reading of and comments on this manuscript, and members of the Fraternali group for valuable discussion. This research was supported by the British Heart Foundation (RE/13/2/30182 to FF and AL), Croucher Foundation Hong Kong (to JCN) and the Medical Research Council (MR/L01257X/1 to FF).

## References

[1] Adrián Blanco-Gómez, Sonia Castillo-Lluva, María Del Mar Sáez-Freire, Lourdes Hontecillas-Prieto, Jian Hua Mao, Andrés Castellanos-Martín, and Jesus Pérez-Losada. Missing heritability of complex diseases: Enlightenment by genetic variants from intermediate phenotypes. BioEssays: news and reviews in molecular, cellular and developmental biology, 38(7):664–73, 07 2016.

[2] Santhosh Girirajan. Missing heritability and where to find it. Genome biology, 18(1):89, 05 2017.

[3] Luisa Azevedo, Matthew Mort, Antonio C Costa, Raquel M Silva, Dulce Quelhas, Antonio Amorim, and David N Cooper. Improving the in silico assessment of pathogenicity for compensated variants. European journal of human genetics: EJHG, 25(1):2–7, 01 2016.

[4] Wei-Feng Guo, Shao-Wu Zhang, Li-Li Liu, Fei Liu, Qian-Qian Shi, Lei Zhang, Ying Tang, Tao Zeng, and Luonan Chen. Discovering personalized driver mutation profiles of single samples in cancer by network control strategy. Bioinformatics (Oxford, England), 34(11):1893–1903, Jun 2018.

[5] Line Lykke Andersen, Ewa Terczyńska-Dyla, Nanna Mørk, Carsten Scavenius, Jan J Enghild, Klara Höning, Veit Hornung, Mette Christiansen, Trine H Mogensen, and Rune Hartmann. Frequently used bioinformatics tools overestimate the damaging effect of allelic variants. Genes and immunity, Dec 2017.

[6] M Miller, Y Bromberg, and L Swint-Kruse. Computational predictors fail to identify amino acid substitution effects at rheostat positions. Scientific reports, 7:41329, Jan 2017.

[7] Nilah M Ioannidis, Joseph H Rothstein, Vikas Pejaver, Sumit Middha, Shannon K McDonnell, Saurabh Baheti, Anthony Musolf, Qing Li, Emily Holzinger, Danielle Karyadi, Lisa A Cannon-Albright, Craig C Teerlink, Janet L Stanford, William B Isaacs, Jianfeng Xu, Kathleen A Cooney, Ethan M Lange, Johanna Schleutker, John D Carpten, Isaac J Powell, Olivier Cussenot, Geraldine Cancel-Tassin, Graham G Giles, Robert J MacInnis, Christiane Maier, Chih-Lin Hsieh, Fredrik Wiklund, William J Catalona, William D Foulkes, Diptasri Mandal, Rosalind A Eeles, Zsofia Kote-Jarai, Carlos D Bustamante, Daniel J Schaid, Trevor Hastie, Elaine A Ostrander, Joan E Bailey-Wilson, Predrag Radivojac, Stephen N Thibodeau, Alice S Whittemore, and Weiva Sieh. Revel: An ensemble method for predicting the pathogenicity of rare missense variants. American journal of human genetics, 99(4):877–885, Oct 2016.

[8] Rong Chen, Lisong Shi, Jörg Hakenberg, Brian Naughton, Pamela Sklar, Jianguo Zhang, Hanlin Zhou, Lifeng Tian, Om Prakash, Mathieu Lemire, Patrick Sleiman, Wei-Yi Cheng, Wanting Chen, Hardik Shah, Yulan Shen, Menachem Fromer, Larsson Omberg, Matthew A Deardorff, Elaine Zackai, Jason R Bobe, Elissa Levin, Thomas J Hudson, Leif Groop, Jun Wang, Hakon Hakonarson, Anne Wojcicki, George A Diaz, Lisa Edelmann, Eric E Schadt, and Stephen H Friend. Analysis of 589,306 genomes identifies individuals resilient to severe mendelian childhood diseases. Nature biotechnology, 34(5):531–8, 05 2016.

[9] Yannick Mahlich, Jonas Reeb, Maximilian Hecht, Maria Schelling, Tjaart Andries Petrus De Beer, Yana Bromberg, and Burkhard Rost. Common sequence variants affect molecular function more than rare variants? Scientific reports, 7(1):1608, May 2017.

[10] Eman Alhuzimi, Luis G Leal, Michael J E Sternberg, and Alessia David. Properties of human genes guided by their enrichment in rare and common variants. Human mutation, 39(3):365–370, Mar 2018.

[11] Anna Laddach, Joseph Chi-Fung Ng, Sun Sook Chung, and Franca Fraternali. Genetic variants and protein-protein interactions: a multidimensional network-centric view. Current opinion in structural biology, 50:82–90, Jan 2018.

[12] Hui-Chun Lu, Julián Herrera Braga, and Franca Fraternali. Pinsnps: structural and functional analysis of snps in the context of protein interaction networks. Bioinformatics (Oxford, England), 32(16):2534–6, 08 2016.

[13] A Gress, V Ramensky, and O V Kalinina. Spatial distribution of disease-associated variants in three-dimensional structures of protein complexes. Oncogenesis, 6(9):e380, Sep 2017.

[14] H Billur Engin, Jason F Kreisberg, and Hannah Carter. Structure-based analysis reveals cancer missense mutations target protein interaction interfaces. PloS one, 11(4):e0152929, 2016.

[15] Mu Gao, Hongyi Zhou, and Jeffrey Skolnick. Insights into disease-associated mutations in the human proteome through protein structural analysis. Structure (London, England: 1993), 23(7):1362–9, Jul 2015.

[16] Alessia David, Rozami Razali, Mark N Wass, and Michael J E Sternberg. Protein-protein interaction sites are hot spots for disease-associated nonsynonymous snps. Human mutation, 33(2):359–63, Feb 2012.

[17] Douglas E V Pires, David B Ascher, and Tom L Blundell. mcsm: predicting the effects of mutations in proteins using graph-based signatures. Bioinformatics (Oxford, England), 30(3):335–42, Feb 2014.

[18] Eduard Porta-Pardo and Adam Godzik. e-driver: a novel method to identify protein regions driving cancer. Bioinformatics (Oxford, England), 30(21):3109–14, Nov 2014.

[19] Eduard Porta-Pardo, Thomas Hrabe, and Adam Godzik. Cancer3d: understanding cancer mutations through protein structures. Nucleic acids research, 43(Database issue):D968–73, Jan 2015.

[20] Matthew H Bailey, Collin Tokheim, Eduard Porta-Pardo, Sohini Sengupta, Denis Bertrand, Amila Weerasinghe, Antonio Colaprico, Michael C Wendl, Jaegil Kim, Brendan Reardon, Patrick Kwok-Shing Ng, Kang Jin Jeong, Song Cao, Zixing Wang, Jianjiong Gao, Qingsong Gao, Fang Wang, Eric Minwei Liu, Loris Mularoni, Carlota Rubio-Perez, Niranjan Nagarajan, Isidro Cortés-Ciriano, Daniel Cui Zhou, Wen-Wei Liang, Julian M Hess, Venkata D Yellapantula, David Tamborero, Abel Gonzalez-Perez, Chayaporn Suphavilai, Jia Yu Ko, Ekta Khurana, Peter J Park, Eliezer M Van Allen, Han Liang, Michael S Lawrence, Adam Godzik, Nuria Lopez-Bigas, Josh Stuart, David Wheeler, Gad Getz, Ken Chen, Alexander J Lazar, Gordon B Mills, Rachel Karchin, and Li Ding. Comprehensive characterization of cancer driver genes and mutations. Cell, 173(2):371–385.e18, Apr 2018.

[21] Holger Franken, Toby Mathieson, Dorothee Childs, Gavain M A Sweetman, Thilo Werner, Ina Tögel, Carola Doce, Stephan Gade, Marcus Bantscheff, Gerard Drewes, Friedrich B M Reinhard, Wolfgang Huber, and Mikhail M Savitski. Thermal proteome profiling for unbiased identification of direct and indirect drug targets using multiplexed quantitative mass spectrometry. Nature protocols, 10(10):1567–93, Oct 2015.

[22] Mingcong Wang, Christina J Herrmann, Milan Simonovic, Damian Szklarczyk, and Christian von Mering. Version 4.0 of paxdb: Protein abundance data, integrated across model organisms, tissues, and cell-lines. Proteomics, 15(18):3163–8, Sep 2015.

[23] Toby Mathieson, Holger Franken, Jan Kosinski, Nils Kurzawa, Nico Zinn, Gavain Sweetman, Daniel Poeckel, Vikram S Ratnu, Maike Schramm, Isabelle Becher, Michael Steidel, Kyung-Min Noh, Giovanna Bergamini, Martin Beck, Marcus Bantscheff, and Mikhail M Savitski. Systematic analysis of protein turnover in primary cells. Nature communications, 9(1):689, 02 2018.

[24] GTEx Consortium. The genotype-tissue expression (gtex) project. Nature genetics, 45(6):580–5, Jun 2013.

[25] Arun Prasad Pandurangan, David B Ascher, Sherine E Thomas, and Tom L Blundell. Genomes, structural biology and drug discovery: combating the impacts of mutations in genetic disease and antibiotic resistance. Biochemical Society transactions, 45(2):303–311, 04 2017.

[26] Melissa J Landrum, Jennifer M Lee, Mark Benson, Garth Brown, Chen Chao, Shanmuga Chitipiralla, Baoshan Gu, Jennifer Hart, Douglas Hoffman, Jeffrey Hoover, Wonhee Jang, Kenneth Katz, Michael Ovetsky, George Riley, Amanjeev Sethi, Ray Tully, Ricardo Villamarin-Salomon, Wendy Rubinstein, and Donna R Maglott. Clinvar: public archive of interpretations of clinically relevant variants. Nucleic acids research, 44(D1):D862–8, Jan 2016.

[27] Simon A Forbes, David Beare, Prasad Gunasekaran, Kenric Leung, Nidhi Bindal, Harry Boutselakis, Minjie Ding, Sally Bamford, Charlotte Cole, Sari Ward, Chai Yin Kok, Mingming Jia, Tisham De, Jon W Teague, Michael R Stratton, Ultan McDermott, and Peter J Campbell. Cosmic: exploring the world’s knowledge of somatic mutations in human cancer. Nucleic acids research, 43(Database issue):D805–11, Jan 2015.

[28] Monkol Lek, Konrad J Karczewski, Eric V Minikel, Kaitlin E Samocha, Eric Banks, Timothy Fennell, Anne H O’Donnell-Luria, James S Ware, Andrew J Hill, Beryl B Cummings, Taru Tukiainen, Daniel P Birnbaum, Jack A Kosmicki, Laramie E Duncan, Karol Estrada, Fengmei Zhao, James Zou, Emma Pierce-Hoffman, Joanne Berghout, David N Cooper, Nicole Deflaux, Mark DePristo, Ron Do, Jason Flannick, Menachem Fromer, Laura Gauthier, Jackie Goldstein, Namrata Gupta, Daniel Howrigan, Adam Kiezun, Mitja I Kurki, Ami Levy Moonshine, Pradeep Natarajan, Lorena Orozco, Gina M Peloso, Ryan Poplin, Manuel A Rivas, Valentin Ruano-Rubio, Samuel A Rose, Douglas M Ruderfer, Khalid Shakir, Peter D Stenson, Christine Stevens, Brett P Thomas, Grace Tiao, Maria T Tusie-Luna, Ben Weisburd, Hong-Hee Won, Dongmei Yu, David M Altshuler, Diego Ardissino, Michael Boehnke, John Danesh, Stacey Donnelly, Roberto Elosua, Jose C Florez, Stacey B Gabriel, Gad Getz, Stephen J Glatt, Christina M Hultman, Sekar Kathiresan, Markku Laakso, Steven McCarroll, Mark I McCarthy, Dermot McGovern, Ruth McPherson, Benjamin M Neale, Aarno Palotie, Shaun M Purcell, Danish Saleheen, Jeremiah M Scharf, Pamela Sklar, Patrick F Sullivan, Jaakko Tuomilehto, Ming T Tsuang, Hugh C Watkins, James G Wilson, Mark J Daly, and Daniel G MacArthur. Analysis of protein-coding genetic variation in 60,706 humans. Nature, 536(7616):285–91, 08 2016.

[29] Sylvain Poux, Cecilia N Arighi, Michele Magrane, Alex Bateman, Chih-Hsuan Wei, Zhiyong Lu, Emmanuel Boutet, Hema Bye-A-Jee, Maria Livia Famiglietti, Bernd Roechert, and The UniProt Consortium. On expert curation and scalability: Uniprotkb/swiss-prot as a case study. Bioinformatics (Oxford, England), 33(21):3454–3460, Nov 2017.

[30] Bronwen L Aken, Sarah Ayling, Daniel Barrell, Laura Clarke, Valery Curwen, Susan Fairley, Julio Fernandez Banet, Konstantinos Billis, Carlos García Girón, Thibaut Hourlier, Kevin Howe, Andreas Kähäri, Felix Kokocinski, Fergal J Martin, Daniel N Murphy, Rishi Nag, Magali Ruffier, Michael Schuster, Y Amy Tang, Jan-Hinnerk Vogel, Simon White, Amonida Zadissa, Paul Flicek, and Stephen M J Searle. The ensembl gene annotation system. Database: the journal of biological databases and curation, 2016, 2016.

[31] Sun Sook Chung, Anna Laddach, N. Shaun Bevan Thomas, and Franca Fraternali. Short loop motif profiling of protein interaction networks in acute myeloid leukaemia. bioRxiv, 2018.

[32] Sandra Orchard, Mais Ammari, Bruno Aranda, Lionel Breuza, Leonardo Briganti, Fiona Broackes-Carter, Nancy H Campbell, Gayatri Chavali, Carol Chen, Noemi del Toro, Margaret Duesbury, Marine Dumousseau, Eugenia Galeota, Ursula Hinz, Marta Iannuccelli, Sruthi Jagannathan, Rafael Jimenez, Jyoti Khadake, Astrid Lagreid, Luana Licata, Ruth C Lovering, Birgit Meldal, Anna N Melidoni, Mila Milagros, Daniele Peluso, Livia Perfetto, Pablo Porras, Arathi Raghunath, Sylvie Ricard-Blum, Bernd Roechert, Andre Stutz, Michael Tognolli, Kim van Roey, Gianni Cesareni, and Henning Hermjakob. The mintact project–intact as a common curation platform for 11 molecular interaction databases. Nucleic acids research, 42(Database issue):D358–63, Jan 2014.

[33] Andrew Chatr-Aryamontri, Rose Oughtred, Lorrie Boucher, Jennifer Rust, Christie Chang, Nadine K Kolas, Lara O’Donnell, Sara Oster, Chandra Theesfeld, Adnane Sellam, Chris Stark, Bobby-Joe Breitkreutz, Kara Dolinski, and Mike Tyers. The biogrid interaction database: 2017 update. Nucleic acids research, 45(D1):D369–D379, Jan 2017.

[34] Damian Szklarczyk, Andrea Franceschini, Stefan Wyder, Kristoffer Forslund, Davide Heller, Jaime Huerta-Cepas, Milan Simonovic, Alexander Roth, Alberto Santos, Kalliopi P Tsafou, Michael Kuhn, Peer Bork, Lars J Jensen, and Christian von Mering. String v10: protein-protein interaction networks, integrated over the tree of life. Nucleic acids research, 43(Database issue):D447–52, Jan 2015.

[35] Ioannis Xenarios, Lukasz Salwínski, Xiaoqun Joyce Duan, Patrick Higney, Sul-Min Kim, and David Eisenberg. Dip, the database of interacting proteins: a research tool for studying cellular networks of protein interactions. Nucleic acids research, 30(1):303–5, Jan 2002.

[36] Suraj Peri, J Daniel Navarro, Troels Z Kristiansen, Ramars Amanchy, Vineeth Surendranath, Babylakshmi Muthusamy, T K B Gandhi, K N Chandrika, Nandan Deshpande, Shubha Suresh, B P Rashmi, K Shanker, N Padma, Vidya Niranjan, H C Harsha, Naveen Talreja, B M Vrushabendra, M A Ramya, A J Yatish, Mary Joy, H N Shivashankar, M P Kavitha, Minal Menezes, Dipanwita Roy Choudhury, Neelanjana Ghosh, R Saravana, Sreenath Chandran, Sujatha Mohan, Chandra Kiran Jonnalagadda, C K Prasad, Chandan Kumar-Sinha, Krishna S Deshpande, and Akhilesh Pandey. Human protein reference database as a discovery resource for proteomics. Nucleic acids research, 32(Database issue):D497–501, Jan 2004.

[37] Pierre C Havugimana, G Traver Hart, Tamás Nepusz, Haixuan Yang, Andrei L Turinsky, Zhihua Li, Peggy I Wang, Daniel R Boutz, Vincent Fong, Sadhna Phanse, Mohan Babu, Stephanie A Craig, Pingzhao Hu, Cuihong Wan, James Vlasblom, Vaqaar un Nisa Dar, Alexandr Bezginov, Gregory W Clark, Gabriel C Wu, Shoshana J Wodak, Elisabeth R M Tillier, Alberto Paccanaro, Edward M Marcotte, and Andrew Emili. A census of human soluble protein complexes. Cell, 150(5):1068–81, Aug 2012.

[38] Thomas Rolland, Murat Taşan, Benoit Charloteaux, Samuel J Pevzner, Quan Zhong, Nidhi Sahni, Song Yi, Irma Lemmens, Celia Fontanillo, Roberto Mosca, Atanas Kamburov, Susan D Ghiassian, Xinping Yang, Lila Ghamsari, Dawit Balcha, Bridget E Begg, Pascal Braun, Marc Brehme, Martin P Broly, Anne-Ruxandra Carvunis, Dan Convery-Zupan, Roser Corominas, Jasmin Coulombe-Huntington, Elizabeth Dann, Matija Dreze, Amélie Dricot, Changyu Fan, Eric Franzosa, Fana Gebreab, Bryan J Gutierrez, Madeleine F Hardy, Mike Jin, Shuli Kang, Ruth Kiros, Guan Ning Lin, Katja Luck, Andrew MacWilliams, Jörg Menche, Ryan R Murray, Alexandre Palagi, Matthew M Poulin, Xavier Rambout, John Rasla, Patrick Reichert, Viviana Romero, Elien Ruyssinck, Julie M Sahalie, Annemarie Scholz, Akash A Shah, Amitabh Sharma, Yun Shen, Kerstin Spirohn, Stanley Tam, Alexander O Tejeda, Shelly A Trigg, Jean-Claude Twizere, Kerwin Vega, Jennifer Walsh, Michael E Cusick, Yu Xia, Albert-László Barabási, Lilia M Iakoucheva, Patrick Aloy, Javier De Las Rivas, Jan Tavernier, Michael A Calderwood, David E Hill, Tong Hao, Frederick P Roth, and Marc Vidal. A proteome-scale map of the human interactome network. Cell, 159(5):1212–1226, Nov 2014.

[39] Edward L Huttlin, Lily Ting, Raphael J Bruckner, Fana Gebreab, Melanie P Gygi, John Szpyt, Stanley Tam, Gabriela Zarraga, Greg Colby, Kurt Baltier, Rui Dong, Virginia Guarani, Laura Pontano Vaites, Alban Ordureau, Ramin Rad, Brian K Erickson, Martin Wühr, Joel Chick, Bo Zhai, Deepak Kolippakkam, Julian Mintseris, Robert A Obar, Tim Harris, Spyros Artavanis-Tsakonas, Mathew E Sowa, Pietro De Camilli, Joao A Paulo, J Wade Harper, and Steven P Gygi. The bioplex network: A systematic exploration of the human interactome. Cell, 162(2):425–440, Jul 2015.

[40] S F Altschul, T L Madden, A A Schäffer, J Zhang, Z Zhang, W Miller, and D J Lipman. Gapped blast and psi-blast: a new generation of protein database search programs. Nucleic acids research, 25(17):3389–402, Sep 1997.

[41] Robert D Finn, Penelope Coggill, Ruth Y Eberhardt, Sean R Eddy, Jaina Mistry, Alex L Mitchell, Simon C Potter, Marco Punta, Matloob Qureshi, Amaia Sangrador-Vegas, Gustavo A Salazar, John Tate, and Alex Bateman. The pfam protein families database: towards a more sustainable future. Nucleic acids research, 44(D1):D279–85, Jan 2016.

[42] Robert D Finn, Jody Clements, and Sean R Eddy. Hmmer web server: interactive sequence similarity searching. Nucleic acids research, 39(Web Server issue):W29–37, Jul 2011.

[43] C Notredame, D G Higgins, and J Heringa. T-coffee: A novel method for fast and accurate multiple sequence alignment. Journal of molecular biology, 302(1):205–17, Sep 2000.

[44] E W Myers and W Miller. Optimal alignments in linear space. Computer applications in the biosciences: CABIOS, 4(1):11–7, Mar 1988.

[45] Li C Xue, Drena Dobbs, and Vasant Honavar. Homppi: a class of sequence homology based protein-protein interface prediction methods. BMC bioinformatics, 12:244, Jun 2011.

[46] Jens Kleinjung and Franca Fraternali. Popscomp: an automated interaction analysis of biomolecular complexes. Nucleic acids research, 33(Web Server issue):W342–6, Jul 2005.

[47] Arianna Fornili, Alessandro Pandini, Hui-Chun Lu, and Franca Fraternali. Specialized dynamical properties of promiscuous residues revealed by simulated conformational ensembles. Journal of Chemical Theory and Computation, 9(11):5127–5147, Nov 2013.

[48] Luigi Cavallo, Jens Kleinjung, and Franca Fraternali. Pops: A fast algorithm for solvent accessible surface areas at atomic and residue level. Nucleic acids research, 31(13):3364–6, Jul 2003.

[49] David T Jones and Domenico Cozzetto. Disopred3: precise disordered region predictions with annotated protein-binding activity. Bioinformatics (Oxford, England), 31(6):857–63, Mar 2015.

[50] Peter V Hornbeck, Indy Chabra, Jon M Kornhauser, Elzbieta Skrzypek, and Bin Zhang. Phosphosite: A bioinformatics resource dedicated to physiological protein phosphorylation. Proteomics, 4(6):1551–61, Jun 2004.

[51] Bethan Yates, Bryony Braschi, Kristian A Gray, Ruth L Seal, Susan Tweedie, and Elspeth A Bruford. Genenames.org: the hgnc and vgnc resources in 2017. Nucleic acids research, 45(D1):D619–D625, Jan 2017.

[52] S. Kumar, D. Clarke, and M. B. Gerstein. Leveraging protein dynamics to identify cancer mutational hotspots using 3D structures. Proc. Natl. Acad. Sci. U.S.A., 116(38):18962–18970, Sep 2019.

[53] P. Ashford, C. S. M. Pang, A. A. Moya-Garcia, T. Adeyelu, and C. A. Orengo. A CATH domain functional family based approach to identify putative cancer driver genes and driver mutations. Sci Rep, 9(1):263, Jan 2019.

[54] R Michael Sivley, Xiaoyi Dou, Jens Meiler, William S Bush, and John A Capra. Comprehensive analysis of constraint on the spatial distribution of missense variants in human protein structures. American journal of human genetics, 102(3):415–426, Mar 2018.

[55] Alexey Sergushichev. An algorithm for fast preranked gene set enrichment analysis using cumulative statistic calculation. bioRxiv, 2016.

[56] Aravind Subramanian, Pablo Tamayo, Vamsi K Mootha, Sayan Mukherjee, Benjamin L Ebert, Michael A Gillette, Amanda Paulovich, Scott L Pomeroy, Todd R Golub, Eric S Lander, and Jill P Mesirov. Gene set enrichment analysis: a knowledge-based approach for interpreting genome-wide expression profiles. Proceedings of the National Academy of Sciences of the United States of America, 102(43):15545–50, Oct 2005.

[57] Minoru Kanehisa, Miho Furumichi, Mao Tanabe, Yoko Sato, and Kanae Morishima. Kegg: new perspectives on genomes, pathways, diseases and drugs. Nucleic acids research, 45(D1):D353–D361, 01 2017.

[58] Malika Charrad, Nadia Ghazzali, Véronique Boiteau, and Azam Niknafs. NbClust: An R package for determining the relevant number of clusters in a data set. Journal of Statistical Software, 61(6):1–36, 2014.

[59] Angelo Canty and B. D. Ripley. boot: Bootstrap R (S-Plus) Functions, 2017. R package version 1.3-20.

[60] Andri Signorellmult. et al. DescTools: Tools for Descriptive Statistics, 2017. R package version 0.99.19.

[61] Gregory R. Warnes, Ben Bolker, Lodewijk Bonebakker, Robert Gentleman, Wolfgang Huber Andy Liaw, Thomas Lumley, Martin Maechler, Arni Magnusson, Steffen Moeller, Marc Schwartz, and Bill Venables. gplots: Various R Programming Tools for Plotting Data, 2016. R package version 3.0.1.

[62] Zuguang Gu, Roland Eils, and Matthias Schlesner. Complex heatmaps reveal patterns and correlations in multidimensional genomic data. Bioinformatics, 2016.

[63] Martin Krzywinski, Jacqueline Schein, Inanç Birol, Joseph Connors, Randy Gascoyne, Doug Horsman, Steven J Jones, and Marco A Marra. Circos: an information aesthetic for comparative genomics. Genome research, 19(9):1639–45, Sep 2009.

[64] Travis E Oliphant. A guide to NumPy, volume 1. Trelgol Publishing USA, 2006.

[65] Eduard Porta-Pardo, Luz Garcia-Alonso, Thomas Hrabe, Joaquin Dopazo, and Adam Godzik. A pan-cancer catalogue of cancer driver protein interaction interfaces. PLoS computational biology, 11(10):e1004518, Oct 2015.

[66] Bert Vogelstein, Nickolas Papadopoulos, Victor E Velculescu, Shibin Zhou, Luis A Diaz, and Kenneth W Kinzler. Cancer genome landscapes. Science (New York, N.Y.), 339(6127):1546–58, Mar 2013.

[67] Henning Stehr, Seon-Hi J Jang, José M Duarte, Christoph Wierling, Hans Lehrach, Michael Lappe, and Bodo M H Lange. The structural impact of cancer-associated missense mutations in oncogenes and tumor suppressors. Molecular cancer, 10:54, May 2011.

[68] Jüri Reimand, Omar Wagih, and Gary D Bader. The mutational landscape of phosphorylation signaling in cancer. Scientific reports, 3:2651, Oct 2013.

[69] Aleksandra Olow, Zhongzhong Chen, R Hannes Niedner, Denise M Wolf, Christina Yau, Aleksandr Pankov, Evelyn Pei Rong Lee, Lamorna Brown-Swigart, Laura J van‘t Veer, and Jean-Philippe Coppé. An atlas of the human kinome reveals the mutational landscape underlying dysregulated phosphorylation cascades in cancer. Cancer research, 76(7):1733–45, 04 2016.

[70] D Menzies and H Ellis. The role of plasminogen activator in adhesion prevention. Surgery, gynecology and obstetrics, 172(5):362–6, May 1991.

[71] Manoj Garg, Glenn Braunstein, and Harold Phillip Koeffler. Lamc2 as a therapeutic target for cancers. Expert opinion on therapeutic targets, 18(9):979–82, Sep 2014.

[72] Family: Npip (pf06409). https://pfam.xfam.org/family/PF06409. [Online; accessed 13-Mar-2018].

[73] Family: Nut (pf12881). https://pfam.xfam.org/family/PF12881. [Online; accessed 13-Mar-2018].

[74] Gregg L Semenza. Vhl and p53: tumor suppressors team up to prevent cancer. Molecular cell, 22(4):437–9, May 2006.

[75] Ian Sillitoe, Tony E Lewis, Alison Cuff, Sayoni Das, Paul Ashford, Natalie L Dawson, Nicholas Furnham, Roman A Laskowski, David Lee, Jonathan G Lees, Sonja Lehtinen, Romain A Studer, Janet Thornton, and Christine A Orengo. Cath: comprehensive structural and functional annotations for genome sequences. Nucleic acids research, 43(Database issue):D376–81, Jan 2015.

[76] David S Wishart, Yannick D Feunang, An C Guo, Elvis J Lo, Ana Marcu, Jason R Grant, Tanvir Sajed, Daniel Johnson, Carin Li, Zinat Sayeeda, Nazanin Assempour, Ithayavani Iynkkaran, Yifeng Liu, Adam Maciejewski, Nicola Gale, Alex Wilson, Lucy Chin, Ryan Cummings, Diana Le, Allison Pon, Craig Knox, and Michael Wilson. Drugbank 5.0: a major update to the drugbank database for 2018. Nucleic acids research, 46(D1):D1074–D1082, Jan 2018.

[77] Rita Santos, Oleg Ursu, Anna Gaulton, A Patrícia Bento, Ramesh S Donadi, Cristian G Bologa, Anneli Karlsson, Bissan Al-Lazikani, Anne Hersey, Tudor I Oprea, and John P Overington. A comprehensive map of molecular drug targets. Nature reviews. Drug discovery, 16(1):19–34, 01 2017.

[78] Germán Rivas and Allen P Minton. Macromolecular crowding in vitro, in vivo, and in between. Trends in Biochemical Sciences, 41(11):970–981, 11 2016.

[79] Pascal Leuenberger, Stefan Ganscha, Abdullah Kahraman, Valentina Cappelletti, Paul J Boersema, Chritian von Mering, Manfred Claassen, and Paola Picotti. Cell-wide analysis of protein thermal unfolding reveals determinants of thermostability. Science (New York, N.Y.), 355(6327), 02 2017.

[80] A. Keinan and A. G. Clark. Recent explosive human population growth has resulted in an excess of rare genetic variants. Science, 336(6082):740–743, May 2012.

[81] J. A. Tennessen, A. W. Bigham, T. D. O’Connor, W. Fu, E. E. Kenny, S. Gravel, S. McGee, R. Do, X. Liu, G. Jun, H. M. Kang, D. Jordan, S. M. Leal, S. Gabriel, M. J. Rieder, G. Abecasis, D. Altshuler, D. A. Nickerson, E. Boerwinkle, S. Sunyaev, C. D. Bustamante, M. J. Bamshad, and J. M. Akey. Evolution and functional impact of rare coding variation from deep sequencing of human exomes. Science, 337(6090):64–69, Jul 2012.

[82] X. Li, Y. Kim, E. K. Tsang, J. R. Davis, F. N. Damani, C. Chiang, G. T. Hess, Z. Zappala, B. J. Strober, A. J. Scott, A. Li, A. Ganna, M. C. Bassik, J. D. Merker, I. M. Hall, A. Battle, S. B. Montgomery, F. Aguet, K. G. Ardlie, B. B. Cummings, E. T. Gelfand, G. Getz, K. Hadley, R. E. Handsaker, K. H. Huang, S. Kashin, K. J. Karczewski, M. Lek, X. Li, D. G. MacArthur, J. L. Nedzel, D. T. Nguyen, M. S. Noble, A. V. Segre, C. A. Trowbridge, T. Tukiainen, N. S. Abell, B. Balliu, R. Barshir, O. Basha, A. Battle, G. K. Bogu, A. Brown, C. D. Brown, S. E. Castel, L. S. Chen, C. Chiang, D. F. Conrad, N. J. Cox, F. N. Damani, J. R. Davis, O. Delaneau, E. T. Dermitzakis, B. E. Engelhardt, E. Eskin, P. G. Ferreira, L. Fresard, E. R. Gamazon, D. Garrido-Martin, A. D. H. Gewirtz, G. Gliner, M. J. Gloudemans, R. Guigo, I. M. Hall, B. Han, Y. He, F. Hormozdiari, C. Howald, H. Kyung Im, B. Jo, E. Yong Kang, Y. Kim, S. Kim-Hellmuth, T. Lappalainen, G. Li, X. Li, B. Liu, S. Mangul, M. I. McCarthy, I. C. McDowell, P. Mohammadi, J. Monlong, S. B. Montgomery, M. Munoz-Aguirre, A. W. Ndungu, D. L. Nicolae, A. B. Nobel, M. Oliva, H. Ongen, J. J. Palowitch, N. Panousis, P. Papasaikas, Y. Park, P. Parsana, A. J. Payne, C. B. Peterson, J. Quan, F. Reverter, C. Sabatti, A. Saha, M. Sammeth, A. J. Scott, A. A. Shabalin, R. Sodaei, M. Stephens, B. E. Stranger, B. J. Strober, J. H. Sul, E. K. Tsang, S. Urbut, M. van de Bunt, G. Wang, X. Wen, F. A. Wright, H. S. Xi, E. Yeger-Lotem, Z. Zappala, J. B. Zaugg, Y. H. Zhou, J. M. Akey, D. Bates, J. Chan, L. S. Chen, M. Claussnitzer, K. Demanelis, M. Diegel, J. A. Doherty, A. P. Feinberg, M. S. Fernando, J. Halow, K. D. Hansen, E. Haugen, P. F. Hickey, L. Hou, F. Jasmine, R. Jian, L. Jiang, A. Johnson, R. Kaul, M. Kellis, M. G. Kibriya, K. Lee, J. Billy Li, Q. Li, X. Li, J. Lin, S. Lin, S. Linder, C. Linke, Y. Liu, M. T. Maurano, B. Molinie, S. B. Montgomery, J. Nelson, F. J. Neri, M. Oliva, Y. Park, B. L. Pierce, N. J. Rinaldi, L. F. Rizzardi, R. Sandstrom, A. Skol, K. S. Smith, M. P. Snyder, J. Stamatoyannopoulos, B. E. Stranger, H. Tang, E. K. Tsang, L. Wang, M. Wang, N. Van Wittenberghe, F. Wu, R. Zhang, C. R. Nierras, P. A. Branton, L. J. Carithers, P. Guan, H. M. Moore, A. Rao, J. B. Vaught, S. E. Gould, N. C. Lockart, C. Martin, J. P. Struewing, S. Volpi, A. M. Addington, S. E. Koester, A. R. Little, L. E. Brigham, R. Hasz, M. Hunter, C. Johns, M. Johnson, G. Kopen, W. F. Leinweber, J. T. Lonsdale, A. McDonald, B. Mestichelli, K. Myer, B. Roe, M. Salvatore, S. Shad, J. A. Thomas, G. Walters, M. Washington, J. Wheeler, J. Bridge, B. A. Foster, B. M. Gillard, E. Karasik, R. Kumar, M. Miklos, M. T. Moser, S. D. Jewell, R. G. Montroy, D. C. Rohrer, D. R. Valley, D. A. Davis, D. C. Mash, A. H. Undale, A. M. Smith, D. E. Tabor, N. V. Roche, J. A. McLean, N. Vatanian, K. L. Robinson, L. Sobin, M. E. Barcus, K. M. Valentino, L. Qi, S. Hunter, P. Hariharan, S. Singh, K. S. Um, T. Matose, M. M. Tomaszewski, L. K. Barker, M. Mosavel, L. A. Siminoff, H. M. Traino, P. Flicek, T. Juettemann, M. Ruffier, D. Sheppard, K. Taylor, S. J. Trevanion, D. R. Zerbino, B. Craft, M. Goldman, M. Haeussler, W. J. Kent, C. M. Lee, B. Paten, K. R. Rosenbloom, J. Vivian, J. Zhu, B. Craft, M. Goldman, M. Haeussler, W. J. Kent, C. M. Lee, B. Paten, K. R. Rosenbloom, J. Vivian, and J. Zhu. The impact of rare variation on gene expression across tissues. Nature, 550(7675):239–243, 10 2017.

[83] D. G. MacArthur, S. Balasubramanian, A. Frankish, N. Huang, J. Morris, K. Walter, L. Jostins, L. Habegger, J. K. Pickrell, S. B. Montgomery, C. A. Albers, Z. D. Zhang, D. F. Conrad, G. Lunter, H. Zheng, Q. Ayub, M. A. DePristo, E. Banks, M. Hu, R. E. Handsaker, J. A. Rosenfeld, M. Fromer, M. Jin, X. J. Mu, E. Khurana, K. Ye, M. Kay, G. I. Saunders, M. M. Suner, T. Hunt, I. H. Barnes, C. Amid, D. R. Carvalho-Silva, A. H. Bignell, C. Snow, B. Yngvadottir, S. Bumpstead, D. N. Cooper, Y. Xue, I. G. Romero, J. Wang, Y. Li, R. A. Gibbs, S. A. McCarroll, E. T. Dermitzakis, J. K. Pritchard, J. C. Barrett, J. Harrow, M. E. Hurles, M. B. Gerstein, and C. Tyler-Smith. A systematic survey of loss-of-function variants in human protein-coding genes. Science, 335(6070):823–828, Feb 2012.

[84] Lluis Quintana-Murci. Understanding rare and common diseases in the context of human evolution. Genome biology, 17(1):225, 11 2016.

[85] Igor Orlov, Alexander G Myasnikov, Leonid Andronov, S Kundhavai Natchiar, Heena Khatter, Brice Bein-steiner, Jean-François Méńetret, Isabelle Hazemann, Kareem Mohideen, Karima Tazibt, Rachel Tabaroni, Hanna Kratzat, Nadia Djabeur, Tatiana Bruxelles, Finaritra Raivoniaina, Lorenza di Pompeo, Morgan Torchy, Isabelle Billas, Alexandre Urzhumtsev, and Bruno P Klaholz. The integrative role of cryo electron microscopy in molecular and cellular structural biology. Biology of the cell, 109(2):81–93, Feb 2017.

[86] Stephen K Burley, Genji Kurisu, John L Markley, Haruki Nakamura, Sameer Velankar, Helen M Berman, Andrej Sali, Torsten Schwede, and Jill Trewhella. Pdb-dev: a prototype system for depositing integrative/hybrid structural models. Structure (London, England: 1993), 25(9):1317–1318, 09 2017.

[87] Adrian W R Serohijos, Zilvinas Rimas, and Eugene I Shakhnovich. Protein biophysics explains why highly abundant proteins evolve slowly. Cell reports, 2(2):249–56, Aug 2012.

[88] Fabrizio Pucci and Marianne Rooman. Improved insights into protein thermal stability: from the molecular to the structurome scale. Philosophical transactions. Series A, Mathematical, physical, and engineering sciences, 374(2080), Nov 2016.

[89] Jianzhi Zhang and Jian-Rong Yang. Determinants of the rate of protein sequence evolution. Nature reviews. Genetics, 16(7):409–20, Jul 2015.

[90] D Allan Drummond and Claus O Wilke. Mistranslation-induced protein misfolding as a dominant constraint on coding-sequence evolution. Cell, 134(2):341–52, Jul 2008.

[91] D. M. Walther, P. Kasturi, M. Zheng, S. Pinkert, G. Vecchi, P. Ciryam, R. I. Morimoto, C. M. Dobson, M. Vendruscolo, M. Mann, and F. U. Hartl. Widespread Proteome Remodeling and Aggregation in Aging C. elegans. Cell, 161(4):919–932, May 2015.

[92] M. R. Nelson, D. Wegmann, M. G. Ehm, D. Kessner, P. St Jean, C. Verzilli, J. Shen, Z. Tang, S. A. Bacanu, D. Fraser, L. Warren, J. Aponte, M. Zawistowski, X. Liu, H. Zhang, Y. Zhang, J. Li, Y. Li, L. Li, P. Woollard, S. Topp, M. D. Hall, K. Nangle, J. Wang, G. Abecasis, L. R. Cardon, S. Zollner, J. C. Whittaker, S. L. Chissoe, J. Novembre, and V. Mooser. An abundance of rare functional variants in 202 drug target genes sequenced in 14,002 people. Science, 337(6090):100–104, Jul 2012.

[93] Alexander S Hauser, Sreenivas Chavali, Ikuo Masuho, Leonie J Jahn, Kirill A Martemyanov, David E Gloriam, and M Madan Babu. Pharmacogenomics of gpcr drug targets. Cell, 172(1-2):41–54.e19, Jan 2018.

