## Supplementary Information for "Missense variants in health and disease affect distinct functional pathways and proteomics features"

### S1 Supplementary Methods

#### S1.1 Data sources

##### S1.1.1 Variant data

ClinVar (dbSNP BUILD ID 149) variant data [1], COSMIC coding mutations (v80) [2] and gnomAD exome data [3], all mapped to the GRCh37 genome build, were obtained in variant call format (VCF). The ClinVar dataset contains variants submitted through clinical channels. Only variants with CLINSIG codes 4 and 5 (i.e. those classified as "probably pathogenic" or "pathogenic") were selected for further analysis. To ensure the quality of our dataset, we selected only variants with "variant suspect reason code" of 0 (unspecified). Additionally, all variants labelled as being somatic were filtered from this dataset. All variant datasets were mapped to Ensembl protein sequences [4] using the Variant Effect Predictor (VEP) [5], and further mapped to canonical UniProt sequences and the respective structures/homologs.

##### S1.1.2 Protein-protein interaction network

A large non-redundant protein-protein interaction network (UniPPIN) [6] was used. This incorporates non-redundant interactions amalgamated from IntAct [7], BioGRID [8], STRING [9], DIP [10] and HPRD [11], as well as recent large-scale experimental studies [12, 13, 14].

##### S1.1.3 Protein sequences and structures

The biounit database of the Protein Data Bank (PDB) was downloaded on 28/04/2017. For mapping purposes, in this study, both the canonical UniProt human protein sequences [15] (for mapping to structures and protein-protein interaction networks) and Ensembl protein sequences [4] (for mapping variant datasets) were used.

##### S1.1.4 Gene and protein annotations

Gene sets for KEGG pathways were obtained from the MSigDB database [16]. Oncogene and tumour suppressor gene annotations were taken from Supplementary Table S2A from Vogelstein and colleagues [17]. Cancer drivers were taken from the Cancer Gene Census (CGC) (COSMIC v84). Genes from both tiers 1 and 2 were included. Conversions between gene symbols, Entrez gene identifiers and UniProt accession numbers were performed using the biomartR package [18, 19]. A list of DNA-binding domains was obtained from the review by Vaquerizas and colleagues [20]. These domains were mapped from InterPro [21] IDs to Pfam IDs using conversion tables in Pfam (v31).

###### S1.1.5 Protein-drug interaction mapping

A mapping of protein-drug interactions was obtained from DrugBank (v5.0.11) [22] (under "Target Drug-UniProt Links") and filtered for human proteins. Drugs were mapped to a Pfam domain-type if at least one domain of that type occurs in a protein a drug is known to interact with. It is, of course, possible that a drug may only interact directly with another domain-type within the protein. However, this approach was chosen due to the fact that if only domain-drug interactions with supporting structural information are accepted, the data becomes both sparse and biased towards structurally resolved domains.

###### S1.1.6 Protemics and transcriptomics data

Protein thermal stability, abundance and half-life data were obtained from separate large-scale studies [23, 24, 25] as detailed in the main text. Gene expression quantification (Reads Per Kilobase of transcript per Million mapped reads [RPKM]) counts per sample (v6p) was downloaded from the GTEx portal [26] and grouped by tissue, according to the sample metadata provided. For each tissue type, we quantified the gene-wise proportion of samples with an RPKM equal to zero. Only those genes with zero counts in $< 10$ % of samples were retained for our analysis.

##### S1.2 ZoomVar Database

###### S1.2.1 Identification of resolved structures/homologs

Canonical UniProt human protein sequences were assigned resolved structures/homologs from the PDB biounit database [27] using BLAST [28]. BLAST searches were carried out using both full-length protein sequences and domain sequences, which were defined by scanning UniProt sequences against the PFAM seed library [29] using HMMER [30]. Hits were only accepted with sequence identity $> 30$ % and E-value $< 0.001$. T-COFFEE [31] was used to obtain a per-residue mapping of queries to structure hits. The quotient solvent accessible surface area [Q(SASA)] of each structure residue was computed using POPS [32].

###### S1.2.2 Mapping of Ensembl proteins

Ensembl protein sequences were mapped to UniProt protein sequences [15], using UniProt ID mapping. Additionally, if UniProt and Ensembl sequences were not of the same length, the sequences were aligned using T-COFFEE [31] to obtain a per-residue mapping. Stretcher [33] was used to align those sequences which were too long to align using T-COFFEE.

##### S1.2.3 Determination of per-residue binding partners

A protein may interact with multiple other proteins. For each of these interactions, a maximum of 10 corresponding best hits (ordered by HomPPI defined score [34]), located in the best populated zone, were considered. If a residue was located at the interaction interface, in at least half of these structures, it was annotated as interacting with that specific protein, otherwise it was annotated as non-interacting.

#### S1.3 Calculation of protein topological network features

Graph representations for protein structures, with mapped missense variants, were constructed. Here protein C$\alpha$ atoms are represented by nodes in the graph, and nodes are connected if C$\alpha$'s are $< 10$ Å apart in 3D space. Similar graph representations are routinely used to create elastic network models of proteins. One class of such models is the Gaussian network model (GNM); the default cut-off distance used by GNMs is commonly 10 Å [35]. Therefore we use this cut-off in the creation networks here. Topological network features were calculated using the python package NetworkX [36]. Specifically, we calculate the degree, degree centrality, betweenness centrality and closeness centrality of residues (nodes) to which variants localise. The degree of a node is defined as the number of nodes it is connected to, and the fraction of connected nodes constitutes its degree centrality. Betweenness centrality $BC$ is defined as the number of pairwise shortest paths which pass through a node:

$$BC(u) = \sum_{s,t \in V} \frac{\sigma(s,t|u)}{\sigma(s,t)} \tag{1}$$

Here V is the set of nodes, $\sigma(s,t)$ is the total number of shortest paths, and $\sigma(s,t|v)$ is the number of shortest paths which pass through the node $u$.

Closeness centrality $CC$ is the reciprocal distance of the sum of the shortest paths from all other $n-1$ nodes to node $v$:

$$CC(u) = \frac{n-1}{\sum_{v=1}^{n-1} d(v,u)} \tag{2}$$

Where $d(v,u)$ is the distance of the shortest path between nodes $v$ and $u$.

#### S1.4 Calculation of protein core density

The density of residue packing in protein cores may impact on their ability to accommodate missense variants. To investigate this, a non-redundant set of representative structures for the UniProt canonical proteome was collated. Here, structures from the ZoomVar database were mapped in order of identity. Additional structures were only added to represent a protein if at least 50 % of the residues they covered were not mapped to a structure with higher identity. Protein core density was only calculated for those proteins with a single mapped structure. This was to avoid problems associated with determining the core density for multi-domain proteins. The number of C$\alpha$ contacts ($< 8$ Å) for each core residue [Q(SASA) $< 0.15$] was counted. The mean number of contacts for all core residues within a protein was used as a proxy for protein density, the assumption being that protein cores with greater density will have a higher number of C$\alpha$ contacts. Only protein cores which comprise of $> 4$ residues were analysed.

#### S1.5 Enrichment analysis of gene sets

##### S1.5.1 Gene set enrichment

Gene enrichment analyses were performed using Gene Set Enrichment Analysis (GSEA) [16], as implemented in the R fgsea package [37]. At the full-length protein level only sets with n $\geq$ 25 were considered. Due to incomplete structural coverage of the proteome, enrichment calculation for protein regions was possible for a smaller number of proteins. For consistency, all pathways analysed at the whole protein level were also analysed at the protein regions.

For GSEA on expression, abundance, stability and half-life data, they were performed as detailed above and in the main text Materials and Methods section, but using these statistics as input to the algorithm:

| | *Metrics used for GSEA* |
|---|---|
| **thermal stability** | Tm - mean(Tm) |
| **abundance** | log(ppm + 1) - mean(log(ppm + 1)) |
| **expression** | log(median(RPKM) + 1) - mean(log(median(RPKM) + 1)) |
| **half life** | log(hours) - mean(log(hours)) |

Table S1: Proteomics and transcriptomics-based metrics used as enrichment statistics for GSEA analysis.

##### S1.5.2 CATH architecture enrichment analysis

Pfam domains [29] were assigned CATH domains [38] using the per-residue mapping of structures to domains, from both data sets, available from the SIFTs resource [39]. A minimum of 50 residues, which mapped to both a particular CATH domain and a particular Pfam domain were required to assign a Pfam domain to a CATH domain. This threshold was used in order to prevent spurious assignments. If a Pfam domain appeared to map to more than one CATH domain, the majority vote, from the residue level mapping, was used. Using these assignments Pfam domains were mapped to CATH architectures, guided by the CATH hierarchy, to create "domain sets" for each architecture.

Architecture enrichment analysis was performed as for gene set enrichment analysis, as described in the main text (Section S1.5.1), however here the VESs calculated from full-length domain-types, as well as the constituent structural regions (for each Pfam domain type) were used as the enrichment statistics. Additionally, as we were interested in the enrichment of individual architectures only "domain" sets of size n $\geq$ 25 were considered at all levels.

#### S1.6 Database implementation

All data, including per-residue mappings, were stored in the ZoomVar MySQL database [40]. A web interface and REST (Representational State Transfer) architecture was implemented, using the Django framework [41], to allow users to query the ZoomVar database. It is available at http://fraternalilab.kcl.

ac.uk/ZoomVar. The database is designed for programmatic access to structurally annotate user-specified variants of interest.

### S2 Supplementary Results

#### S2.1 Population and disease-associated variants have different topological structural network properties

Here we ask whether variants from different datasets have distinct topological properties according to their structural localisation. We define these topological properties by representing protein structures at networks, in which nodes consist of C$\alpha$'s, and those C$\alpha$'s within 10 Å of one another are connected by edges (see Section S1.3 for more details. Due to this representation, we are able to calculate topological network properties of the residues to which variants localise. Such properties give insight into the connectivity and neighbourhood of the affected residues.

The results of the analysis are depicted in Figure S1. Given that we find protein cores to be enriched in ClinVar variants, it is perhaps unsurprising that we find these variants to localise to residues with a significantly higher degree (are more highly connected), than do variants from the other datasets (pairwise Mann-Whitney test, see Section S4.4 for values). We also find that disease-associated variants show significantly higher values for several other centrality measures. Interestingly, the most significant difference between datasets (Kruskal-Wallis test, see Section S4.4), after degree, is betweenness centrality. This metric can be seen to highlight bottlenecks within the network, as it is a measure of the fraction of shortest paths which pass through a node. This suggests that disease-associated variants may target residues which play an important role in communication through the structural network. Conversely, ClinVar variants localise to residues with only slightly higher median degree centrality (the fraction of connected nodes) than population variants, and slightly lower median degree centrality than COSMIC driver variants. This suggests that disease-associated variants localise to residues with a higher degree as they occur in proteins in which all residues are more highly connected. Although we see significant differences between the datasets, it is clear there is a large overlap in their distributions of topological network features (see Figure S1).

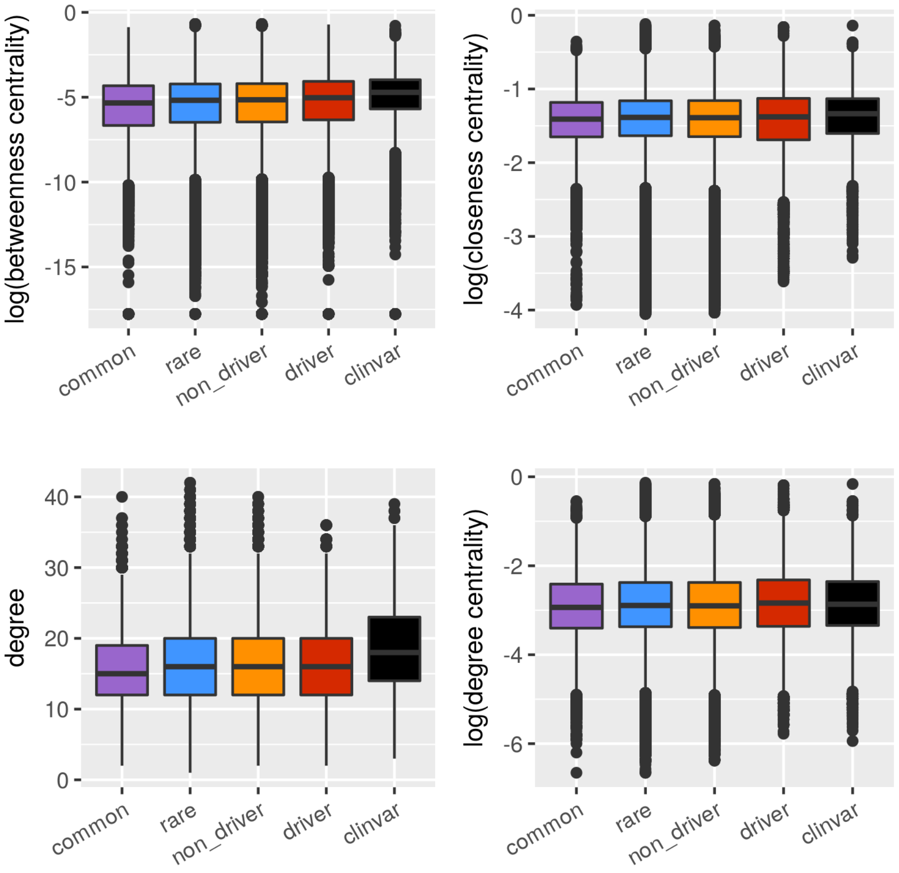


Figure S1: C$\alpha$ structural network topological features of variants. Proteins are transformed into networks based on positions of C$\alpha$ carbons. Network topological features for the gnomAD common and rare, Cosmic and ClinVar datasets are compared. See Section S4.4 for associated statistics.

#### S2.2 The localisation of variants to CATH domain architectures

We studied whether disease-associated missense variants would localise preferentially to domains with specific architectures. To accomplish this we made use of the CATH protein domain classification system [38] and focussed on the architectural level, which groups domains with similar secondary structural orientations, thereby capturing tertiary structural features. We mapped Pfam domain definitions to those used by CATH, and created domain sets (analogous to gene sets) for each CATH architecture. Enrichment was calculated at both the domain-type level (i.e. localisation of missense variants to a domain-type, for example fibronectin type-III (Fn3), in comparison to localisation of missense variants to all other domain types, see Figure 2), and at the domain-type region level (i.e. localisation of missense variants to core residues within a domain-type, for example all Fn3 core residues, in comparison to the localisation of missense variants to all other residues within a domain-type).

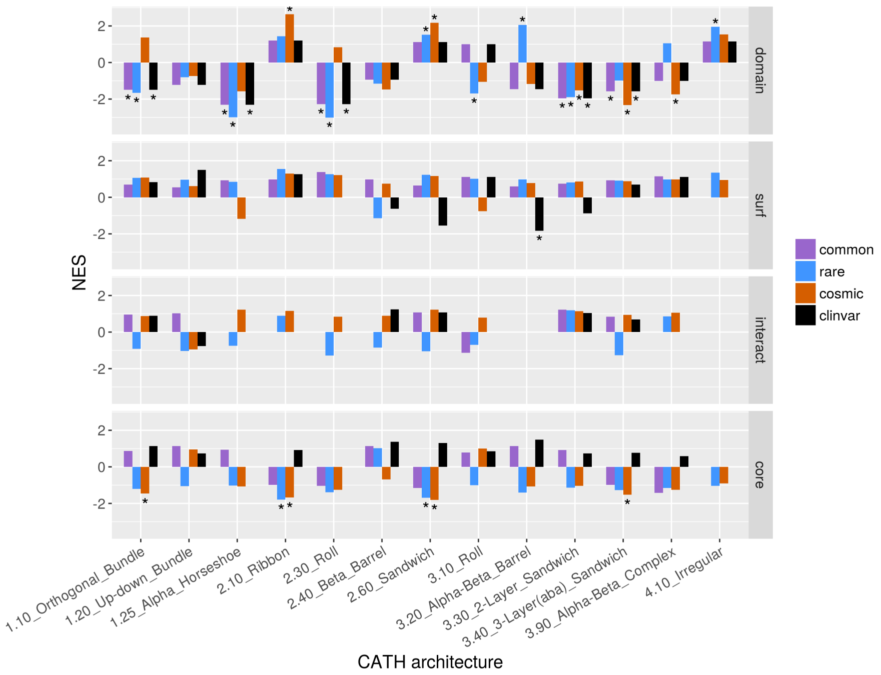


Figure S2: The enrichment CATH architectures in Pfam domains according to missense variant enrichment. Results are depicted at the domain level and the domain region level. * indicates q-value $< 0.05$.

As depicted in Figure S2, the results show that, at the whole domain level, the data sets show similar trends in variant localisation. A number of architectures, such as the Alpha Horseshoe architecture, show depletion of variants in all data sets, in contrast to other architectures, for example the Beta Sandwich and Irregular architectures, which show enrichment in all data sets. Few architectures, such as the Alpha-Beta Barrel, which is enriched in the gnomAD rare data but depleted for the other data sets, show markedly different patterns of enrichment at the domain architecture level. At the protein region level, the picture diversifies with the ClinVar data generally showing enrichment in architecture cores, although not significantly, in contrast to the other data sets which are more frequently depleted of variants in this region. Interestingly, this trend is particularly marked for the Alpha-Beta Barrel architecture. Thus, although gnomAD rare variants are enriched in this architecture, it is clear that very few of these localise to this architecture's core.

#### S2.3 Case study - DNA-binding proteins

This case study focusses on patterns of variant enrichment observed for DNA-binding proteins: a wealth of analyses have established the fundamental role of structural properties in the interaction of these proteins with DNA [42, 43, 44].

We considered a list of DNA-binding domains (DBD) curated in the literature [20], and compared their whole-domain and region VESs. Figure S3 shows that DBDs vary considerably in their targeting by disease-associated and population variants (stacked bar-graphs in Figure S3). Visualisation of the whole-domain level enrichment statistics shows that some DBDs (e.g. Forkhead, Homeobox) are only enriched in COSMIC and ClinVar variants, while zf-H2C2_2 domains, which are numerous in zinc finger (ZNF) proteins, appear to be enriched only in rare variants. Other DBDs, e.g. Myb_DNA_binding, appear depleted in all types of variants. Different regional enrichment patterns are also observed: ClinVar mutations are typically enriched in the core but devoid at the surface of these domains, whereas in the COSMIC data, as well as the two nominally healthy datasets derived from the gnomAD database, variants which localise to the surface are more common (Figure S3).

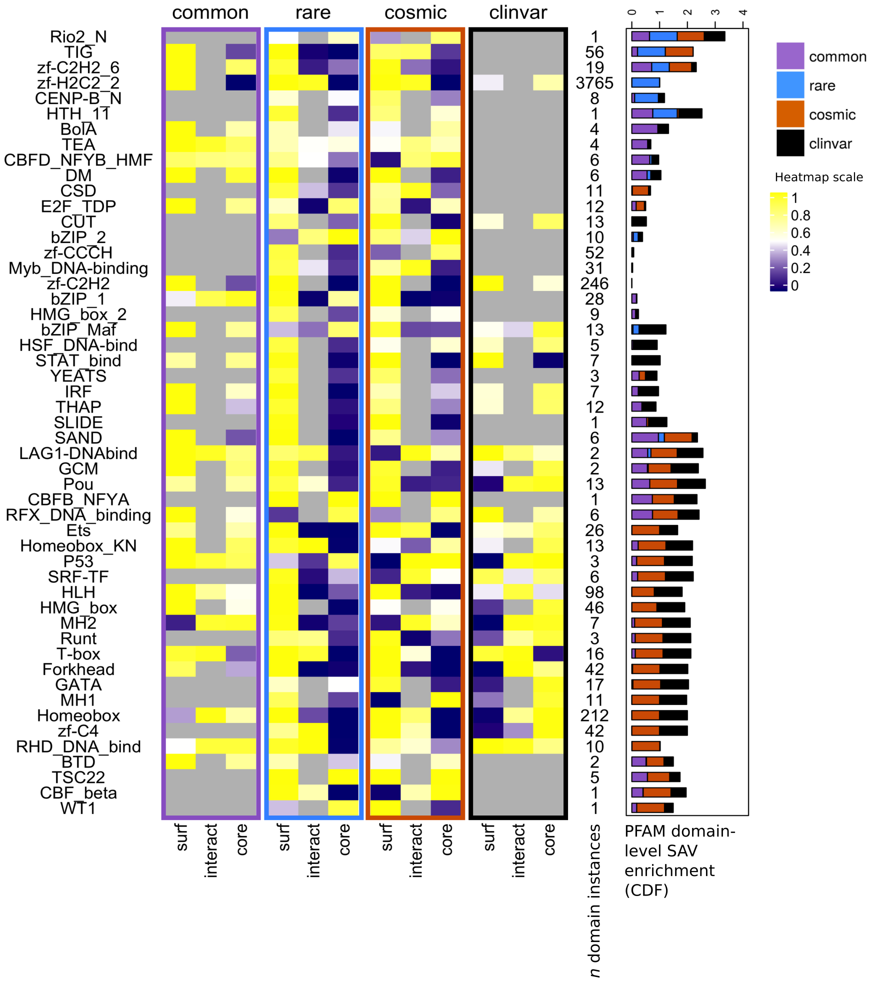


Figure S3: Landscape of variant enrichment in DNA-binding domains (DBD). Here all DBDs, as curated in [20], with structural coverage are considered, and their domain-region level enrichment is depicted. Each row corresponds to a Pfam domain type. Region (surface, interface and core) enrichments are shown in the heat maps for variants from the gnomAD common, gnomAD rare, COSMIC and ClinVar datasets. Grey cells correspond to the absence of interaction site mappings (for interface data), or the absence of mutations in our dataset. Each row is annotated with the number of domain instances of each type present within our data. A stacked bar graph shows the enrichment at the whole protein level for each data set.

### S3 Supplementary Figures

*From the next page onwards.*

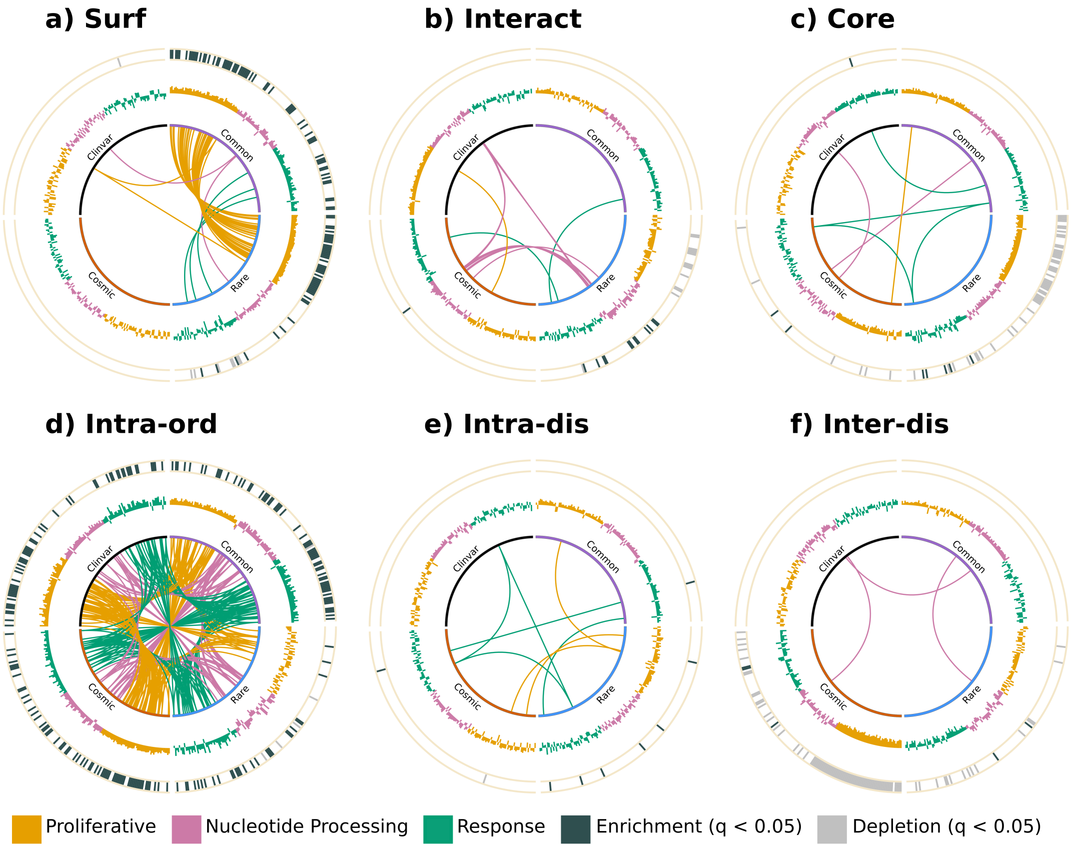


Figure S4: Functional enrichment for each variant dataset, at different region levels (a-f), visualised on a Circos plot. The normalised enrichment score for each pathway is plotted as a bar graph (the further from the centre, the more positive) in the middle layer of the plot. In the outermost layer of the plot, significant enrichment (dark grey) or depletion (light grey) of a pathway ($q < 0.05$) is depicted.

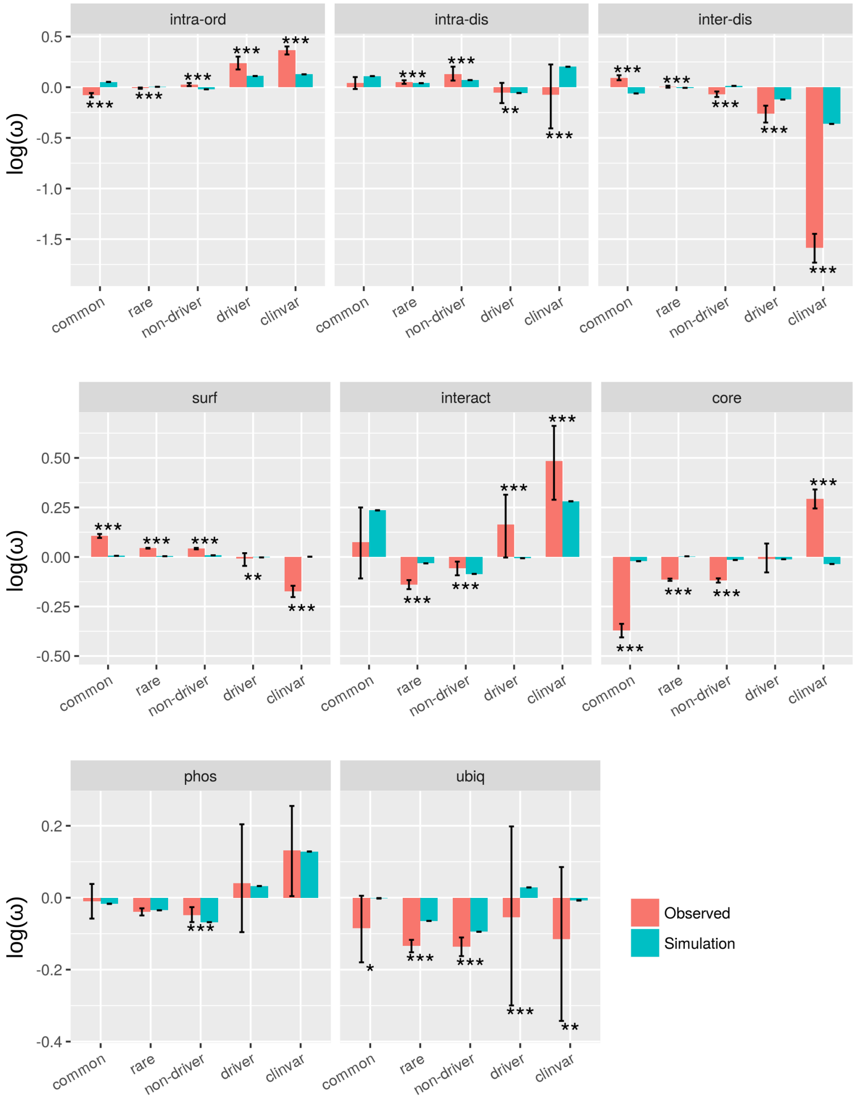


Figure S5: The density of mutations in different protein regions. Density ($\omega$) values were taken logarithm such that negative values indicate depletion while positive values indicate enrichment. Observed densities (pink), and densities derived from simulated null distributions (turquoise) are shown. Error bars depict 95% confidence intervals, for observed densities these were obtained by bootstrapping. Significance was calculated by comparison of observed values to simulated null missense variant distributions (significance level indicated by: * q-value $< 0.05$, ** q-value $< 0.001$, *** q-value $< 0.0001$).

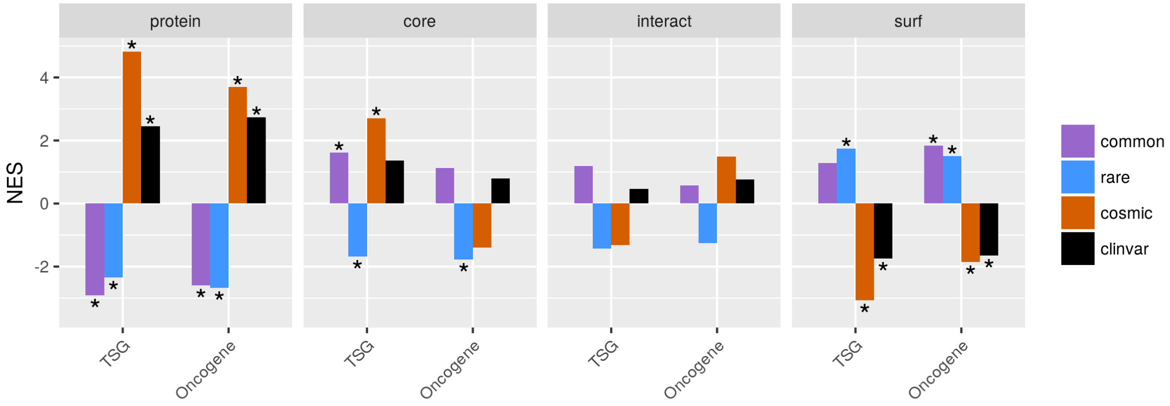


Figure S6: The enrichment of tumour suppressor and oncogene gene sets in the gnomAD common, gnomAD rare, COSMIC and ClinVar data. Enrichment calculated using the GSEA algorithm using the Variant Enrichment Score as the enrichment statistic. Tumour suppressor and oncogene sets were defined as described in the Methods.

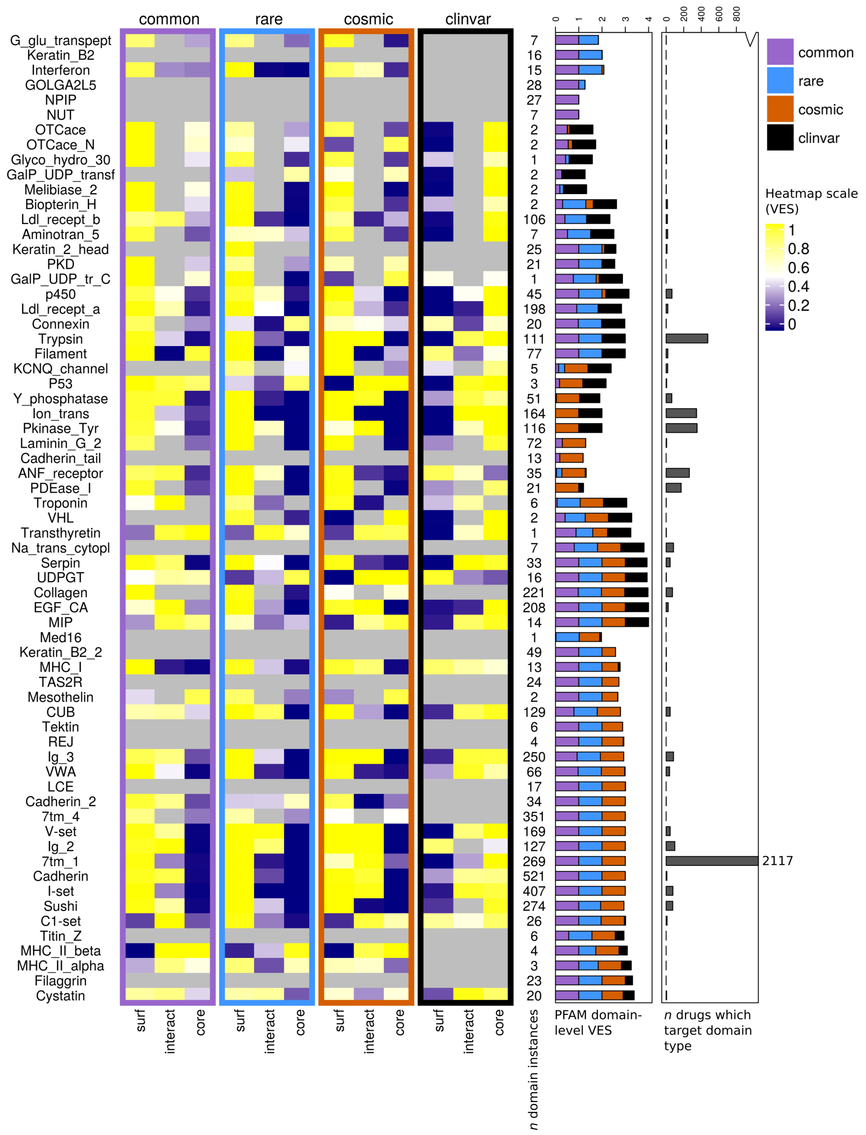


Figure S7: The landscape of variant enrichment over a list of domain-types most enriched in pathogenic and population variants. Here the union of the top 20 most enriched domain-types for each datset is depicted: each role corresponds to a Pfam domain-type. The plots are arranged in the same way as in Figure S3, showing from left to right the following visualisations: first, on the left, structural region (surface, interface and core) enrichments are shown in the heat maps for variants from the gnomAD common, gnomAD rare, COSMIC and ClinVar datasets. Grey cells correspond to the absence of interaction site mappings (for interface data), or the absence of mutations in our dataset. Each row is annotated with the number of domain instances of each type present within our data. Second, a stacked bar graph shows the enrichment at the whole protein level for each data set. On the rightmost bar graph, the numbers of drugs known to target proteins containing each domain-type are shown. Note the cut numeric axis; the number of drugs which target the only outlier, the 7tm_1 domain, is noted on the plot.

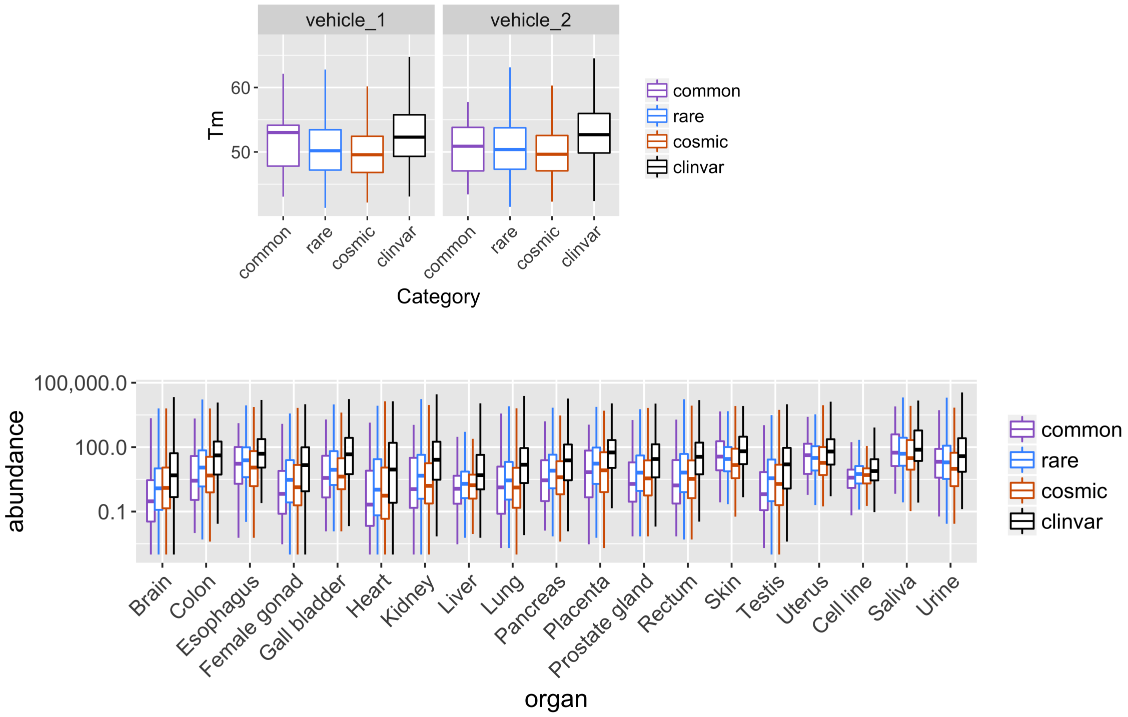


Figure S8: Stability and abundance of proteins enriched in variants in each dataset. For each dataset, here we considered the whole-protein level statistics and extracted proteins which were enriched (q-value $< 0.05$) in variants, and plotted distributions of their stability (melting temperature, or Tm [$^o$C], top panel) and abundance (ppm, bottom panel). Note these values correspond to the stability/abundance measurements of the wild-type protein, i.e. values at the ‘base line’ without any missense changes. Note again for the Tm data, measurements were done on two replicates (“vehicles”). Both datasets were considered here and were shown in two separate panels (top).

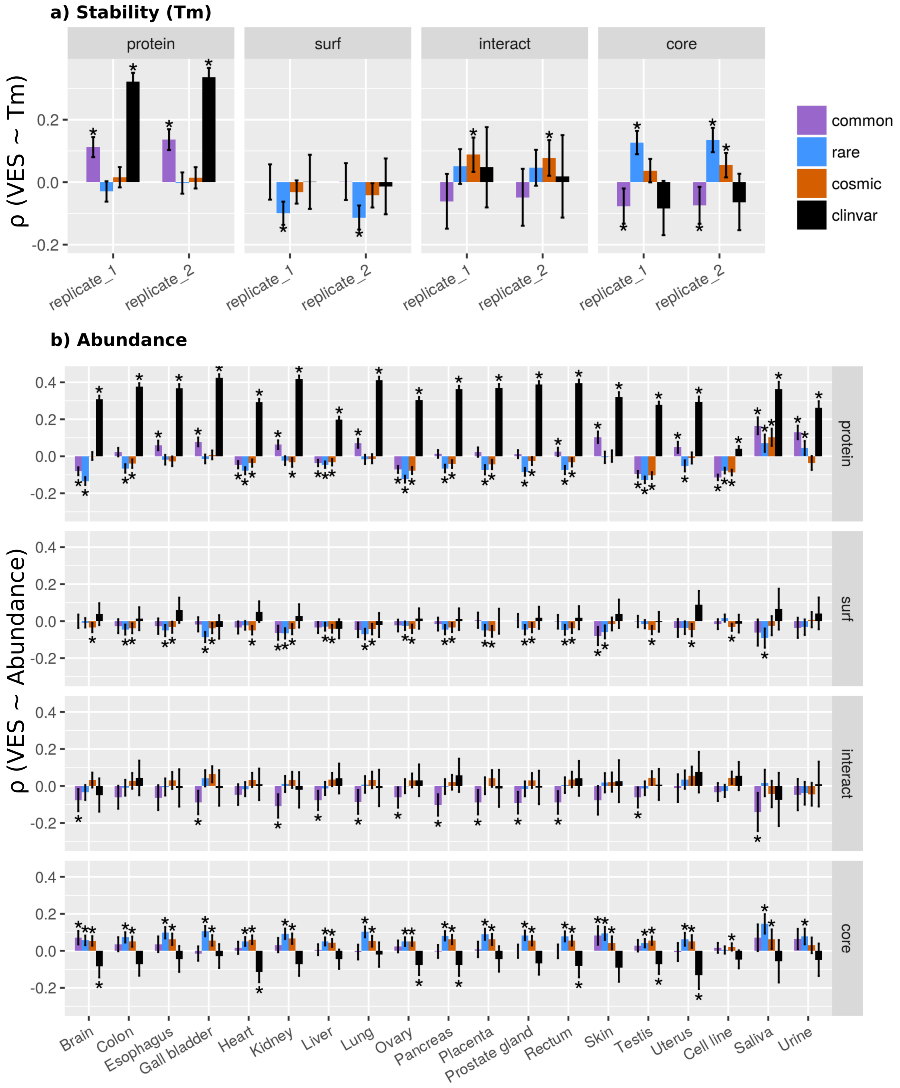


Figure S9: Spearman correlations for missense variant enrichment (quantified as VESs) with (a) protein thermal stability (Tm, in $^o$C) and (b) protein abundance (ppm). Correlations with VESs calculated at the full-length "protein" level, and the surface (`surf`), core and interacting interface (`interact`). Error bars indicate 95% confidence intervals. * indicates q-value $<$ 0.05. Selected organs/cell types are shown in the main text.

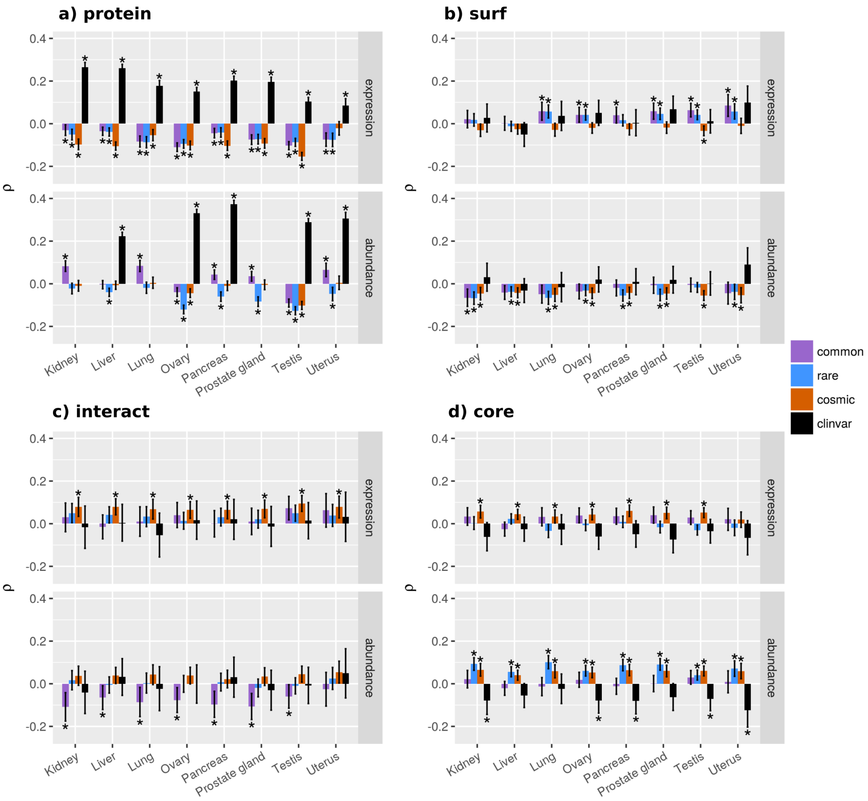


Figure S10: The enrichment of missense variants (VES) in comparison to protein abundance (ppm) and expression (median count) at the a) full-length protein level, b-d) surface (`surf`), core and interacting interface (`interact`). Spearman correlations calculated using only those proteins present in both abundance and expression data. Error bars indicate 95 % confidence intervals. * indicates q-value $< 0.05$.

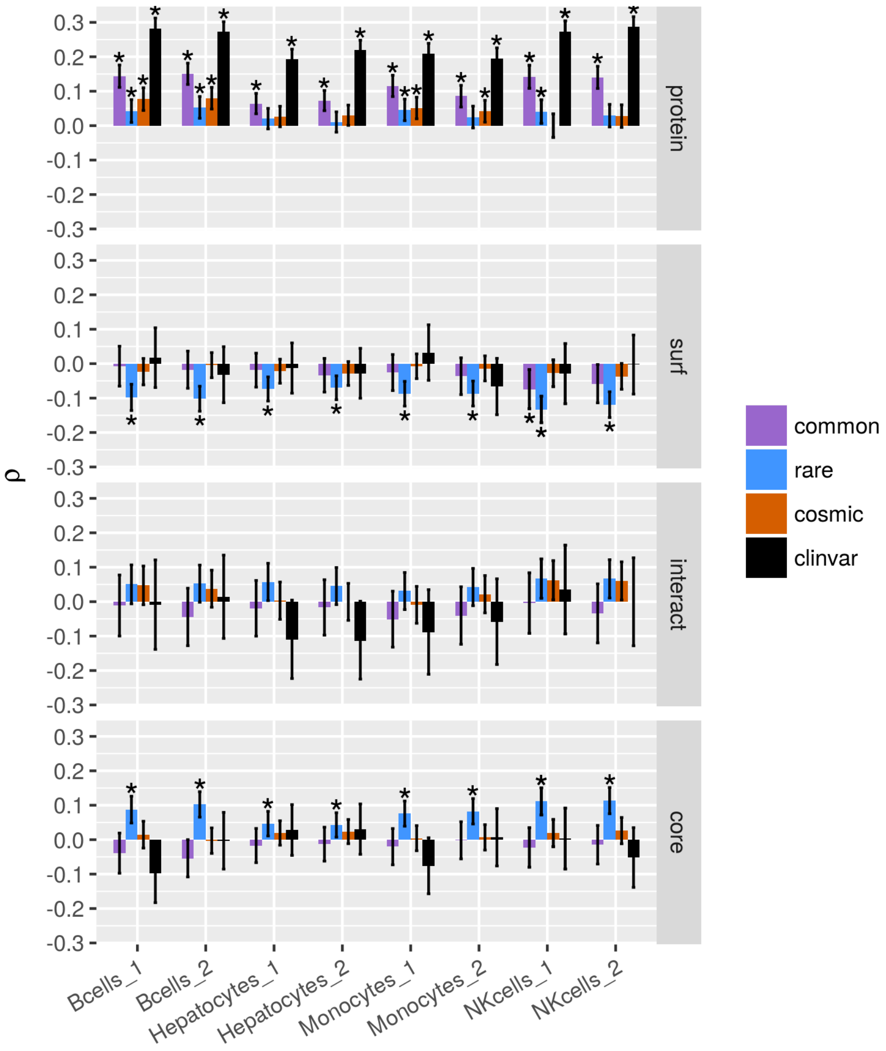


Figure S11: The Spearman correlation of the enrichment of missense variants (VES) with protein half life data (hours). Error bars indicate 95 % confidence intervals. * indicates q-value < 0.05.

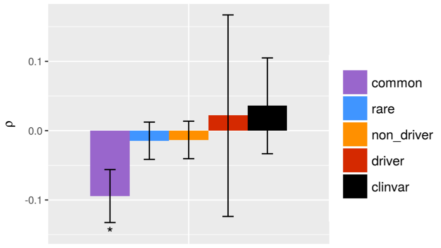


Figure S12: The Spearman correlation of the enrichment of missense variants in protein cores (VES) with a proxy for protein core density (see Methods). Error bars indicate 95 % confidence intervals. * indicates q-value $<$ 0.05.

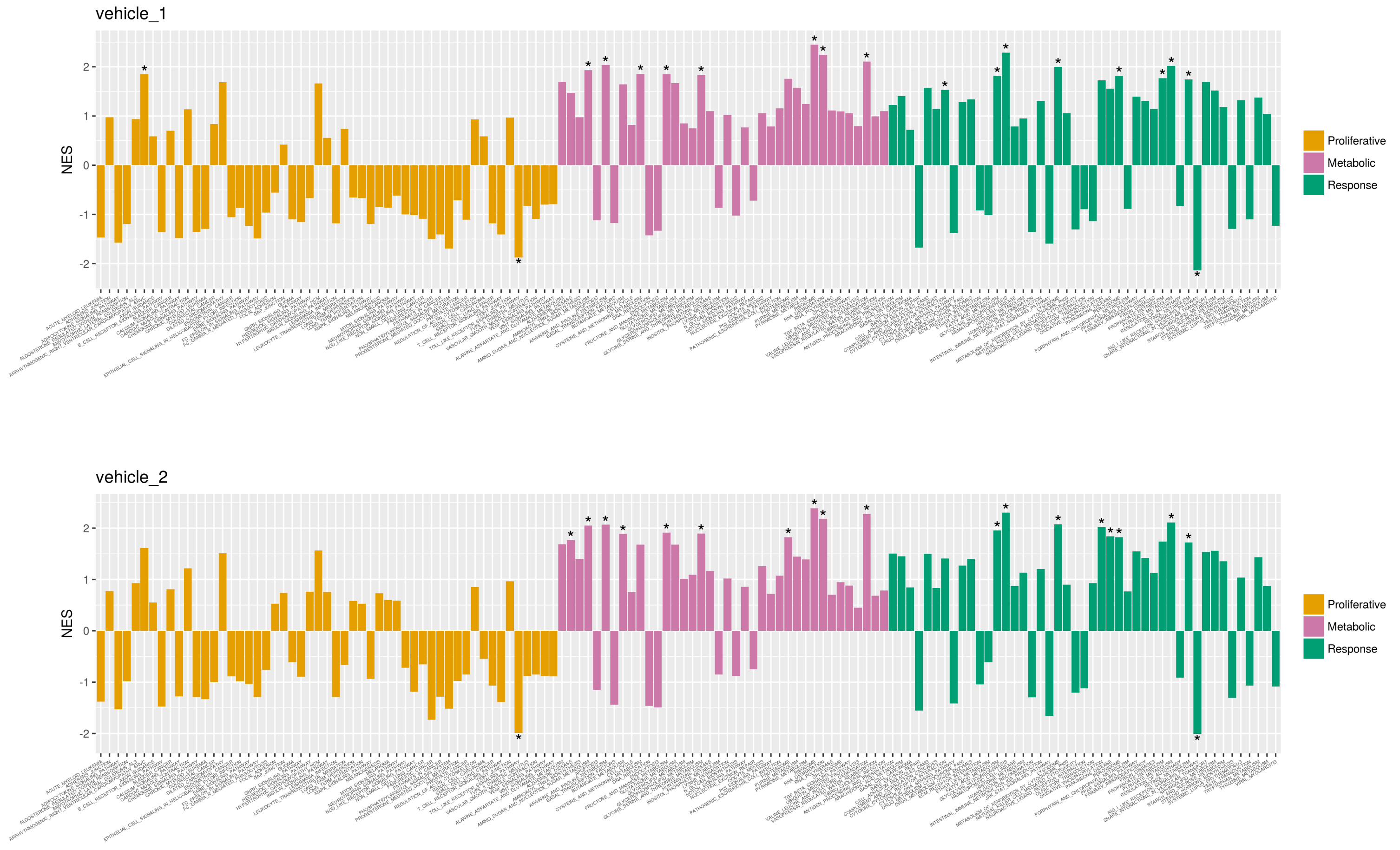


Figure S13: The functional enrichment of proteins in KEGG pathways according to thermal stability. Pathways have been mapped to the 3 clusters defined in the main text.

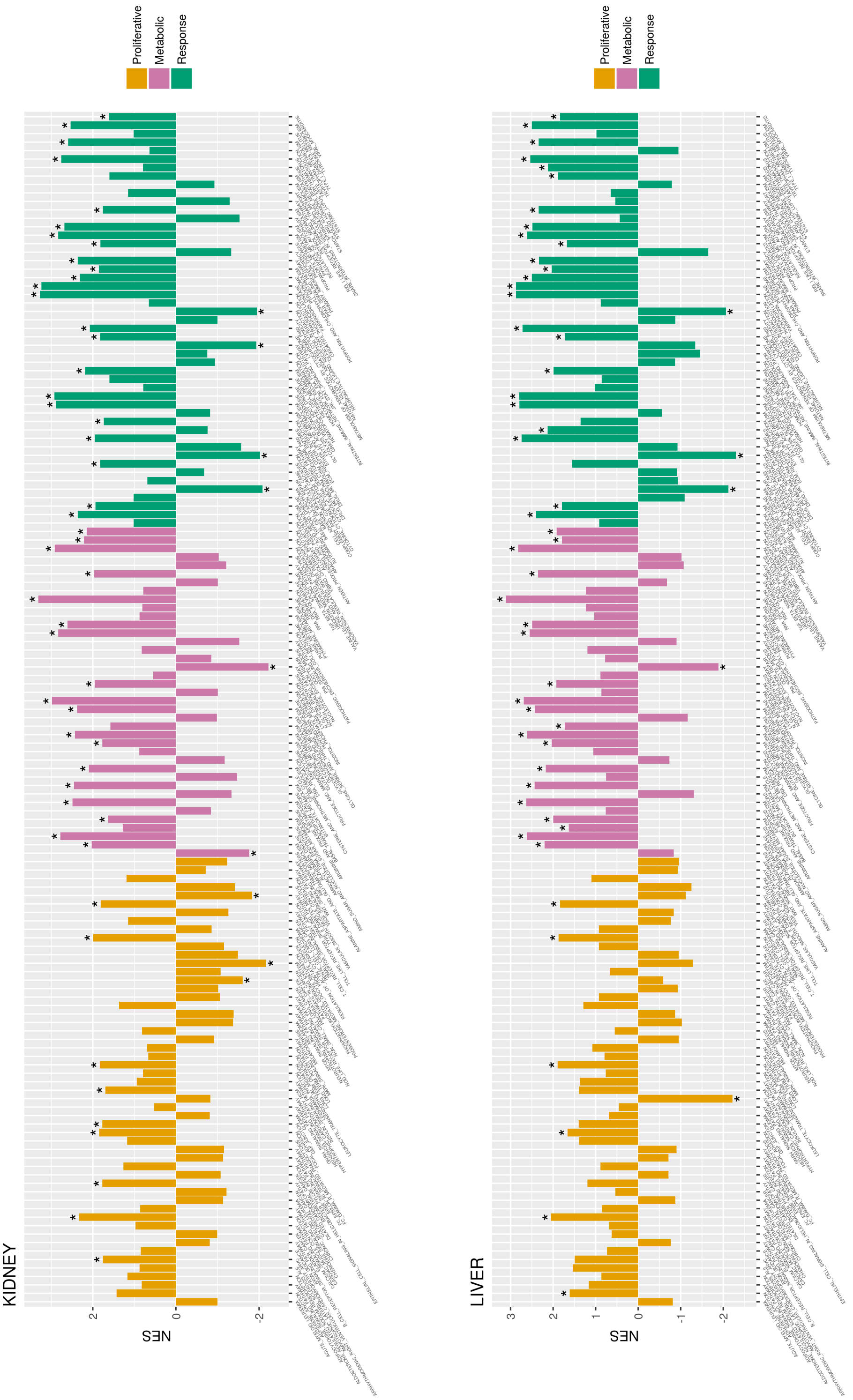
KIDNEY
LIVER
NES
Proliferative
Metabolic
Response

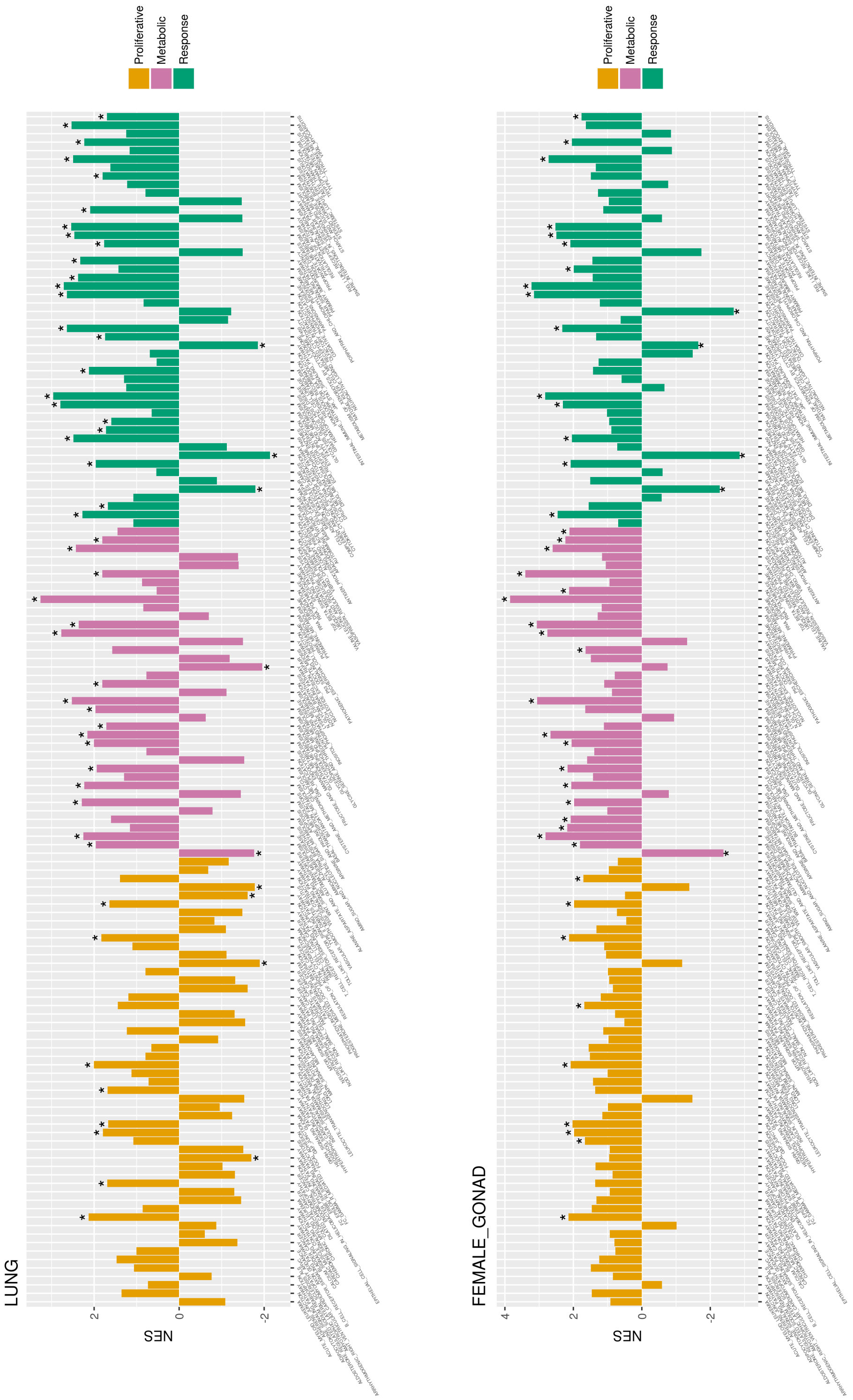

LUNG
FEMALE_GONAD
NES
Proliferative
Metabolic
Response

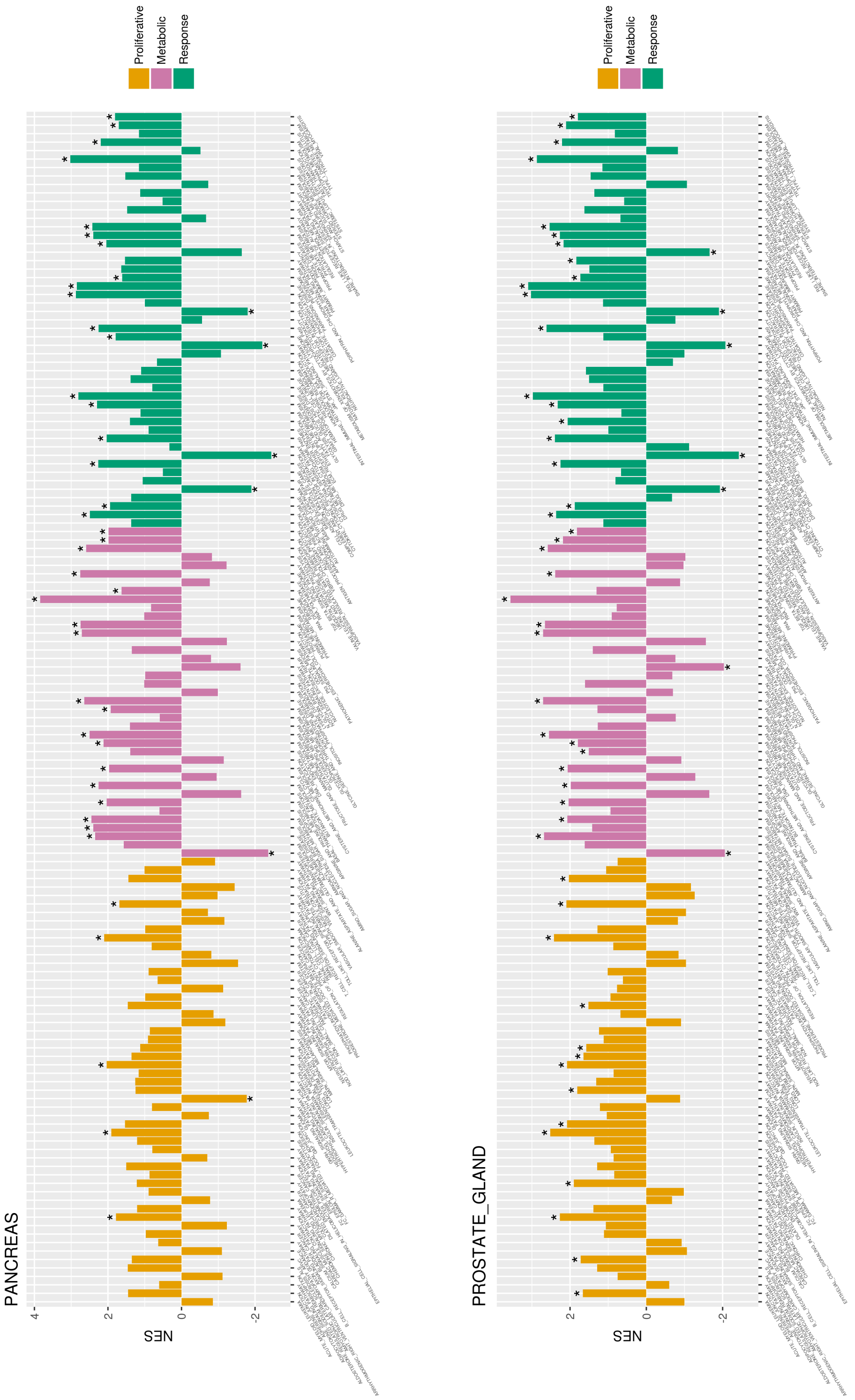
PANCREAS
PROSTATE_GLAND
NES
Proliferative
Metabolic
Response

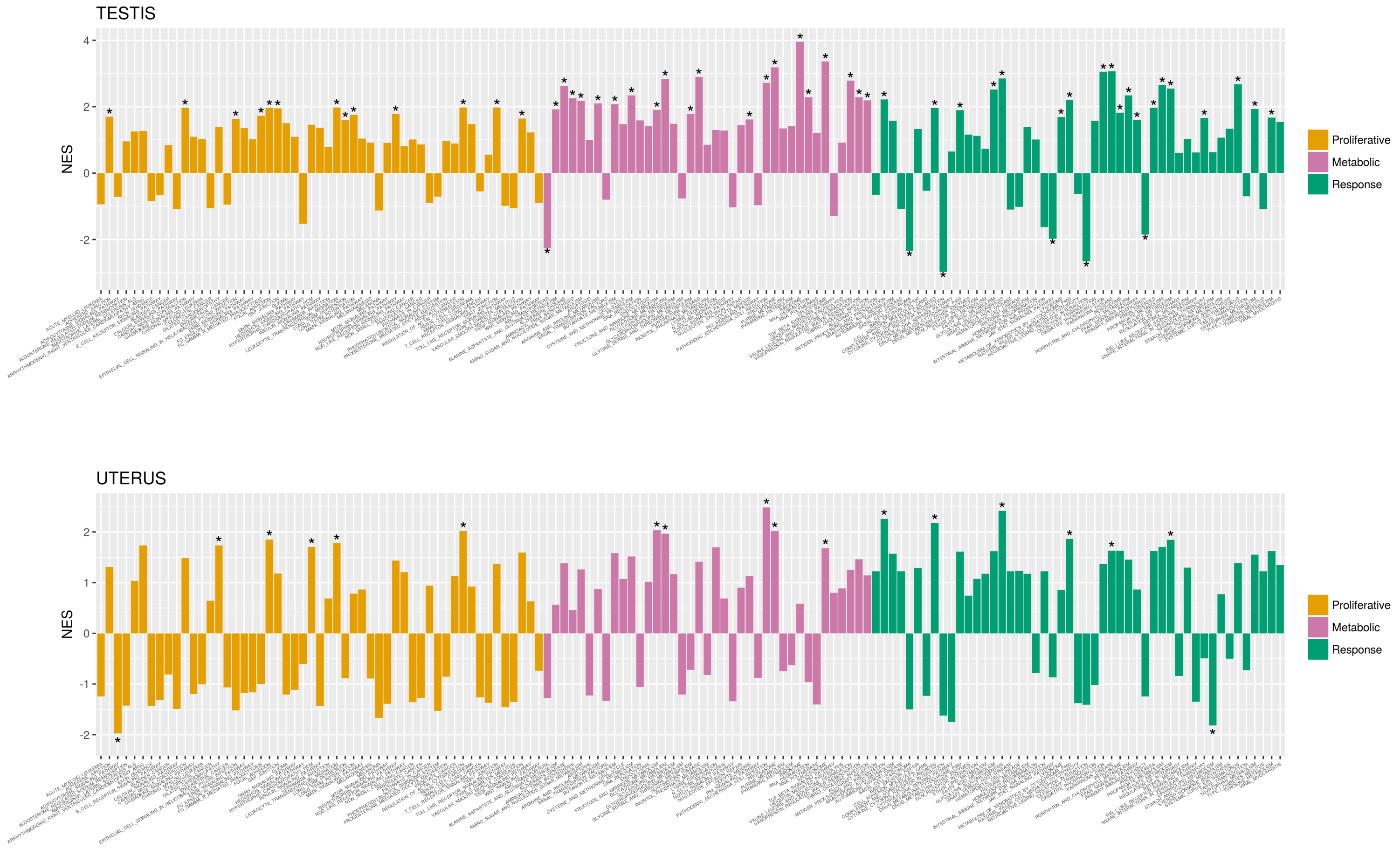


Figure S11: The functional enrichment of proteins in KEGG pathways according to protein abundance. Pathways have been mapped to the 3 clusters defined in the main text.

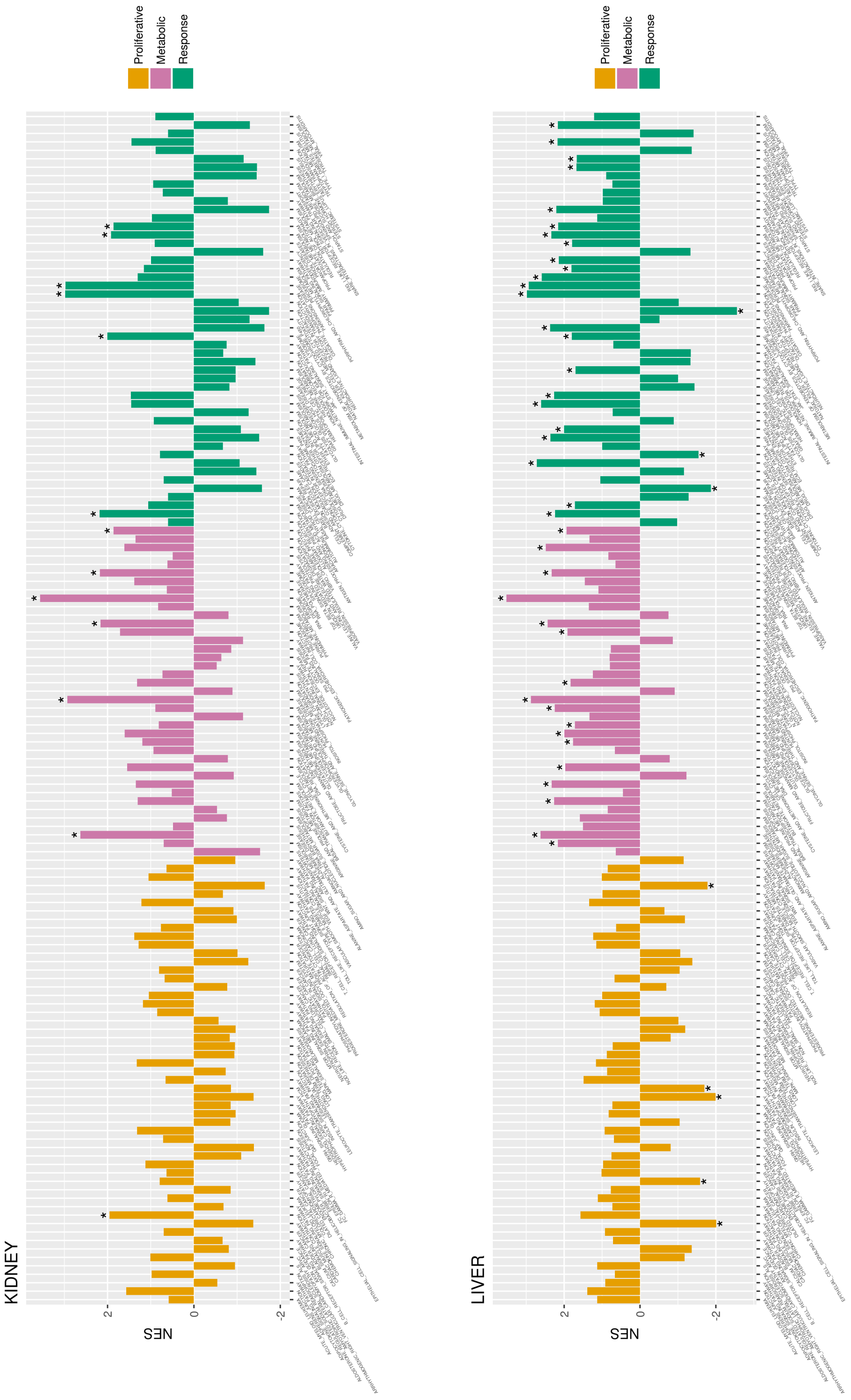
KIDNEY
LIVER
NES
Proliferative
Metabolic
Response

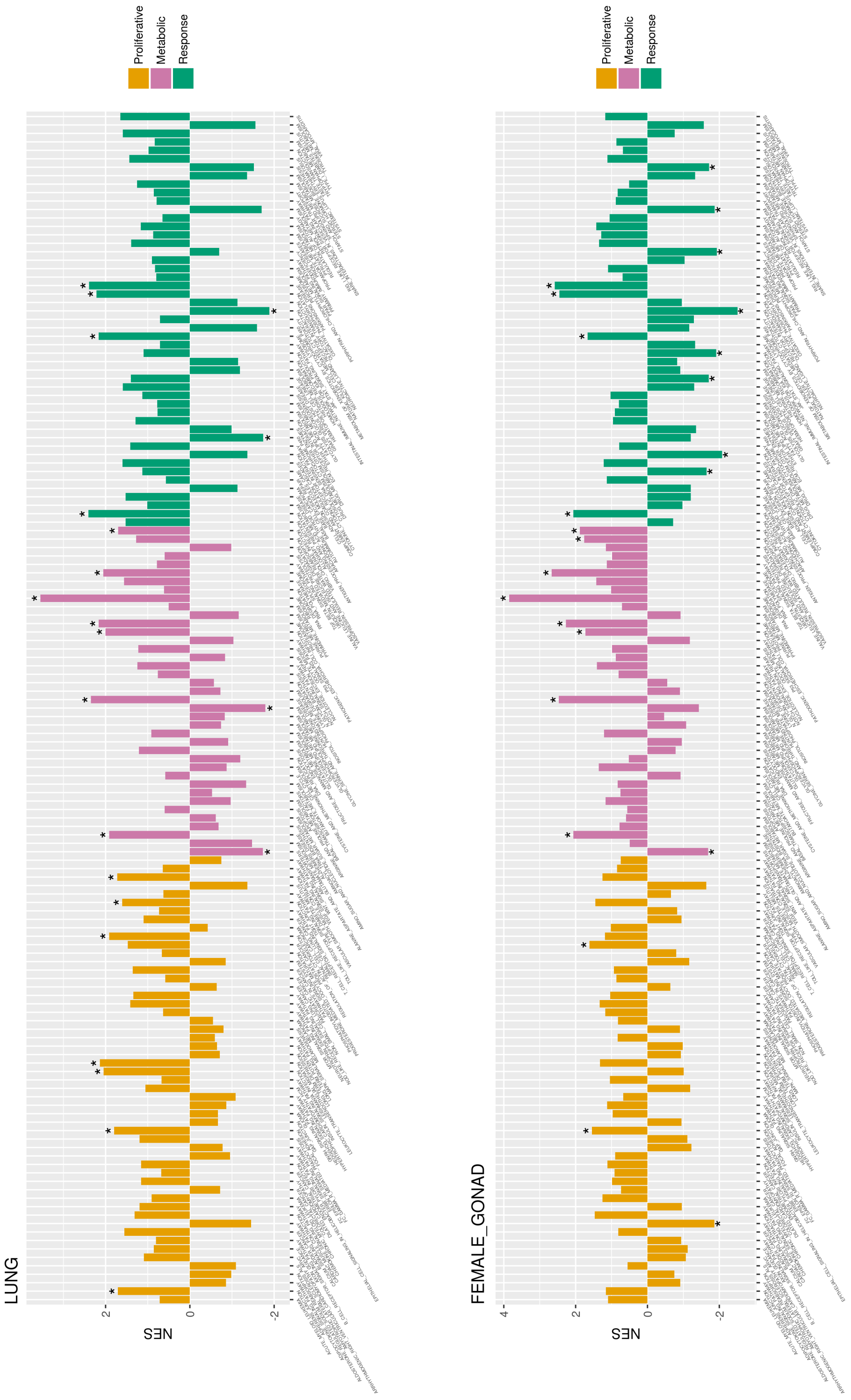
LUNG
FEMALE_GONAD
NES
Proliferative
Metabolic
Response

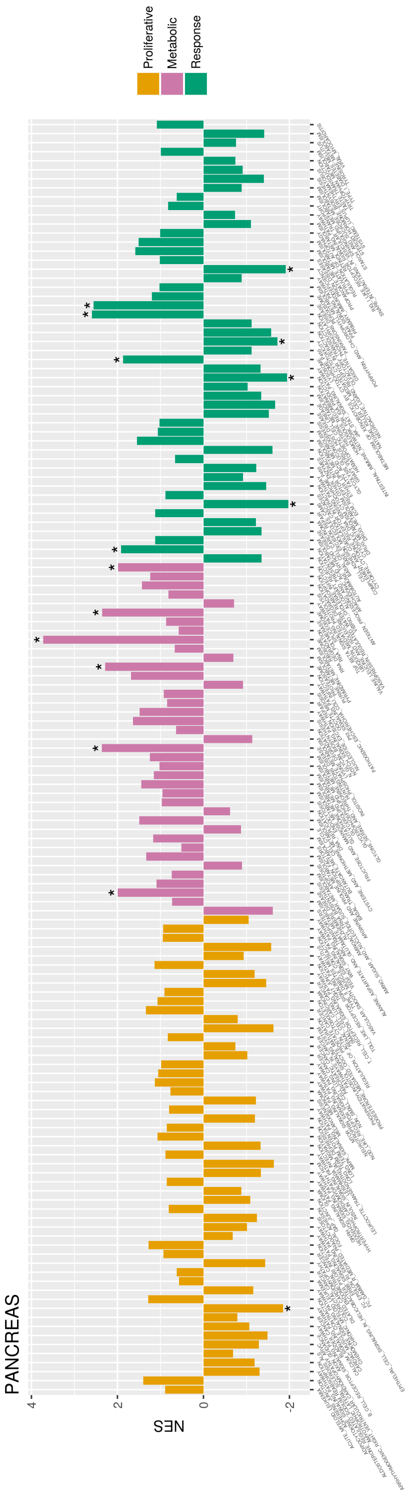
PANCREAS
NES
4
2
0
-2
Proliferative
Metabolic
Response

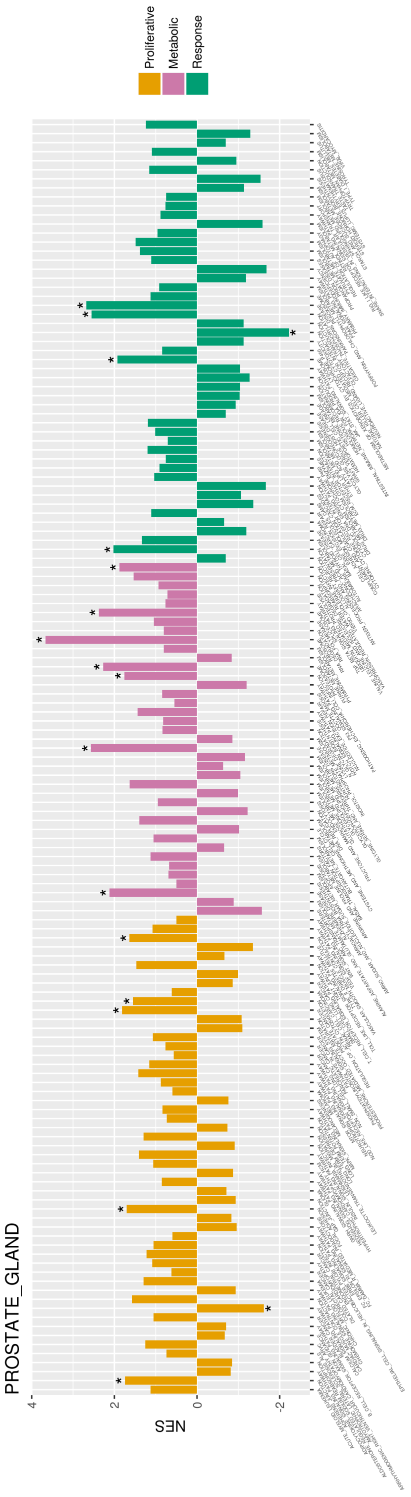
PROSTATE_GLAND
NES
4
2
0
-2
Proliferative
Metabolic
Response

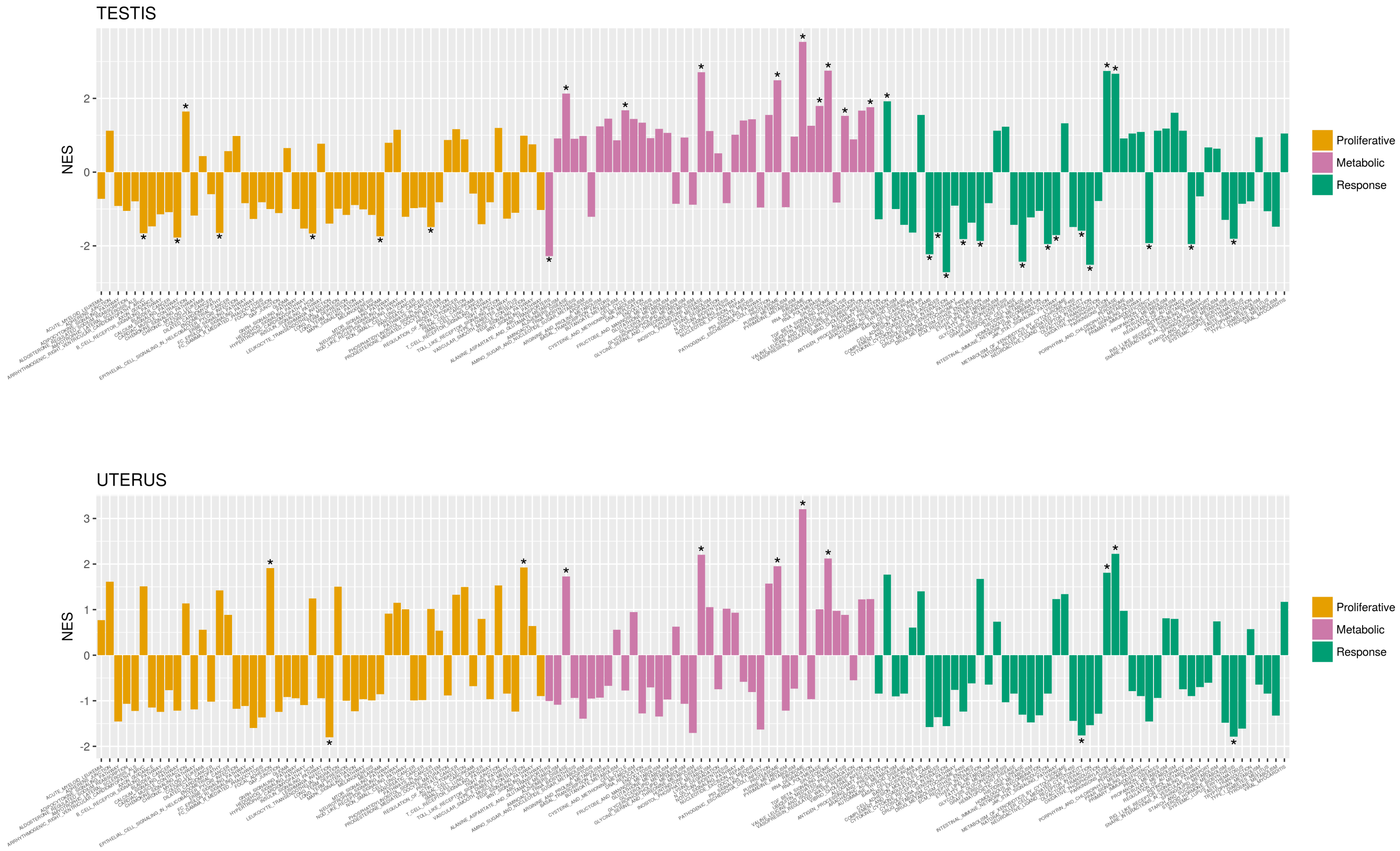


Figure S12: The functional enrichment of proteins in KEGG pathways according to transcript expression. Pathways have been mapped to the 3 clusters defined in the main text.

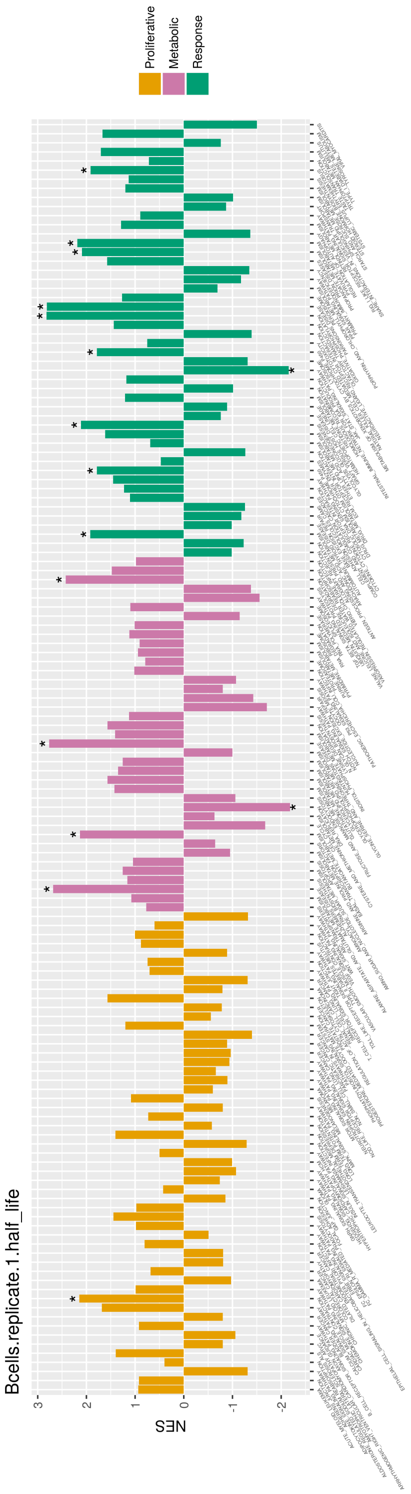
Bcells.replicate.1.half_life
NES
Proliferative
Metabolic
Response

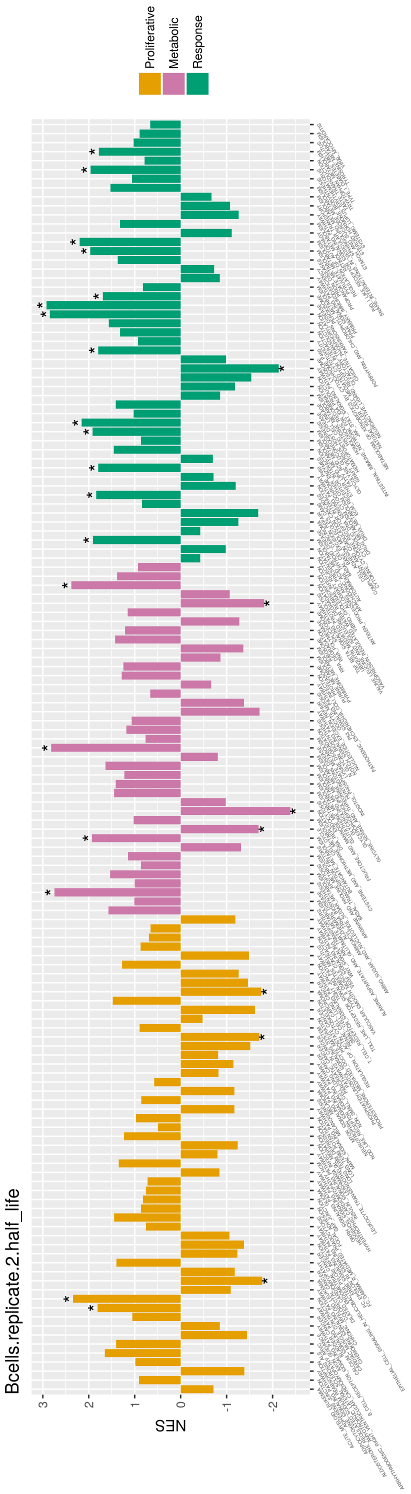
Bcells.replicate.2.half_life
NES
Proliferative
Metabolic
Response

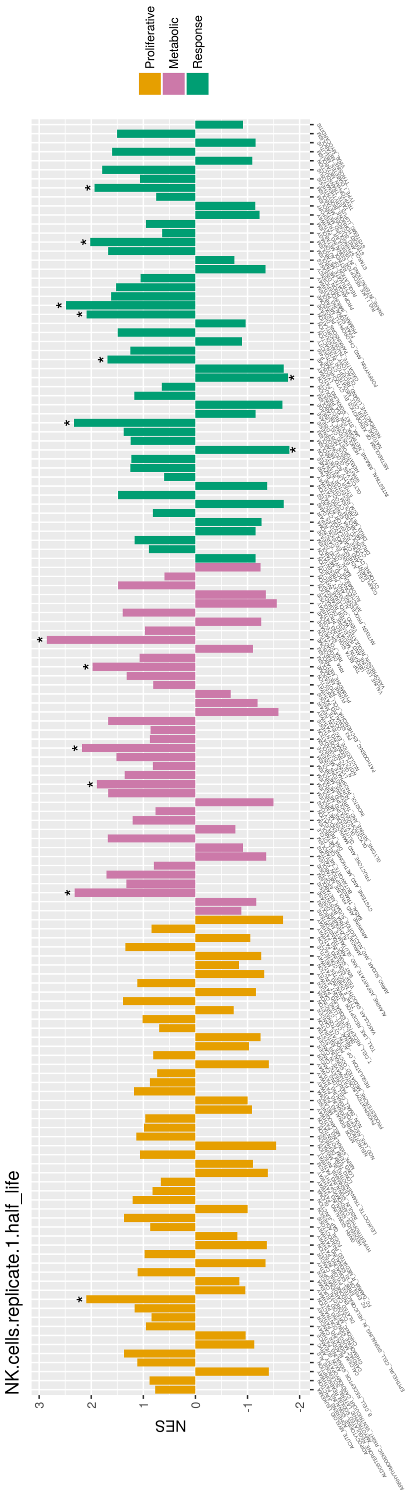
NK.cells.replicate.1.half_life
NES
Proliferative
Metabolic
Response

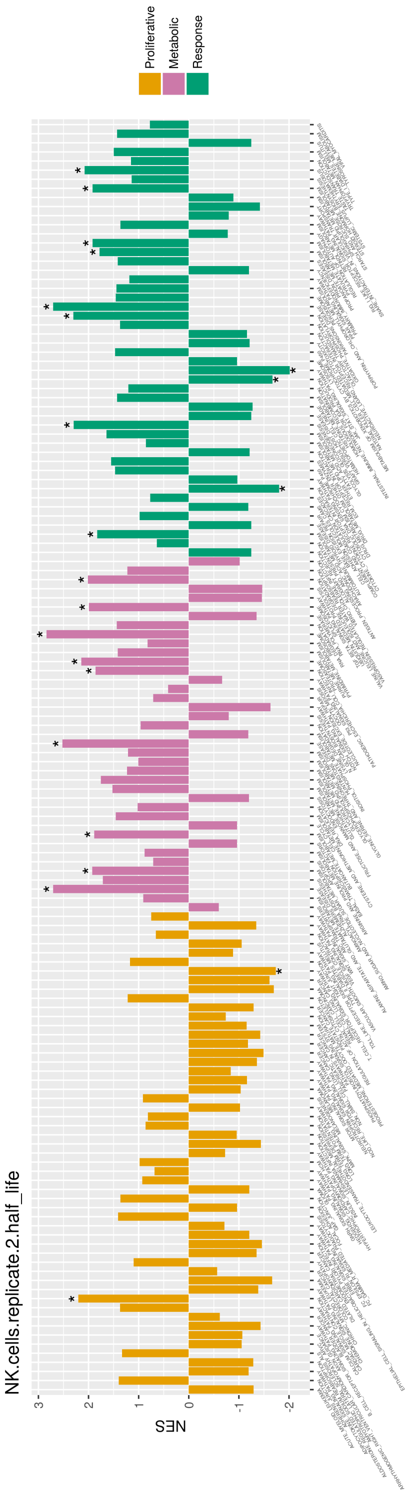
NK.cells.replicate.2.half_life
NES
Proliferative
Metabolic
Response

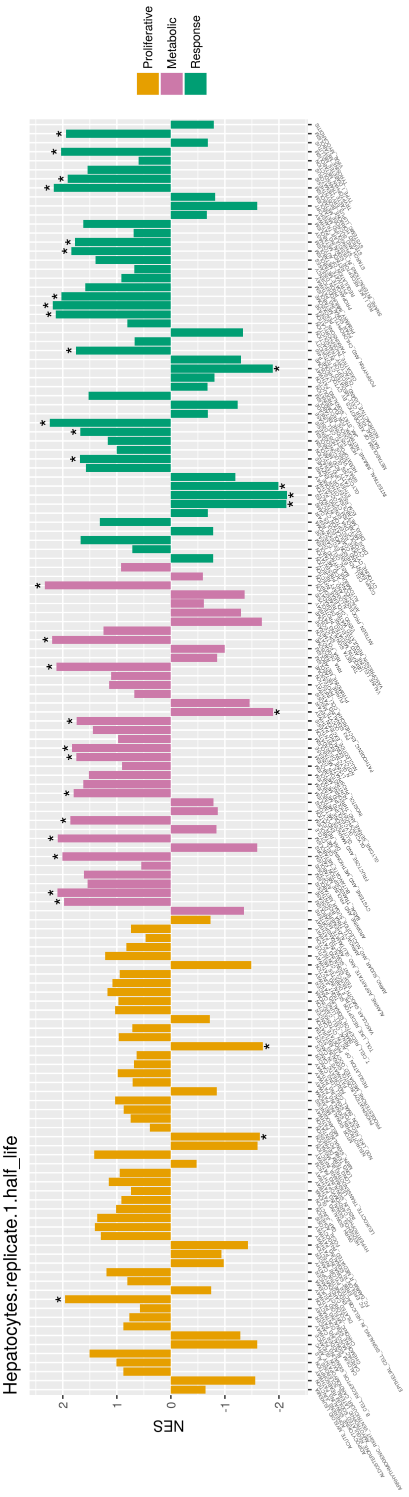
Hepatocytes.replicate.1.half_life
Proliferative
Metabolic
Response
NES
2
1
0
-1
-2

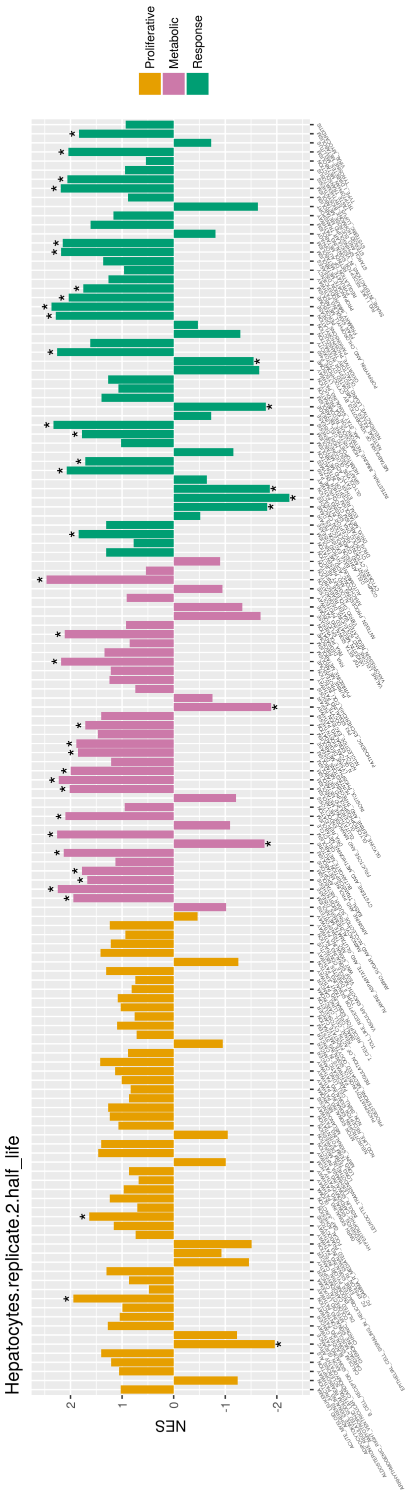
Hepatocytes.replicate.2.half_life
Proliferative
Metabolic
Response
NES
2
1
0
-1
-2

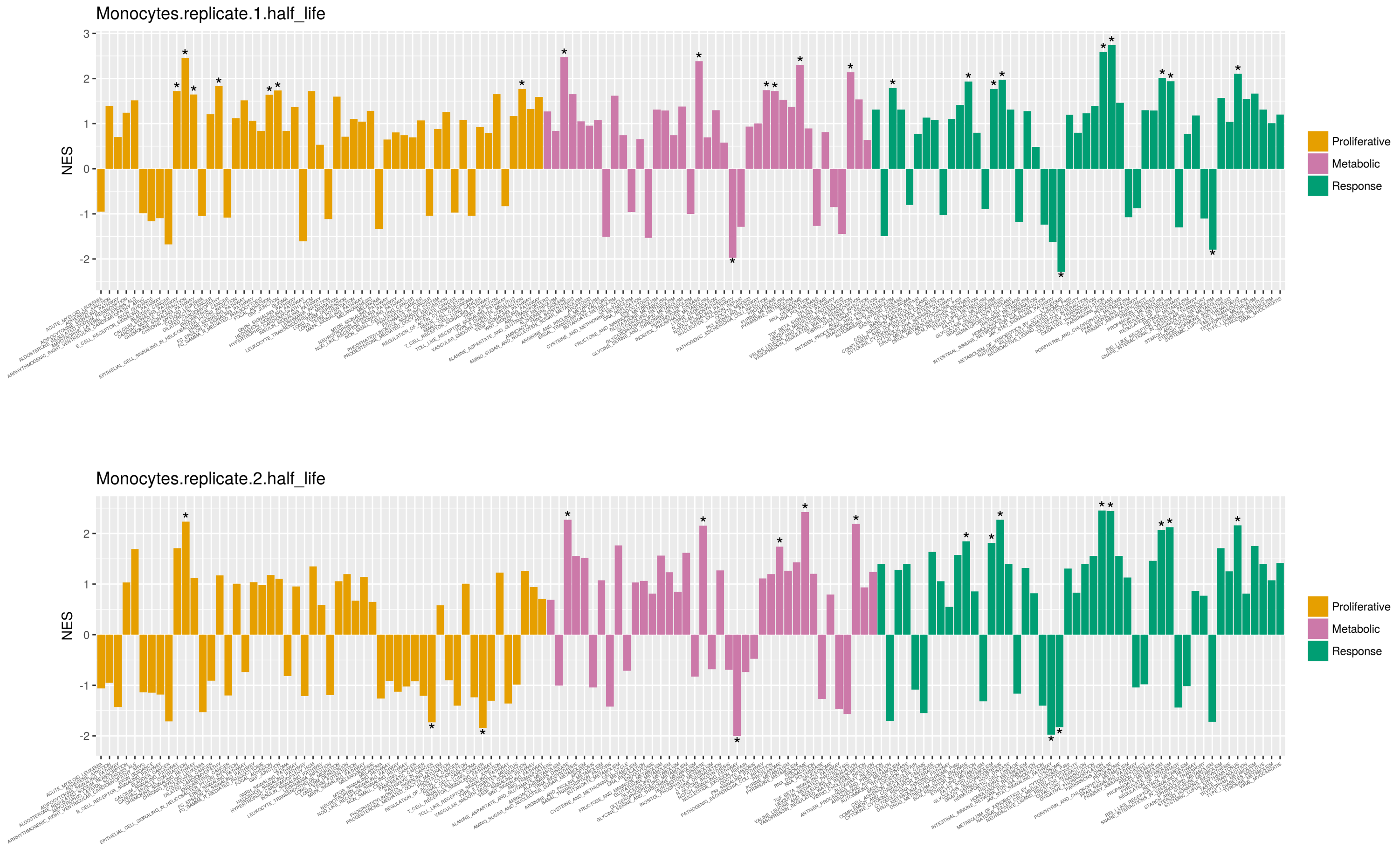


Figure S13: The functional enrichment of proteins in KEGG pathways according to protein half life. Pathways have been mapped to the 3 clusters defined in the main text.

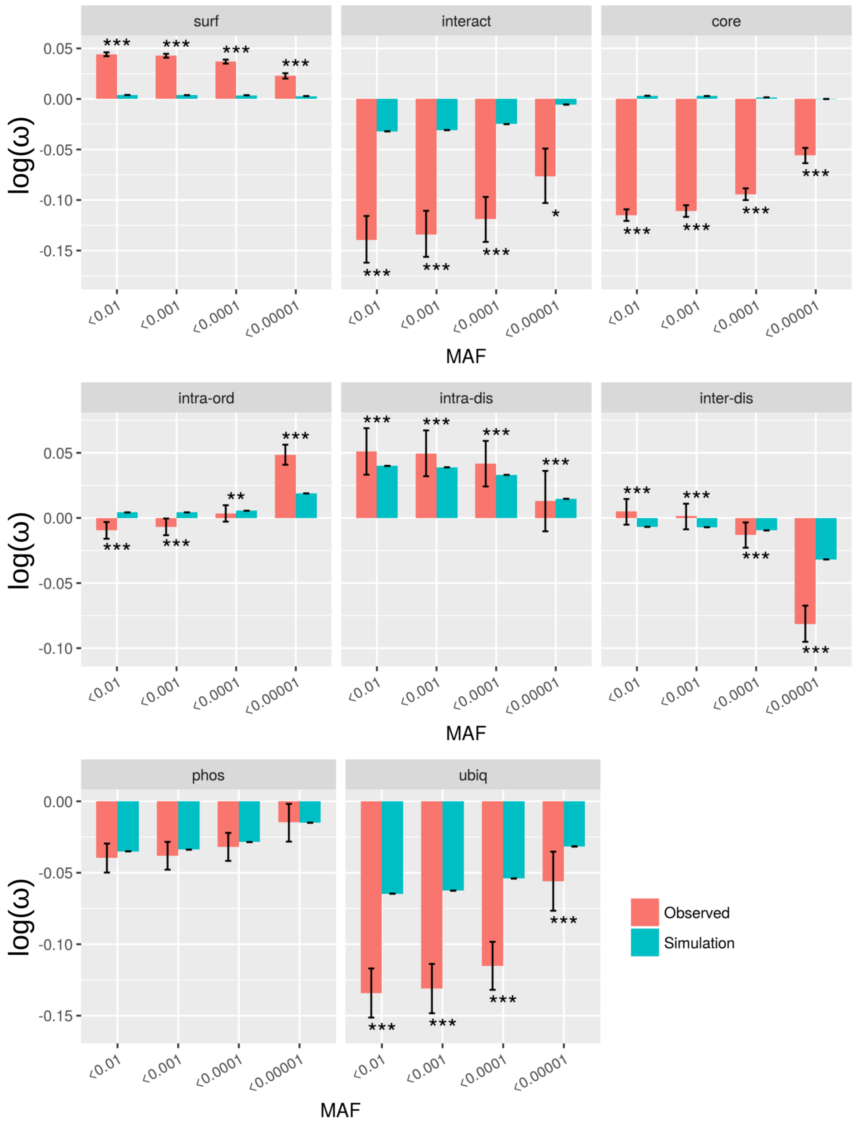


Figure S14: The density of rare mutations from the gnomAD data in different protein regions. Rare variants have been defined using different MAF cutoffs as shown in the x axis. Both observed densities (pink), and densities derived from simulated null distributions (turquoise) are shown. Density ($\omega$) values were taken logairthm such that negative values indicate depletion while positive values indicate enrichment. Error bars depict 95% confidence intervals, for observed densities these were obtained by bootstrapping. Significance was calculated by comparison of observed values to simulated null missense variant distributions (significance level indicated by: * q-value $<$ 0.05, ** q-value $<$ 0.001, *** q-value $<$ 0.0001).

#### S4 Supplementary Data

Supplementary data can be downloaded at http://fraternalilab.kcl.ac.uk/ZoomVar/downloads/

This consists of:

1. Details of the number of missense variants which localise to different protein regions in the gnomAD common and rare, COSMIC and Clinvar datasets.

2. A list of pathways annotated by functional cluster ("proliferation", "nucleotide processing" and "response").

3. Proteins enriched in COSMIC non-driver variants at protein-protein interaction sites.

4. Statistics for comparisons of structural network features between datasets.

5. Numbers of proteins and missense variants which underlie correlations between proteomic/transcriptomic features and variant enrichment.

6. Pairwise Spearman correlations between all studied proteomics and transcriptomics features.

[5] William McLaren, Laurent Gil, Sarah E Hunt, Harpreet Singh Riat, Graham R S Ritchie, Anja Thormann, Paul Flicek, and Fiona Cunningham. The ensembl variant effect predictor. *Genome biology*, 17(1):122, 06 2016.

[6] Sun Sook Chung, Anna Laddach, N. Shaun Bevan Thomas, and Franca Fraternali. Short loop motif profiling of protein interaction networks in acute myeloid leukaemia. *bioRxiv*, 2018.

[17] Bert Vogelstein, Nickolas Papadopoulos, Victor E Velculescu, Shibin Zhou, Luis A Diaz, and Kenneth W Kinzler. Cancer genome landscapes. *Science (New York, N.Y.)*, 339(6127):1546–58, Mar 2013.

[18] Steffen Durinck, Yves Moreau, Arek Kasprzyk, Sean Davis, Bart De Moor, Alvis Brazma, and Wolfgang Huber. Biomart and bioconductor: a powerful link between biological databases and microarray data analysis. *Bioinformatics*, 21:3439–3440, 2005.

[19] Steffen Durinck, Paul T. Spellman, Ewan Birney, and Wolfgang Huber. Mapping identifiers for the integration of genomic datasets with the r/bioconductor package biomart. *Nature Protocols*, 4:1184–1191, 2009.

[20] Juan M Vaquerizas, Sarah K Kummerfeld, Sarah A Teichmann, and Nicholas M Luscombe. A census of human transcription factors: function, expression and evolution. *Nature reviews. Genetics*, 10(4):252–63, 04 2009.

[21] Robert D Finn, Teresa K Attwood, Patricia C Babbitt, Alex Bateman, Peer Bork, Alan J Bridge, Hsin-Yu Chang, Zsuzsanna Dosztányi, Sara El-Gebali, Matthew Fraser, Julian Gough, David Haft, Gemma L Holliday, Hongzhan Huang, Xiaosong Huang, Ivica Letunic, Rodrigo Lopez, Shennan Lu, Aron Marchler-Bauer, Huaiyu Mi, Jaina Mistry, Darren A Natale, Marco Necci, Gift Nuka, Christine A Orengo, Youngmi Park, Sebastien Pesseat, Damiano Piovesan, Simon C Potter, Neil D Rawlings, Nicole Redaschi, Lorna Richardson, Catherine Rivoire, Amaia Sangrador-Vegas, Christian Sigrist, Ian Sillitoe, Ben Smithers, Silvano Squizzato, Granger Sutton, Narmada Thanki, Paul D Thomas, Silvio C E Tosatto, Cathy H Wu, Ioannis Xenarios, Lai-Su Yeh, Siew-Yit Young, and Alex L Mitchell. Interpro in 2017-beyond protein family and domain annotations. *Nucleic acids research*, 45(D1):D190–D199, Jan 2017.

[22] David S Wishart, Yannick D Feunang, An C Guo, Elvis J Lo, Ana Marcu, Jason R Grant, Tanvir Sajed, Daniel Johnson, Carin Li, Zinat Sayeeda,

Nazanin Assempour, Ithayavani Iynkkaran, Yifeng Liu, Adam Maciejewski, Nicola Gale, Alex Wilson, Lucy Chin, Ryan Cummings, Diana Le, Allison Pon, Craig Knox, and Michael Wilson. Drugbank 5.0: a major update to the drugbank database for 2018. *Nucleic acids research*, 46(D1):D1074–D1082, Jan 2018.

[23] Mingcong Wang, Christina J Herrmann, Milan Simonovic, Damian Szklarczyk, and Christian von Mering. Version 4.0 of paxdb: Protein abundance data, integrated across model organisms, tissues, and cell-lines. *Proteomics*, 15(18):3163–8, Sep 2015.

[24] Holger Franken, Toby Mathieson, Dorothee Childs, Gavain M A Sweetman, Thilo Werner, Ina Tögel, Carola Doce, Stephan Gade, Marcus Bantscheff, Gerard Drewes, Friedrich B M Reinhard, Wolfgang Huber, and Mikhail M Savitski. Thermal proteome profiling for unbiased identification of direct and indirect drug targets using multiplexed quantitative mass spectrometry. *Nature protocols*, 10(10):1567–93, Oct 2015.

[25] Toby Mathieson, Holger Franken, Jan Kosinski, Nils Kurzawa, Nico Zinn, Gavain Sweetman, Daniel Poeckel, Vikram S Ratnu, Maike Schramm, Isabelle Becher, Michael Steidel, Kyung-Min Noh, Giovanna Bergamini, Martin Beck, Marcus Bantscheff, and Mikhail M Savitski. Systematic analysis of protein turnover in primary cells. *Nature communications*, 9(1):689, 02 2018.

[26] GTEx Consortium. The genotype-tissue expression (gtex) project. *Nature genetics*, 45(6):580–5, Jun 2013.

[27] Helen Berman, Kim Henrick, and Haruki Nakamura. Announcing the worldwide protein data bank. *Nature structural biology*, 10(12):980, Dec 2003.

[28] S F Altschul, T L Madden, A A Schäffer, J Zhang, Z Zhang, W Miller, and D J Lipman. Gapped blast and psi-blast: a new generation of protein database search programs. *Nucleic acids research*, 25(17):3389–402, Sep 1997.

[29] Robert D Finn, Penelope Coggill, Ruth Y Eberhardt, Sean R Eddy, Jaina Mistry, Alex L Mitchell, Simon C Potter, Marco Punta, Matloob Qureshi, Amaia Sangrador-Vegas, Gustavo A Salazar, John Tate, and Alex Bateman. The pfam protein families database: towards a more sustainable future. *Nucleic acids research*, 44(D1):D279–85, Jan 2016.

[30] Robert D Finn, Jody Clements, and Sean R Eddy. Hmmer web server: interactive sequence similarity searching. *Nucleic acids research*, 39(Web Server issue):W29–37, Jul 2011.

[31] C Notredame, D G Higgins, and J Heringa. T-coffee: A novel method for fast and accurate multiple sequence alignment. *Journal of molecular biology*, 302(1):205–17, Sep 2000.

[32] Luigi Cavallo, Jens Kleinjung, and Franca Fraternali. Pops: A fast algorithm for solvent accessible surface areas at atomic and residue level. *Nucleic acids research*, 31(13):3364–6, Jul 2003.

[33] E W Myers and W Miller. Optimal alignments in linear space. *Computer applications in the biosciences : CABIOS*, 4(1):11–7, Mar 1988.

[34] Li C Xue, Drena Dobbs, and Vasant Honavar. Homppi: a class of sequence homology based protein-protein interface prediction methods. *BMC bioinformatics*, 12:244, Jun 2011.

[35] Ahmet Bakan, Lidio M Meireles, and Ivet Bahar. Prody: protein dynamics inferred from theory and experiments. *Bioinformatics (Oxford, England)*, 27(11):1575–7, Jun 2011.

[36] Aric Hagberg, Pieter Swart, and Daniel S Chult. Exploring network structure, dynamics, and function using networkx. Technical report, Los Alamos National Lab.(LANL), Los Alamos, NM (United States), 2008.

[37] Alexey Sergushichev. An algorithm for fast preranked gene set enrichment analysis using cumulative statistic calculation. *bioRxiv*, 2016.

[38] Ian Sillitoe, Tony E Lewis, Alison Cuff, Sayoni Das, Paul Ashford, Natalie L Dawson, Nicholas Furnham, Roman A Laskowski, David Lee, Jonathan G Lees, Sonja Lehtinen, Romain A Studer, Janet Thornton, and Christine A Orengo. Cath: comprehensive structural and functional annotations for genome sequences. *Nucleic acids research*, 43(Database issue):D376–81, Jan 2015.

[39] Sameer Velankar, José M Dana, Julius Jacobsen, Glen van Ginkel, Paul J Gane, Jie Luo, Thomas J Oldfield, Claire O'Donovan, Maria-Jesus Martin, and Gerard J Kleywegt. Sifts: Structure integration with function, taxonomy and sequences resource. *Nucleic acids research*, 41(Database issue):D483–9, Jan 2013.

[40] AB MySQL. *Mysql 5.1 reference manual*, 2008.

[41] Django. http://djangoproject.com.

[42] Nicholas M Luscombe and Janet M Thornton. Protein-dna interactions: amino acid conservation and the effects of mutations on binding specificity. *Journal of molecular biology*, 320(5):991–1009, Jul 2002.

[43] Remo Rohs, Xiangshu Jin, Sean M West, Rohit Joshi, Barry Honig, and Richard S Mann. Origins of specificity in protein-dna recognition. *Annual review of biochemistry*, 79:233–69, 2010.

[44] Bohdan Schneider, Jirí Cernỳ, Daniel Svozil, Petr Cech, Jean-Christophe Gelly, and Alexandre G de Brevern. Bioinformatic analysis of the protein/dna interface. *Nucleic acids research*, 42(5):3381–94, Mar 2014.
